# Inferring fluctuating interaction probabilities in ecological networks across environmental change

**DOI:** 10.64898/2026.05.06.723199

**Authors:** Phuong L. Nguyen, Luis J. Gilarranz, Rudolf P. Rohr

## Abstract

Knowledge of species’ interactions unlocks our understanding of how ecological communities respond to climate change or habitat loss, explaining their resilience and robustness. Such knowledge requires inferring the presence, sign, and *per capita* strength of species interactions, as well as species’ intrinsic growth rates. While various studies have attempted to infer these parameters in isolation, none has successfully inferred them simultaneously. Here, we solve this grand challenge using an integrative approach combining ecological mechanistic models and statistical inference to simultaneously infer these parameters across time, capturing environmental change. We validate our approach on synthetic data in constant and changing environments with both abrupt and gradual changes. Applying our method to a manipulated microcosm experiment demonstrates its ability to infer environmental change in empirical data. Finally, its application to a mesocosm experiment recovers the expectations from biological knowledge and unveils network rewiring due to unplanned environmental change. Our approach goes one step further in bridging the gap between mechanistic models and empirical ecology. It advances the understanding of ecological networks and their dynamics, thereby helping to validate existing hypotheses, spark new theories, and guide ecological management and conservation.

## 1 Introduction

Ecological communities are, at a fundamental level, described by networks of species interactions [1, 2]. Decades of research have shown that network structure largely governs community dynamics and species coexistence [3, 4]. Recording the presence of interactions allows the quantification of properties like network connectance, as well as ubiquitous patterns – such as modularity in antagonistic networks or nestedness in mutualistic networks. These network properties strongly influence community stability and biodiversity [5, 2, 6, 7, 8]. Subsequently, interaction strengths have also been shown to modulate community stability and persistence [9, 10, 11]. Moreover, the turnover of species interactions across seasonal fluctuations determines community persistence amid environmental change [12]. This fruitful research program on ecological networks has shown that, to understand the dynamical characteristics of ecological communities and predict their future in yet-to-be-observed scenarios, knowledge of the structure of species interactions, their *per capita* interaction strengths, and species’ intrinsic growth rates is necessary [13, 4, 14, 15].

Retrieving ecological networks from empirical data, however, poses a great challenge. Direct observations of interactions – such as plant-pollinator interactions based on flower visitation or food webs based on gut content analysis – are limited in their choice of biological systems and the interaction mechanism [16, 17]. They are often static (interactions are rarely tracked over time), and the dynamical effect of the observed interactions is unknown. In the absence of direct observations, ecologists have tried to infer species interactions based on species co-occurrence [18]. However, species co-occurrence may not imply interactions [19]. In parallel, the availability of community-wide time series of population abundances has allowed the use of S-maps [20] – an approach based on local linear regressions on the total species abundance – to infer the time-varying Jacobian matrix [21, 22]. However, the Jacobian matrix is devised to assess local stability – return to an equilibrium point after a sufficiently small perturbation – and does not capture the fundamental *per capita* interactions between species, which are among the key parameters in ecology [23, 24, 25, 26, 27]. Moreover, the reconstruction of ecological networks fundamentally entails inferring the presence of interactions. Regularization techniques have then been used to capture interaction presence [28]. However, regularization is known to bias parameter estimates in both their magnitude and sign [29], thereby biasing the elements of the Jacobian matrix. While the Lotka-Volterra map (LV-map) [26] – an approach based on local linear regressions on species’ *per capita* growth – can obtain the unbiased *per capita* interaction strengths, it cannot obtain the presence of interactions. Therefore, to this day, there is no method that can simultaneously infer all three ingredients that are needed to understand community dynamics: the presence of interactions, the unbiased interaction strengths, and the intrinsic growth rates.

Here, we present the MA-LVmap, a method designed to infer species’ pairwise interaction probabilities, unbiased *per capita* interaction strengths, and intrinsic growth rates from time series of species abundances, along with their potential variation across time (figure 1). To do so, we combine the LV-map [26] with model averaging techniques [30, 31, 29]. The LV-map, based on well-established ecological mechanisms, provides estimates of intrinsic growth rates and *per capita* interactions that vary over time and therefore along environmental variation, while model averaging techniques assign a probability to each possible interaction network, from which pairwise probabilities can be extracted. As a result, the MA-LVmap can quantify how known environmental change affects the inferred parameters through their variation across time. Because unanticipated environmental shifts leave a similar signature – temporal variation in these same parameters – the method can also detect such changes even when they are not known in advance.

**Fig 1.**
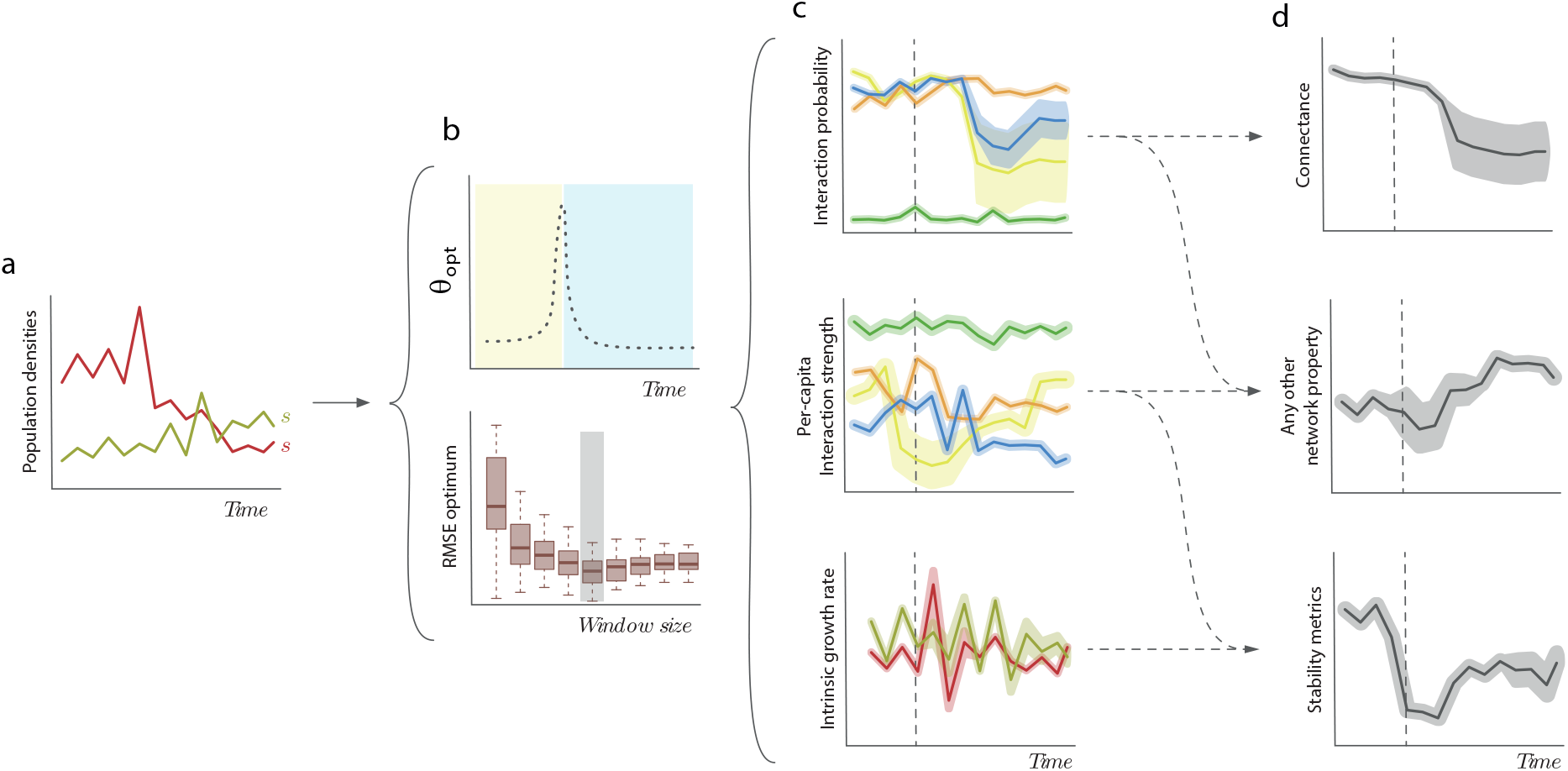
Workflow for the MA-LVmap approach. Using only time series of population densities (**a**), the MA-LVmap is capable of obtaining the best window size (indicated by the gray-shaded boxplot) and the time-varying localization parameter (*θ_opt_*(*t*)), both aimed at capturing environmental variation (**b**) as well as time-varying interaction probabilities (*p*^*_ij_*(*t*)), *per capita* interaction strengths (*ε*^*_ij_*(*t*)), and intrinsic growth rates (*r*^*_i_*(*t*)) (**c**). The MA-LVmap allows us to track any network property and the uncertainty around its estimation, as well as stability metrics (**d**) over time.

We validate the MA-LVmap at different levels. First, using synthetic and semi-synthetic data, we show that the MA-LVmap properly infers ecological networks, including network structures, intrinsic growth rates, and *per capita* interaction strengths. More importantly, our method can infer the effects of environmental change on interactions and intrinsic growth rates over time regardless of the speed of the environmental change. Then, as a proof of concept, we apply our method to time series of a microcosm experiment controlled by two treatments: constant versus changing environment. We show that our method correctly distinguishes the two treatments and infers reasonable ecological parameters. Finally, we apply our method to time series of a long-term mesocosm experiment involving a plankton community isolated from the Baltic Sea, where there is an unexpected change in nutrient concentration and photoperiod [32]. We demonstrate that the MA-LVmap detects the effect of this unexpected change in environmental conditions (nutrient concentration and photoperiod) on both the structure and interaction strengths of the ecological network. These results show that our method enables a mechanistic understanding of both ecological networks and interaction strengths, offering insightful perspectives on community structures, dynamics, and the mechanisms maintaining biodiversity.

## 2 Methods

### The LV-map approach [26]

It is grounded in the ecological principles and mechanisms of multi-species Lotka-Volterra (time-continuous) or Ricker (time-discrete) population dynamics [33]. Specifically, the *per capita* rate of change for each species depends on (i) its intrinsic growth rate *r_i_*(*t*) and (ii) *per capita* interactions with the other species *ε_ij_*(*t*). By definition, the intrinsic growth rate is the *per capita* rate of change when all species are rare, and the *per capita* interaction is the *per capita* effect of one species on the *per capita* rate of change of another species [33]. Mathematically, this is formulated as the well-known Lotka-Volterra or Ricker models,

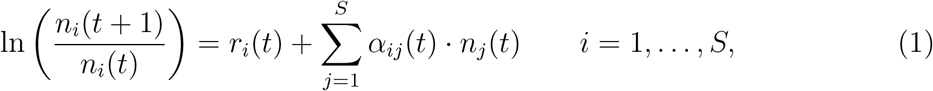

where *n_i_* is the density of species *i* and *S* is the total number of species. The LV-map correctly infers the intrinsic growth rate *r_i_*(*t*) and the *per capita* interaction strength *ε_ij_*(*t*), along with their potential variation over time. From a technical perspective, this is done using local linear regressions and weighting kernels (see Supplementary Information S1 and Nguyen et al. [26]).

### Combination of LV-map and model averaging to obtain interaction probabilities

The *per capita* interactions inferred by the LV-map are never exactly zero, since in a linear regression – even in the case where the true slope is zero – the inferred value will always deviate from zero. Thus, inferring the presence of an interaction means identifying the true non-zero interaction strengths, which, from a statistical perspective, can be seen as finding the “statistically significant” interaction strengths.

Many approaches are available to determine whether or not a parameter is statistically significant [30, 34, 29]. For example, the significance can be based on whether or not the confidence interval includes zero. Information criteria such as AIC and BIC, with backward and forward selection, can also be used, or a log-likelihood ratio test for each possible interaction is another alternative approach. However, the principles of determining significant parameters are often rooted in selecting a single “best” model, leading to near-arbitrary decisions. Take the example of the model selection approach using given information criteria: if the two best models have extremely similar criterion values but differ in a single interaction, what is the conclusion for this interaction? Importantly, how do we justify a binary choice for the presence of the interaction given that the binary choice does not allow us to consider the inference uncertainty?

To address these challenges, we propose inferring interactions as probabilities of presence, with their confidence intervals, using model averaging [30, 31, 29]. It works as follows. For each species *i*, it considers all possible interaction combinations *k* with every other species *j*: *c_ijk_* = 1 if species *j* interacts with species *i* in combination *k*. Note that, for a given species *i*, there are 2*^S→^*^1^ combinations as we consider intraspecific interaction to be always present. This choice is justified because, in model averaging, one should consider a set of reasonable biological models [30, 31, 35], and it is ecologically reasonable to assume that intraspecific interactions are always present [36, 33]. For each combination of interactions *k*, we perform local linear regression between the *per capita* growth of species *i* and the abundances of the other species ln(*n_i_*(*t* + 1)*/n_i_*(*t*)) = *r_i_*(*t*)+ *c_ijk_ · ε_ij_*(*t*) *· n_j_*(*t*). This results in a set of BIC values (BIC(*t*)*_ik_*). The posterior probability of each of these combinations – a potential set of interactions – is given by its Bayesian weight,

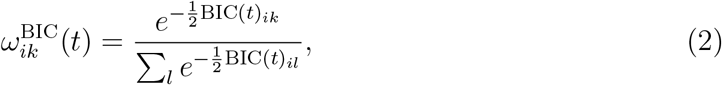

from which the pairwise interaction probabilities are estimated as

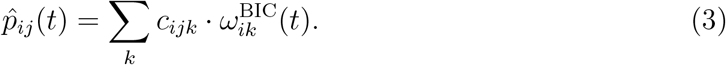

The term *c_ijk_*= 1 if species *j* interacts with species *i* in the combination *k* and 0 otherwise.

Statistically speaking, *p*^*_ij_*(*t*) is the probability of inclusion of the parameter *ε_ij_* in the model, that is, the probability of an interaction from *j* to *i*. The method is presented here with BIC weights, but AIC weights can also be chosen. The choice between the AIC and BIC weights is a matter of the aims of the study. The BIC puts a heavier penalty on complex models and asymptotically tends to select the true model if it is in the set of proposed models [30, 31, 35]. On the other hand, the AIC is known to select the best model for prediction. The BIC is therefore inference-focused because it is more consistent and better at identifying true interactions, while the AIC is more prediction-focused and better at finding predictors for the *per capita* growth rates. More specifically, we expect the BIC weight to perform better under low connectance and strong interaction strengths, whereas we expect the AIC weight – which penalizes additional parameters less heavily and is therefore more permissive of weak interactions – to perform better under high connectance and weak interaction strengths.

Model averaging allows us to obtain all pairwise interspecific interaction probabilities. In combination with the LV-map, we obtain these probabilities as a function of time, which enables the detection of network rewiring. Rewiring events are shown as large shifts in interaction probabilities, from low to high probabilities or vice versa.

### Inference across environmental change

In the LV-map approach, a weighting kernel was used to allow variation of the intrinsic growth rates and *per capita* interaction strengths across time. This is essentially a local linear regression where the parameters are estimated at each time point along the time series. The weighting kernel is defined as follows. For a focal time point *t*, the weight of each other time point *l* is defined as

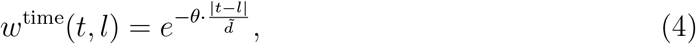

where *d̃* denotes the average distance between all pairs of time points. The parameter *θ* defines the inverse of the kernel width, which we therefore call the localization parameter. A large *θ* implies a highly local linear regression, where only the time points closest to the focal point *t* are accounted for. In contrast, a small *θ* indicates that more distant time points are considered. In the original LV-map, expanding-window cross-validation was used on the full dataset to estimate a single best value of *θ* [37, 26].

However, the best localization parameter *θ* may change over time due to changes in the environment. Therefore, a single value of *θ* may not adequately capture changes in environmental conditions, resulting in inaccurate inference of interaction probabilities, *per capita* interaction strengths, and intrinsic growth rates. To obtain a time-varying *θ*, we calculate the optimum *θ* and its root mean square error (RMSE) within a moving window [37]. This window has a fixed size (*m*), defined as the proportion of data points relative to the full time series, and moves along the time series. Consequently, for a chosen window size (*m*), we obtain a time series of optimum *θ* values along with their corresponding RMSE values. The window size itself is chosen based on the minimum median value of the RMSE. Then, the corresponding time series of optimum *θ* values is generated. The impact of environmental change can be seen at different levels. The first-level signal of time-varying parameters is a moving window size that is not, or almost not, the whole time series. The second-level signal is an abrupt increase in the optimum *θ*, indicating a sudden environmental shift. The third-level signal is the time variation in the inferred interaction probabilities, strengths, and/or intrinsic growth rates.

## 3 Results

We demonstrate the effectiveness of the MA-LVmap in detecting interactions and their strengths and signs. We first assess the performance of the MA-LVmap in inferring synthetic networks in a constant environment, i.e., with fixed parameters. We then focus on results with changing environments by simulating systems with various speeds of change. Finally, as a proof of concept, we apply the MA-LVmap to empirical datasets of microbial communities. Details of the methods, as well as supplementary results, are in the Supplementary Information.

### (a) Validation in a constant environment

We simulated synthetic networks, for which we controlled the number of species *S*, connectance *c*, and *σ* – the *per capita* interspecific interaction strengths “relative” to intraspecific ones. The parameter *σ* corresponds to the standard deviation of the sampled interspecific interactions. Thus, a large *σ* corresponds to a distribution with strong interaction strengths, while a small *σ* corresponds to weak interaction strengths. The MALVmap proves effective in inferring species interactions (figure S2–S12). The MA-LVmap also performs well in inferring network structure – modularity and nestedness – for semi-synthetic networks (figure S14, S15). While inference ability is not affected by community size, it varies depending on the level of interaction strength (*σ*) and connectance (*c*).

An important feature of the MA-LVmap is that it allows us to infer high-probability weak interactions because strong interactions are not always correlated with high probabilities and vice versa (figure S13). In nature, weak interactions have been shown to be of great importance for community dynamics [9, 38, 11]. This contrasts with other techniques like regularization, which would set weak interactions to zero, biasing the quantification of network properties. More importantly, and related to the aforementioned fact, the MA-LVmap excels in the inference of the correct interaction sign, outperforming the regularization techniques by a factor of two (figure S17). Moreover, like the LV-map, the MA-LVmap correctly infers the intrinsic growth rates and *per capita* interaction strengths (figure S16, S17).

Although model averaging can be performed using AIC or BIC weights, and the BIC weight is generally preferred for inference, the choice may depend on the nature of the network. While the BIC weight is highly efficient for networks with low connectance (*c* = 0.05) and strong interaction strengths (*σ* = 0.1) (figure S2a, S3a, c, e), the AIC weight is more suitable for networks with high connectance (*c* = 0.5) and weak interactions (figure S2j, S7b, d, f). Because natural interaction networks often combine low connectance with weak interaction strengths, neither criterion is uniformly optimal, and the choice between AIC and BIC should be guided by the specific characteristics of the system under study. Reassuringly, this choice affects only the ability to detect interactions: both criteria yield similar estimates of the intrinsic growth rates and *per capita* interaction strengths (figure S16a–h, S17).

### (b) Validation in changing environments

Next, to evaluate the ability of the MA-LVmap to infer ecological networks across environmental change, we simulate a community of two prey (*R*_1_, *R*_2_) that share a common predator (*C*). To include environmental change in the model, we consider the scenario where a parasite specialized on prey *R*_1_ is introduced, causing a reduction in its intrinsic growth rate. We assume that this results in a shift in the predator’s preference toward the other prey *R*_2_, so that both *ε_CR_*_1_ and *ε_R_*_1*C*_ tend to zero. This leads to a change in the expected total number of interactions from 9 to 7. Moreover, we control the speed of the environmental change, from no change at all (control case) to slow and intermediate change speeds and finally an extremely abrupt change; see panels a–d in figure 2.

**Fig 2.**
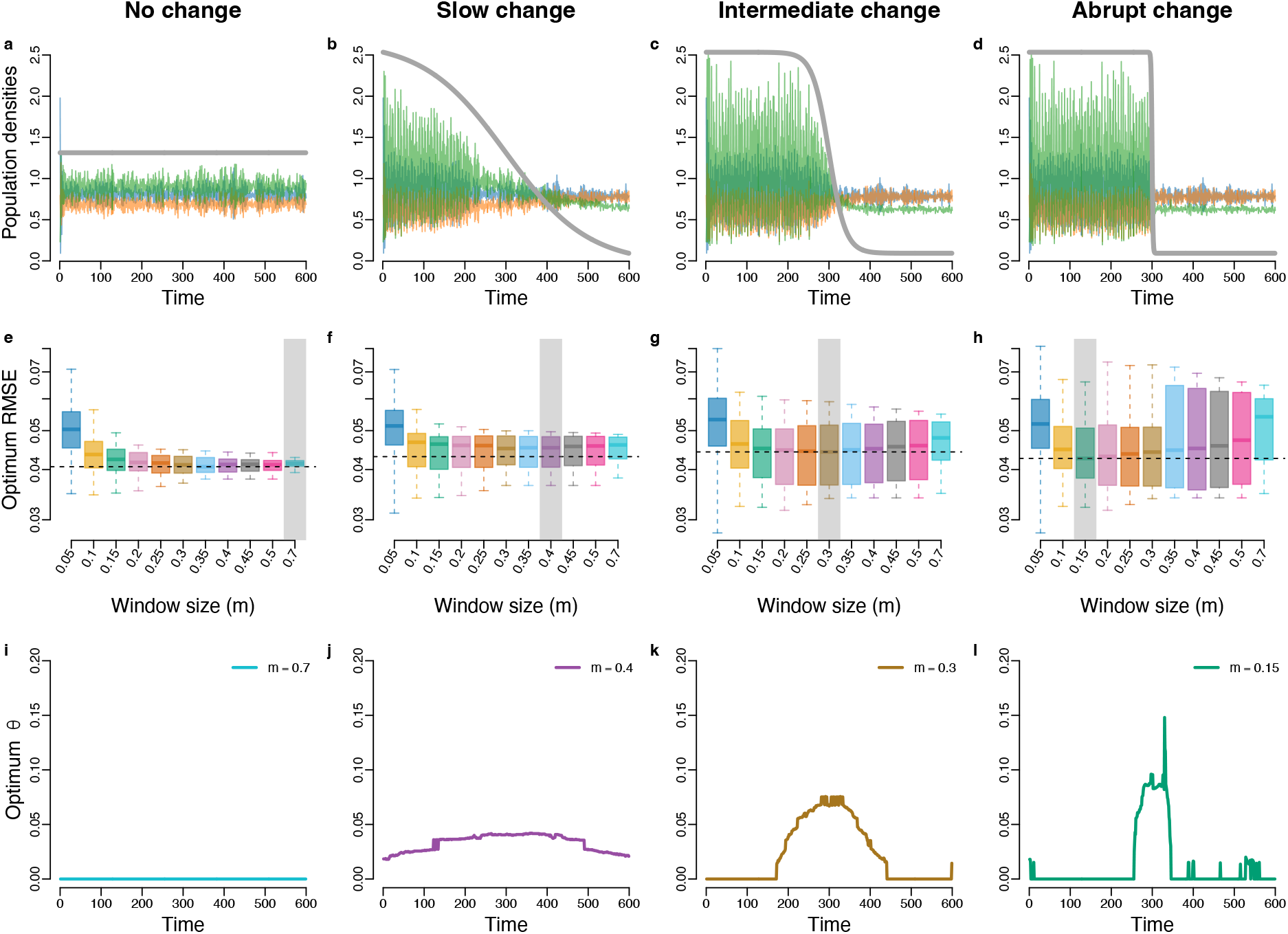
Inference of the synthetic predator-prey network across environmental change. **(a–d)** Population densities of the predator-prey community under different scenarios of environmental change. Gray lines illustrate the speed of environmental change. **(e–h)** Optimum RMSE corresponding to different window sizes. Gray rectangles indicate the best window size. **(i–l)** Time series of optimum *θ* values corresponding to the best window size.

As a baseline, the first column in figure 2 presents the results in a constant environment. Panel e in figure 2 shows the RMSE as a function of the window size. As expected, larger window sizes correspond to lower RMSE medians and smaller variation. In this scenario, the best window size is the largest one (*m* = 0.7). In turn, panel i provides the optimum localization parameter (*θ*) as a function of time for the best window size. In a constant environment, *θ* is constant and equal to 0.

In scenarios with abrupt environmental changes, panels g and h indicate that the more abrupt the change is, the smaller the optimum window size (figure 2f–h). Moreover, the optimum *θ* deviates from zero (figure 2j–l). In particular, panels k and l show that the more abrupt the change is, the narrower the peak in the localization parameter *θ* centered around the change will be. This behavior is expected because indications of a change in environmental conditions are most perceptible when the change is abrupt – as reflected in a smaller optimum window size and a spike pattern in the optimum *θ* time series. This is intuitive, as the slower the effect of environmental change, the longer the transition phase of the population dynamics, making it harder to detect the dynamical changes.

When the environmental change is very gradual (figure 2b, f, j), the optimum window size is less obvious, and a range of optimum window sizes may be chosen as their RMSE values are similar (figure 2f). In such a case, additional analysis of the interaction probabilities and strengths may be required to draw proper conclusions.

Figure 3 shows the inferred interaction probabilities, interaction strengths, expected number of interactions, and intrinsic growth rates obtained using the best window size for each scenario of environmental change. The MA-LVmap captures the decline in connectance and correctly infers the true connectance value as well as the interaction probabilities at all speeds of environmental change (figure 3b–d, f–h, j–l). The method also produces reliable estimates of intrinsic growth rates and *per capita* interaction strengths (figure 3, S18–S21).

**Fig 3.**
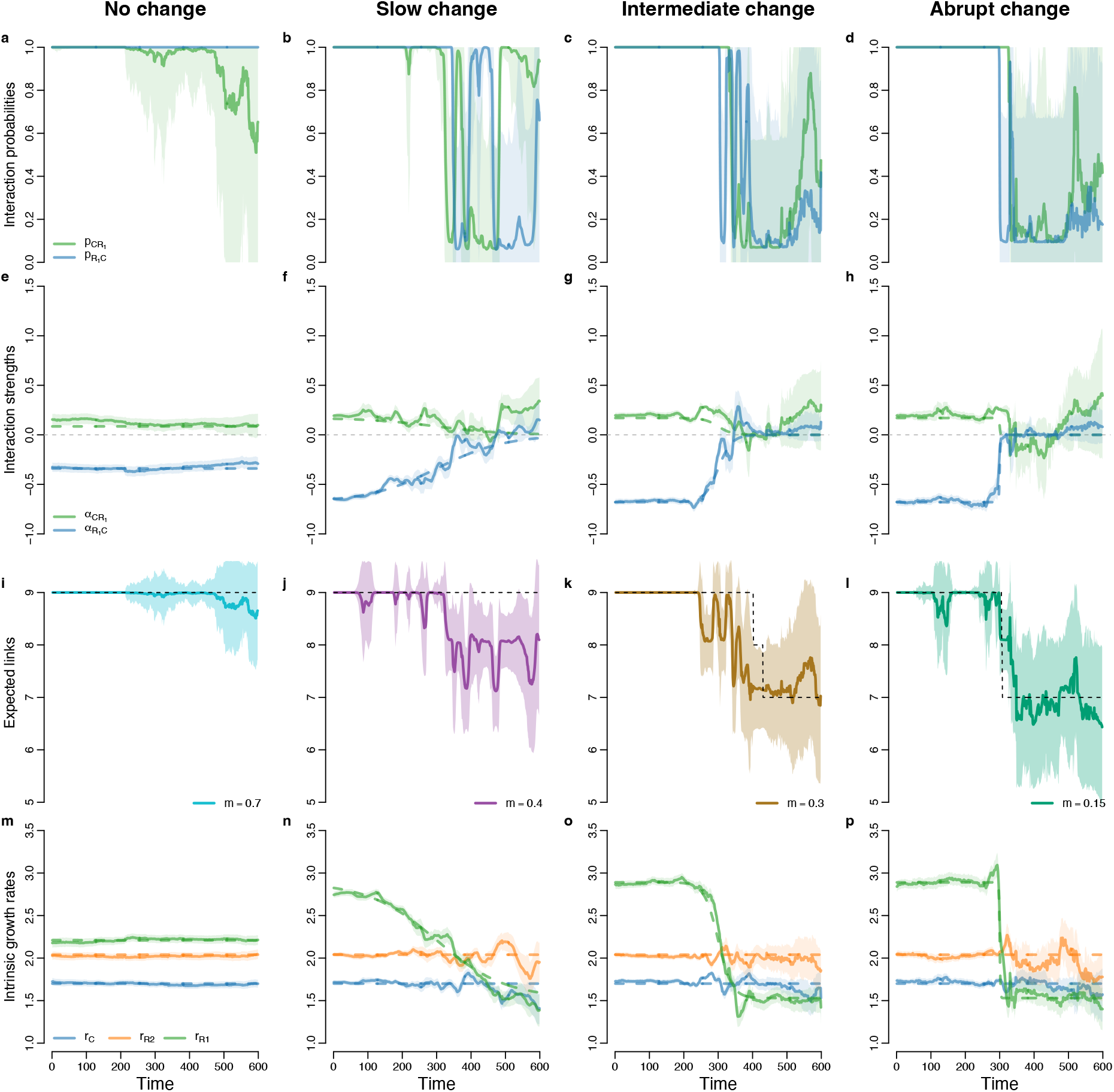
Parameter inference for the synthetic predator-prey system. **(a–d)** Interaction probabilities across time for different scenarios of environmental change. **(e–h)** Interaction strengths between the common predator (*C*) and prey 1 (*R*_1_) across time. **(i–l)** Expected number of links across time using the best window size. **(m–p)** Inference of intrinsic growth rates for both prey (*R*_1_, *R*_2_) and the common predator (*C*).

### (c) Proof of concept 1: Microcosm experiment

As a proof of concept, we apply the MA-LVmap to a microcosm experiment in which environmental conditions are systematically manipulated [39]. The experiment involves a variety of communities of phytoplankton and zooplankton that are cultured in 30 bottles divided into two treatments: constant light environment (15 bottles) and gradually decreasing light intensity (15 bottles). Details of the experiment can be found in Supplementary Information S4.1.

As could be expected, the results for optimum RMSE as a function of window size and the time series of optimum *θ* differ substantially between constant and decreasing light treatments. For the constant light treatment, the optimum window size is generally larger than for the decreasing light treatment (figure 4c, d, S22). Note that, because the main purpose is to compare the two treatments, we present results using only the BIC weight.

**Fig 4.**
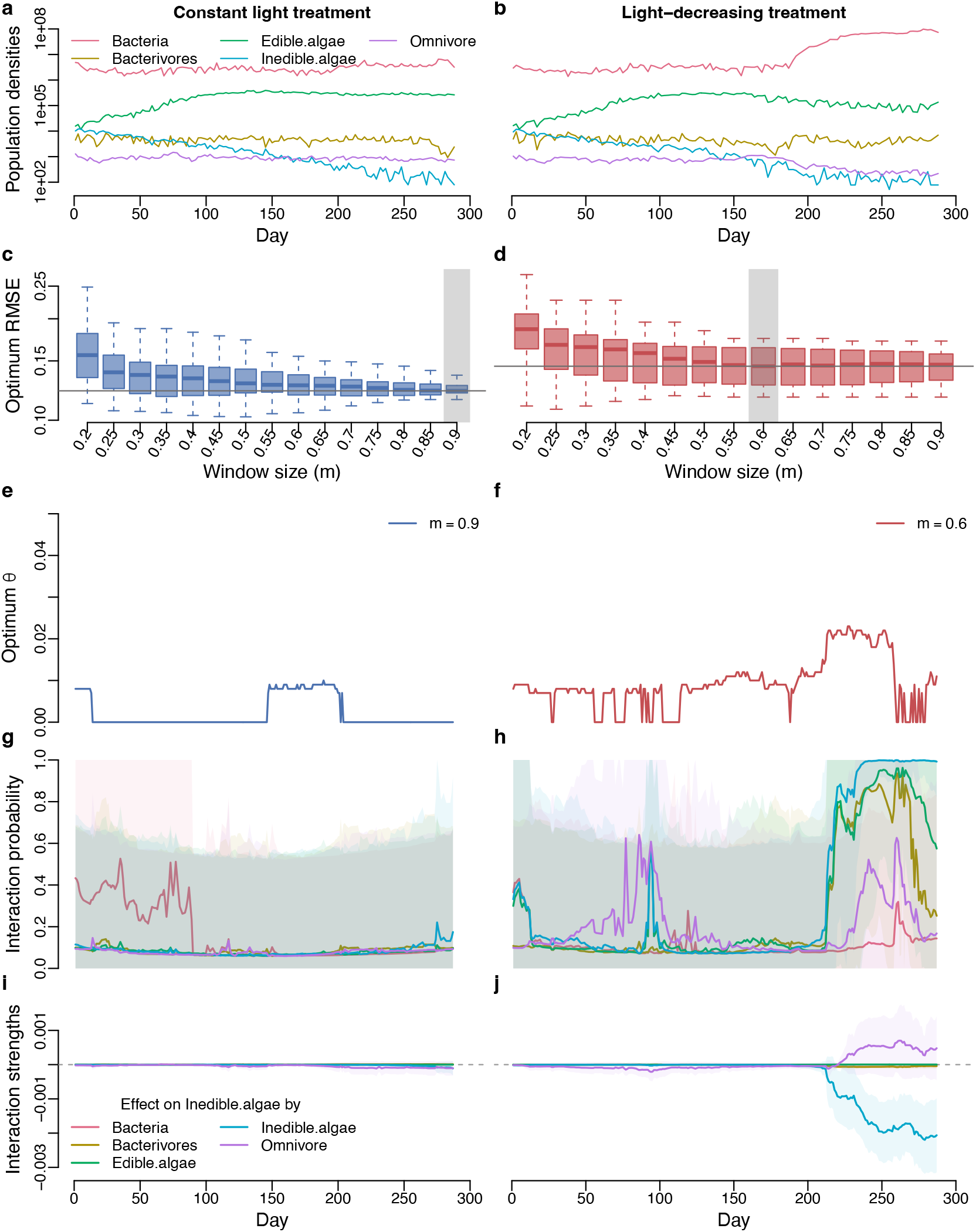
Comparison between the constant light treatment (bottle 21) and the decreasing light treatment (bottle 10). These two bottles share the same species composition with 10 species categorized into five functional groups. **(a, b)** Population densities. **(c, d)** Optimum RMSE corresponding to different window sizes. The gray rectangle indicates the best window size. **(e, f)** Time series of optimum *θ* for the best window size. **(g, h)** Changes in the interaction probabilities between inedible algae and other functional groups. **(i, j)** Changes in the *per capita* interaction strengths between inedible algae and other functional groups in bottles 21 and 10, respectively. Other interactions are shown in figure S23.

When comparing the optimum *θ* time series, we observe that in the decreasing light treatment, there is an increase in the optimum *θ* around day 200. However, unexpectedly, the optimum *θ* values for the constant light treatment slightly deviate from zero, which might be due to the fact that while light was constant, other abiotic factors were not controlled.

When comparing interaction probabilities and interaction strengths, we see that for the decreasing light treatment, the peak of optimum *θ* around day 200 corresponds to an increase in the intraspecific interaction strengths of inedible algae and interaction strengths between this group and the omnivore, and a sharp increase in interaction probabilities (figure 4h, j). For the constant light treatment, there is almost no change in the inferred interaction strengths and interaction probabilities (figure 4g, i, S23).

These are the general differences between the constant and decreasing light treatments, but there are exceptions. Community responses to the decreasing light treatment differ greatly. Some communities experience an increase in connectance, some a decrease, and others no change (figure S24–S51). This suggests that community composition plays a key role in how a community responds to environmental change. Nevertheless, the MA-LVmap is effective at quantifying the consequences of these changes for the interactions.

### (d) Proof of concept 2: Mesocosm experiment

To further showcase the effectiveness of the MA-LVmap, we apply our method to a mesocosm experiment [40, 32]. The studied community includes five functional groups: calanoid copepods, rotifers, nanoflagellates, picocyanobacteria, and bacteria (figure 5a). Details of the data are in Supplementary Information S4.2.

**Fig 5.**
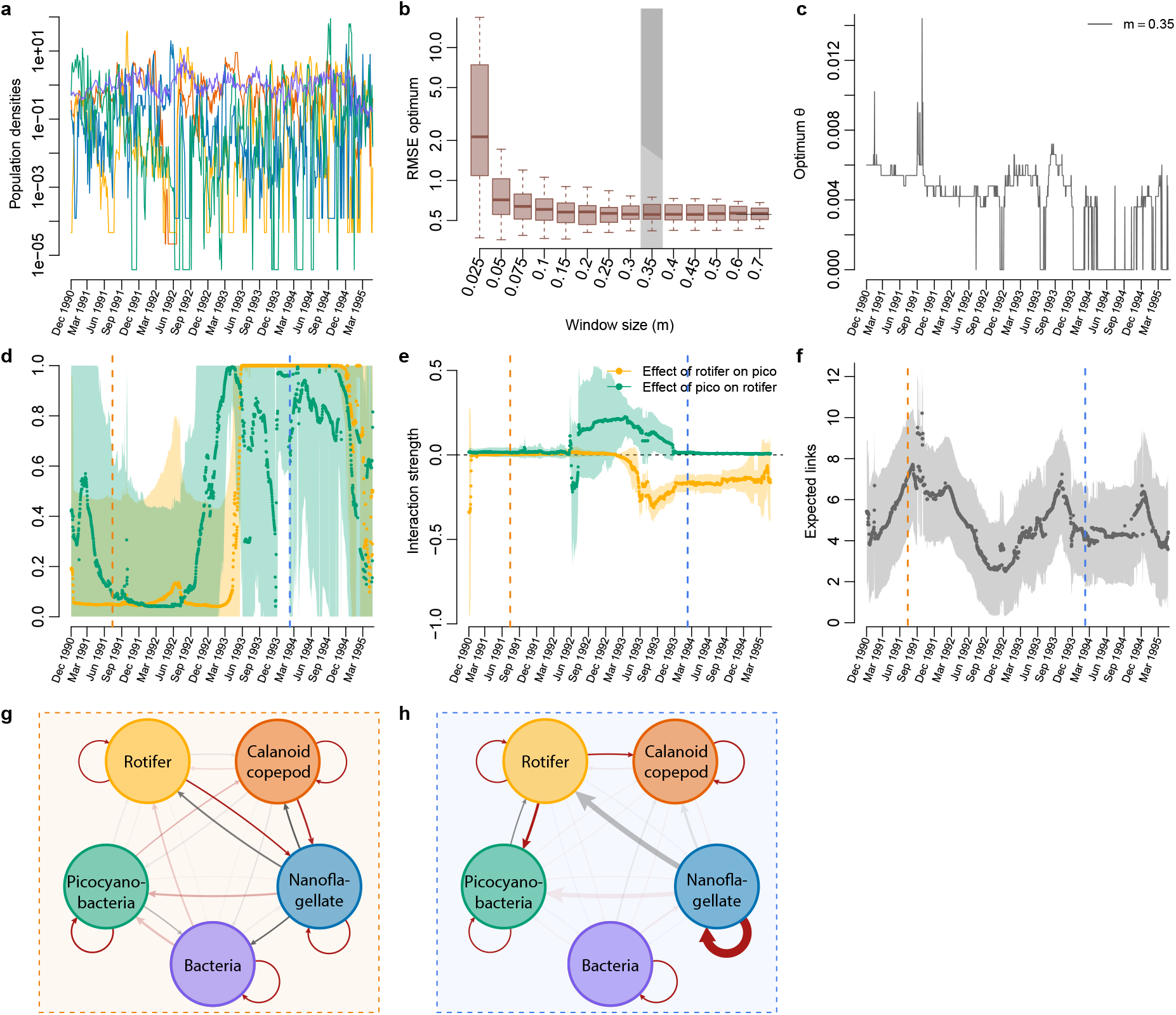
Inference of the mesocosm experiment. **(a)** Population densities. **(b)** Optimum RMSE with respect to different window sizes. **(c)**Time series of optimum *θ*. **(d)** Inferred interaction probabilities between rotifer and picocyanobacteria. **(e)** Inferred *per capita* interaction strengths between rotifer and picocyanobacteria. **(f)** Inferred expected number of links across time. **(g)** The inferred community network on 14 July 1991 (dashed vertical orange line). **(h)** The inferred community network on 8 February 1994 (dashed vertical blue line). Link thickness indicates interaction strength. Link color indicates whether the interaction is positive (gray) or negative (red), and link transparency indicates interaction probability. We present the inferred networks using the BIC weight. Results using the AIC weight can be found in the Supplementary Figures. The expected number of links inferred using the AIC weight is slightly larger than that inferred using the BIC weight, but the main interactions remain (figure S61).

Both the optimum window size and the optimum *θ* time series (figure 5b, c) suggest a gradual environmental change. This corresponds to an unplanned change in photoperiod from 16 to 12 hours around April 1991, as well as a gradual reduction in *NO*_2_ and soluble reactive phosphorus (figure S53).

We observe a clear change in interaction probabilities and strengths. The initial increase in the expected links – from four to seven links – was followed by a reduction in the expected number of links synchronous with the change in photoperiod. The rewiring event includes changes in several pairwise interactions. Notably, before the decline in nutrients and photoperiod, rotifers and calanoid copepods preyed mainly on nanoflagellates. After the decline in nutrients and photoperiod, rotifers became specialized on picocyanobacteria. The probability of interaction between rotifers and nanoflagellates became low despite a stronger interaction strength than before the environmental change. This result is in line with the conclusion of Beninc’a et al. [32], who analyzed only data after the disturbance of the photoperiod and nutrients, from June 1991 until October 1997. The authors initially assumed that rotifers feed on both phytoplankton groups, but later revealed that rotifers primarily feed on picocyanobacteria while calanoid copepods mainly feed on nanoflagellates during this period [32], which matches what MA-LVmap infers.

## 4 Discussion

We have shown how the MA-LVmap simultaneously infers interaction probabilities, *per capita* interaction strengths, and intrinsic growth rates. These are the key ecological parameters required to achieve a mechanistic understanding of ecological communities [5, 33, 24]. By inferring how these parameters change over time, our method reveals the impact of environmental change, along with the corresponding changes in community structure and dynamics. By estimating the probability of species’ interactions and the variance around the expected interaction strengths, our approach acknowledges the fundamentally stochastic nature of the world, while providing analytical tools to study ecological communities within this probabilistic context [41].

Being able to infer interaction probabilities allows us to retrieve the full network architecture of ecological communities, which would be impossible without the inference of true zeros. In the literature, regularization techniques have been used to attempt to determine when two species do not interact [42, 28]. With regularization, weak interactions are often set to zero; that is, they are often considered insignificant in statistical models. However, weak interactions have been consistently shown to be fundamental to our understanding of community dynamics [9, 38, 11]. Showing that interaction probability is uncorrelated with interaction strength, see figure S13, highlights the importance of the MA-LVmap’s ability to infer highly probable weak interactions. These facts contribute to the excellent performance of the MA-LVmap in identifying the correct sign of interactions compared to other methods (figure S17), and explain why regularization-based methods largely misidentify the sign of interactions. While regularization also systematically biases the inferred interaction strengths to improve forecasting [29], our approach – focused on inference – produces unbiased values that represent biologically meaningful time-varying parameters.

The MA-LVmap approach, therefore, allows us to unveil how network structures change over time in response to environmental change. In nature, variation in network structure along environmental gradients is the norm rather than the exception [43]. Under different conditions, species can adopt different strategies, such as changing prey preference [44, 45] or exerting facilitation instead of competition with other co-occurring species [46]. It is this rewiring in network structure that enables community stability under environmental disturbances [44] or seasonality [12]. Rewiring is also the reason why species coexistence can be guaranteed as biodiversity increases within a community [45]. In general, expert knowledge covers only a specific time point or a short period of time [47]. Without inferring how communities respond to their environment over time, our ability to validate different sources of ecological knowledge might be severely hampered.

The remaining challenge is how to interpret changes in ecological parameters and network architectures over time. While the presence and speed of environmental change can be assessed by a combined examination of different signals – including the optimum RMSE across different window sizes, the time series of optimum *θ* for the chosen window size, and the changes in the interaction probabilities and interaction strengths – assigning that change to a specific driver requires further analysis. The change could be driven by abiotic factors such as light, temperature, and nutrients, but it could very well be the result of evolutionary processes, which require more in-depth analysis or dedicated experiments to unravel.

Finally, while inference using the MA-LVmap carries a well-defined ecological interpretation, as it is solidly based on mechanistic ecological models, the successful inference of all these parameters requires time series to capture *per capita* population growth. Sparse time-series data – relative to the organisms’ generation time – may result in the misinterpretation of the inferred ecological parameters. Thanks to the ongoing developments in automated population monitoring in natural systems [48, 49], we anticipate broad applicability of the MA-LVmap to various types of ecological communities [50, 51, 42]. Our approach represents a substantial leap toward bridging theoretical and empirical approaches, thereby opening the gates to testing a plethora of theories in community ecology, and creating a clear path between theory and monitoring. Such empirical tests will ultimately help us obtain a deeper understanding of the challenges faced by ecological communities in a world of ever-changing environments.

## Data and Code Availability

Once the manuscript is accepted, the code will be made public on zenodo. The raw empirical data used in this study are freely and publicly available, and the clean data that is used in this manuscript can also be found on the zenodo link 10.5281/zenodo.19571454.

