## Supplementary Information for "Inferring fluctuating interaction probabilities in ecological networks across environmental change"

**S1 Extended methods** **3**

### S1 Extended methods

#### S1.1 LV-map as a local linear regression

Inferring the *per capita* interaction strength $\alpha_{ij}(t)$ and intrinsic growth rate $r_i(t)$ is an autoregressive process of order 1 with the per capita growth $\ln(n_i(t+1)/n_i(t))$ of each species as the response variable, and the abundances $n_i(t)$ as the explanatory variables (equ. 1) [26]. Within this setting, $r_i(t)$ corresponds to the intercept and $\alpha_{ij}(t)$ to the slopes. Note that this formulation assumes additive Gaussian noise on the *per capita* growth $\ln(n_i(t+1)/n_i(t))$, which implies multiplicative Gaussian noise on the abundances $n_i(t)$. This is biologically reasonable, as change in abundances is a multiplicative process.

To infer variation with time in the parameters $r_i(t)$ and $\alpha_{ij}(t)$, we perform local linear regressions. That is, for a focal time point $t$, we set the weighting kernel $w^{\text{time}}(t,l)$ (equ. 4), which gives the weight of all other time points $l$. The parameters at time $t$ are then estimated by minimizing the weighted residual sum of squares

$$\text{RSS}_w = \sum_l \left( w^{\text{time}}(t,l) \cdot \left( \ln\left(\frac{n_i(t+1)}{n_i(t)}\right) - r_i(t) - \sum_j \alpha_{ij}(t) \cdot n_j(t) \right)\right)^2 \qquad \text{(S1)}$$

for each species $i$ as a response variable [29, 26].

#### S1.2 Time-varying weighting kernel

The localization parameter $\theta$ is estimated by cross-validation and usually assumed to be fixed (time-independent). Notably, in the original LV-map, we use the expanding-window cross-validation [37, 26]. To capture changing environments, we extended the cross-validation by the use of moving windows, allowing the variation of the localization parameter $\theta(t)$ across time. Briefly, for each time point $t$, we perform a local cross-validation within a window and estimate its corresponding optimal $\theta(t)$. The local window is defined by its central time point $t$ and its width $m$. The expanding-window cross-validation is achieved by considering only the time points within the window of the center point $t$. Note that windows corresponding to time points located at the beginning and end are truncated, i.e., their lengths are smaller than $m$. Therefore, the window size $m$ has to be sufficiently long to cover the necessary information of the dynamics.

#### S1.3 Extracting interaction probabilities

Computing the matrix of interaction probability, as well as the intrinsic growth rate and *per capita* interaction strength, for each time point $t$ is achieved as follows. First, the localization parameter $\theta(t)$ is estimated using all species (full model). Second, to apply the model averaging technique, we consider for a given species $i$ (response variable) all possible combinations of species $j$ (explanatory variable) that interact with $i$ (all possible sub-models). This defines the term $c_{ijk}$ where $c_{ijk} = 1$ if species $j$ interacts with $i$ in combination $k$, and $c_{ijk} = 0$ otherwise. For example, in a community of three species, the following combination of interactions $\ln(n_i(t+1)/n_i(t)) = r_i(t) + \alpha_{i1}(t) \cdot n_1(t) + \alpha_{i3}(t) \cdot n_3(t)$ leads to $c_{i1k} = 1$, $c_{i2k} = 0$, and $c_{i3k} = 1$. Third, for each combination $k$, we perform a local linear regression at time $t$, using the value $\theta(t)$ of the corresponding window, from which we extract the parameter values $r_{ik}(t)$ and $\alpha_{ijk}(t)$ as well as the $AIC_{ik}(t)$ and $BIC_{ik}(t)$. Fourth, we compute the AIC and BIC weights ($\omega_{ik}^{AIC}(t)$ and $\omega_{ik}^{BIC}(t)$). The BIC weight is given by equation (2), while for the AIC weight, we replace BIC by AIC [30, 31, 29]. Fifth, the probabilities of interaction at the time $t$ are computed by equation (3). Finally, we compute the average parameter estimators as follows:

$$\hat{r}_i(t) = \frac{\sum_k \hat{r}_{ik} \cdot c_{ijk} \cdot \omega_{ik}^{BIC}(t)}{\sum_k c_{ijk} \cdot \omega_{ik}^{BIC}(t)} \quad \text{and} \quad \hat{\alpha}_{ij}(t) = \frac{\sum_k \hat{\alpha}_{ijk} \cdot c_{ijk} \cdot \omega_{ik}^{BIC}(t)}{\sum_k c_{ijk} \cdot \omega_{ik}^{BIC}(t)} \tag{S2}$$

For the AIC, we simply replace BIC with AIC. We repeat all these steps for each species $i$ (response variable) and at each time step.

### S2 Synthetic data in a constant environment

We validate our method using synthetic random networks and semi-synthetic networks. The synthetic random networks are entirely randomly sampled, whereas the semi-synthetic networks make use of interguild empirical networks from the Web of Life (`https://www.web-of-life.es`).

Synthetic data is simulated from a discrete-time Lotka-Volterra model with environmental noise. The equation is given by:

$$n_i(t+1) = n_i(t) \cdot \exp\left( r_i + \sum_{j=1}^{S} \alpha_{ij} \cdot n_j(t) + \epsilon_i(t) \right) \qquad i = 1, ..., S. \tag{S3}$$

The random variables $\epsilon_i(t)$ represent the environmental noise and are drawn independently at random from a centered normal distribution with a standard deviation proportional to the $r_i$, i.e., $\epsilon_i(t) \sim \mathcal{N}(0, r_i \cdot \sigma_e)$.

#### S2.1 Random networks

For synthetic random networks, the protocol to generate the intrinsic growth rates and *per capita* interactions is as follows:

1. We set all intraspecific interactions, i.e., diagonal elements of the interaction matrix $\boldsymbol{\alpha}$, to $\alpha_{ii} = -1$.
2. We randomly choose a fraction $c$ of non-zero interspecific interactions, i.e., off-diagonal elements of the interaction matrix $\alpha_{ij} \quad (i \neq j)$. The fraction $c$ is the

connectance.

3. The non-zero interspecific interaction values (sampled at point 2) are drawn from a normal distribution of mean 0 and standard deviation $\sigma$, i.e., $\alpha_{ij} \sim \mathcal{N}(0, \sigma)$. The standard deviation $\sigma$ represents the amplitude in the variability of the interspecific interactions over the intraspecific ones, i.e., the "relative" strength of the interspecific over intraspecific interaction.

4. We simulate the population density at equilibrium $\mathbf{N}^*$ following a log-normal distribution $\log(n_i^*) \sim \mathcal{N}(0, 1)$.

5. We calculate the corresponding vector of intrinsic growth rates by setting $r_i = -\sum_j^S \alpha_{ij} n_j^*$.

To evaluate the robustness of our method, we vary the "relative" strengths of interactions $\sigma$, connectance $c$ and community size $S$. For each set of parameters, we simulate the population dynamics of 100 communities. Parameters used for simulation can be found in Table S1.

#### S2.2 Semi-synthetic empirical networks

For semi-synthetic networks, we download empirical networks from the Web of Life (`https://www.web-of-life.es/`). They are bipartite networks between two guilds A and B, where A and B represent plants and pollinators or hosts and parasites, respectively. From an empirical bipartite network, we generate its corresponding unipartite interaction matrix for a community (Supplementary Fig S1). Specifically, we duplicate the bipartite empirical matrix into two block matrices, in which one bipartite matrix is the transpose of the other ($\mathbf{W}_{BA}^T$ and $\mathbf{W}_{BA}$ in Supplementary Fig S1). The intraguild matrices are sampled following a protocol similar to that for the synthetic random networks.

We use plant-pollinator networks as representations of mutualistic interactions, where

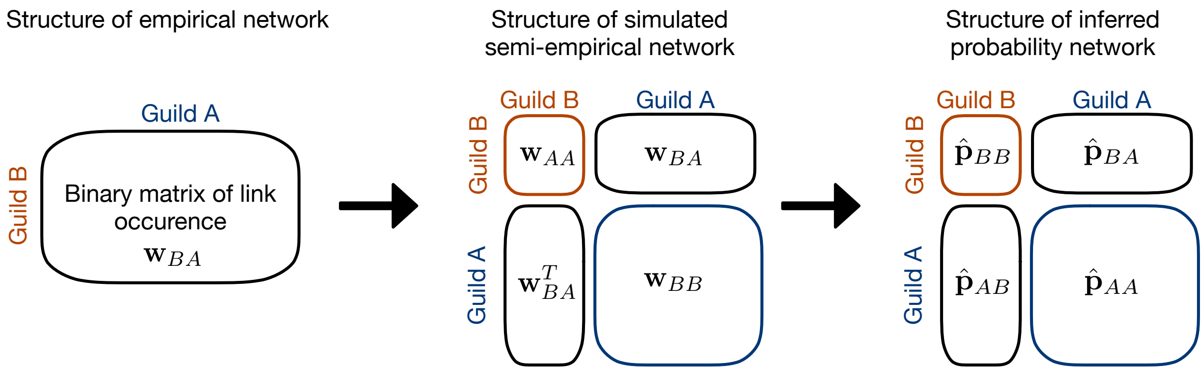

**Supplementary Fig S1. Procedure to construct a community matrix from empirical networks informed by an observational approach.**

the two block bipartite matrices contain positive interaction values, i.e., $\mathbf{W}_{BA}, \mathbf{W}_{BA}^T > 0$. Mutualistic interaction matrices are generated as follows:

1. All intraspecific interaction strengths are set to $-1$.

2. Intraguild interactions are sampled following the process of random networks. However, as these interactions are assumed to be competitions, their values follow a folded normal distribution $\alpha_{ij} \sim -|\mathcal{N}(0, \sigma_w)|$.

3. Interspecific interactions between plants and pollinators are sampled following the protocol of Saavedra et al. [12]. In particular, they are drawn from a log-normal distribution $ln(\alpha_{ij}) \sim \mathcal{N}(\gamma \cdot y_{ij}/d_i, \sigma_b)$, where $y_{ij}$ indicates the presence or absence of interaction between species $i$ and $j$, while $d_i$ is the number of interactions of species $i$, and $\gamma$ is the coefficient of mutualistic strength.

We use host-parasite networks as representations of antagonistic interactions, where the two block bipartite matrices contain opposite interaction values, i.e., $\mathbf{W}_{BA} < 0, \mathbf{W}_{BA}^T > 0$. Antagonistic interaction matrices are generated as follows:

1. All intraspecific interaction strengths are set to $-1$.

2. Intraguild interactions are all set to 0, as there is no information about intra-host interactions while intra-parasite interactions are less likely to exist.

3. Interspecific interactions of hosts on parasites follow a log-normal distribution $\ln(\alpha_{ij}) \sim \mathcal{N}(\epsilon \cdot \gamma \cdot y_{ij}/d_i, \sigma_b)$, where $\epsilon$ represents the yield coefficient. By contrast, the interspecific interactions of parasites on hosts follow a log-normal distribution with a superimposed negative sign $\ln(-\alpha_{ij}) \sim \mathcal{N}(\gamma \cdot y_{ij}/d_i, \sigma_b)$ because the parasites are harmful to the hosts.

| **Parameter** | **Description** | **Value** |
|---|---|---|
| **Random synthetic network** | | |
| $S$ | Number of species | 5, 7 & 10 |
| $\sigma$ | SD for normal distribution of interspecific interaction strengths | 0.01, 0.05, & 0.1 |
| $c$ | Connectance | 0.05, 0.1, 0.2, 0.3, & 0.5 |
| **Semi-synthetic network** | | |
| $\gamma$ | Interaction coefficients | 0.0005 |
| $\sigma_b$ | SD for normal distribution of inter-guild interaction strengths | 0.1 |
| $\sigma_w$ | SD for normal distribution of intra-guild interaction strengths | 0.005 |
| $c$ | Connectance for intra-guild in plant-pollinator systems | 0.3 |
| $\epsilon$ | Yield in host-parasite interactions | 1 |
| $\sigma_e$ | SD for normal distribution of environmental noise | 0.01 |

Table S1: Parameter values used for simulation

#### S2.3 Statistics to evaluate the performance of the MA-LVmap

The result of the MA-LVmap approach for each simulated community is a matrix containing probabilities of pairwise links. We use it to sample 1000 matrices of pairwise links, i.e., matrices that contain only 0 and 1 values. As a result, for each community, we obtain a distribution of the following metrics:

- Ratio between inferred and true connectance
- Sensitivity – the fraction of absent interactions correctly detected, computed as the ratio between correctly inferred zero interspecific interactions and true zero interspecific interactions
- Specificity – the fraction of present interactions correctly detected, computed as the ratio between correctly inferred non-zero interspecific interactions and true non-zero interspecific interactions
- Difference between true and inferred nestedness
- Difference between true and inferred modularity

Note that connectance for the random network is the number of realized links over all possible links, whereas connectance for the empirical network is the number of realized links of the interguild matrix over all possible interguild links. Nestedness and modularity are only applicable to empirical networks, as these indices are not relevant to random networks. We use the nestedness algorithm developed by Almeida-Neto et al. [52], implemented in the function *nestednodf* from the *vegan* package [53], and the modularity algorithm developed by Newman [54], implemented in the function *computeModules* from the *bipartite* package [55].

Another important note is that, in the inferred matrix $\hat{\mathbf{p}}$ of the empirical network, species $i$ in one guild may have a non-zero effect on species $j$ in the other guild, but not vice versa, because the selecting process is performed for each species separately. Consequently, entry $\hat{p}_{ij}$ is likely to differ from entry $\hat{p}_{ji}$, suggesting that plant (or host) $i$ may interact with pollinator (or parasite) $j$, but not vice versa. This results in two asymmetric interguild matrices of probability (figure S1).

### S3 Synthetic data in changing environments

Communities are always subject to changing environmental conditions, resulting in variations of connectance and network topology across time [56, 12]. To validate this ability, we simulate a Lotka-Volterra discrete-time model of a predator-prey community with two prey that share a common predator. The community experiences two different environments with a drop in connectance from 9 links in the first environment to 7 links in the second environment. The intrinsic growth rates, *per capita* interactions, and network links in the first environment are:

$$\mathbf{r_b} = \begin{pmatrix} 1.7 \\ 2.04 \\ 2.89 \end{pmatrix}, \boldsymbol{\alpha_b} = \begin{pmatrix} -2.55 & 0.391 & 0.17 \\ -0.51 & -1.7 & -0.51 \\ -0.68 & -0.60 & -1.7 \end{pmatrix} \text{ and } \boldsymbol{p_b} = \begin{pmatrix} 1 & 1 & 1 \\ 1 & 1 & 1 \\ 1 & 1 & 1 \end{pmatrix}, \tag{S4}$$

whereas the parameters in the second environment are:

$$\mathbf{r_f} = \begin{pmatrix} 1.70 \\ 2.04 \\ 1.53 \end{pmatrix}, \boldsymbol{\alpha_f} = \begin{pmatrix} -2.55 & 0.391 & 0 \\ -0.51 & -1.7 & -0.51 \\ 0 & -0.595 & -1.7 \end{pmatrix} \text{ and } \mathbf{p_f} = \begin{pmatrix} 1 & 1 & 0 \\ 1 & 1 & 1 \\ 0 & 1 & 1 \end{pmatrix}. \tag{S5}$$

The simulation runs for 600 time points where the environmental change starts around time point 300 with three levels of graduality, reflected by a sigmoid function. The values of intrinsic growth rates and *per capita* interaction strengths at each time point are as follows:

$$r(t) = r_b + (r_f - r_b) * f_c(t) \tag{S6}$$

$$\alpha(t) = \alpha_b + (\alpha_f - \alpha_b) * f_c(t) \tag{S7}$$

$$f_c(t) = \frac{1}{1 + e^{-\beta(t-300)}}. \tag{S8}$$

The parameter $\beta$ indicates the gradualness, where $\beta = 0$ means that there is no environmental change, $\beta = 0.01$ indicates an extremely gradual change, $\beta = 0.05$ indicates an intermediate gradual change, and $\beta = 0.9$ indicates an abrupt change.

### S4 Empirical time-series data

#### S4.1 Time series from microcosm experiment

As the first proof of concept, we apply the MA-LVmap to a microcosm experiment from Limberger et al. [39]. This microcosm experiment is comprised of two treatments: (i) constant light treatment and (ii) decreasing light treatment for different microbial communities. The microbial communities are combinations of five different functional groups: bacteria, edible algae, inedible algae, bacterivore, and omnivore. The experiment was conducted for 9 months. The decreasing light treatment was gradual, such that for the first 3 months, constantly high light was maintained, followed by 3 months of gradual light decline and 3 months of constant low light. Because the sampling was done three times per week, we interpolated the values for the days without sampling to obtain daily time series.

#### S4.2 Time series from mesocosm experiment

As the second proof of concept, we use the time-series data from a mesocosm experiment from Benincà et al. [32]. This time-series data is the result of a long-term mesocosm experiment involving a plankton community isolated from the Baltic Sea. The mesocosm experiment was conducted under relatively constant environmental conditions from 31 March 1989 until 20 October 1997 [57, 40, 32].

The mesocosm time-series data was retrieved from the Supporting Information from Benincà et al. [32]. The raw data contains the abundance of 10 functional groups (cyclopoids, calanoid copepods, rotifers, protozoa, nanoflagellates, picocyanobacteria, filamentous diatoms, ostracods, harpacticoids, and bacteria). They make up a community with four trophic levels. We focus on five functional groups – bacteria as the primary resource, picocyanobacteria and nanoflagellates as prey, and calanoid copepods and rotifers as predators – because other groups contain a large number of zero and missing values in their time-series data. In addition, we use a subset of the time-series data from 07 December 1990 until 24 April 1995, during which data were recorded at the highest frequency. This subset of time-series data still contains missing and exact-zero values. We replace exact-zero values with the corresponding minimum abundance values of each group divided by two. We then interpolate the missing values on the log scale of the population abundance to obtain daily measurements.

### S5 Supplementary figures

#### S5.1 Synthetic random networks

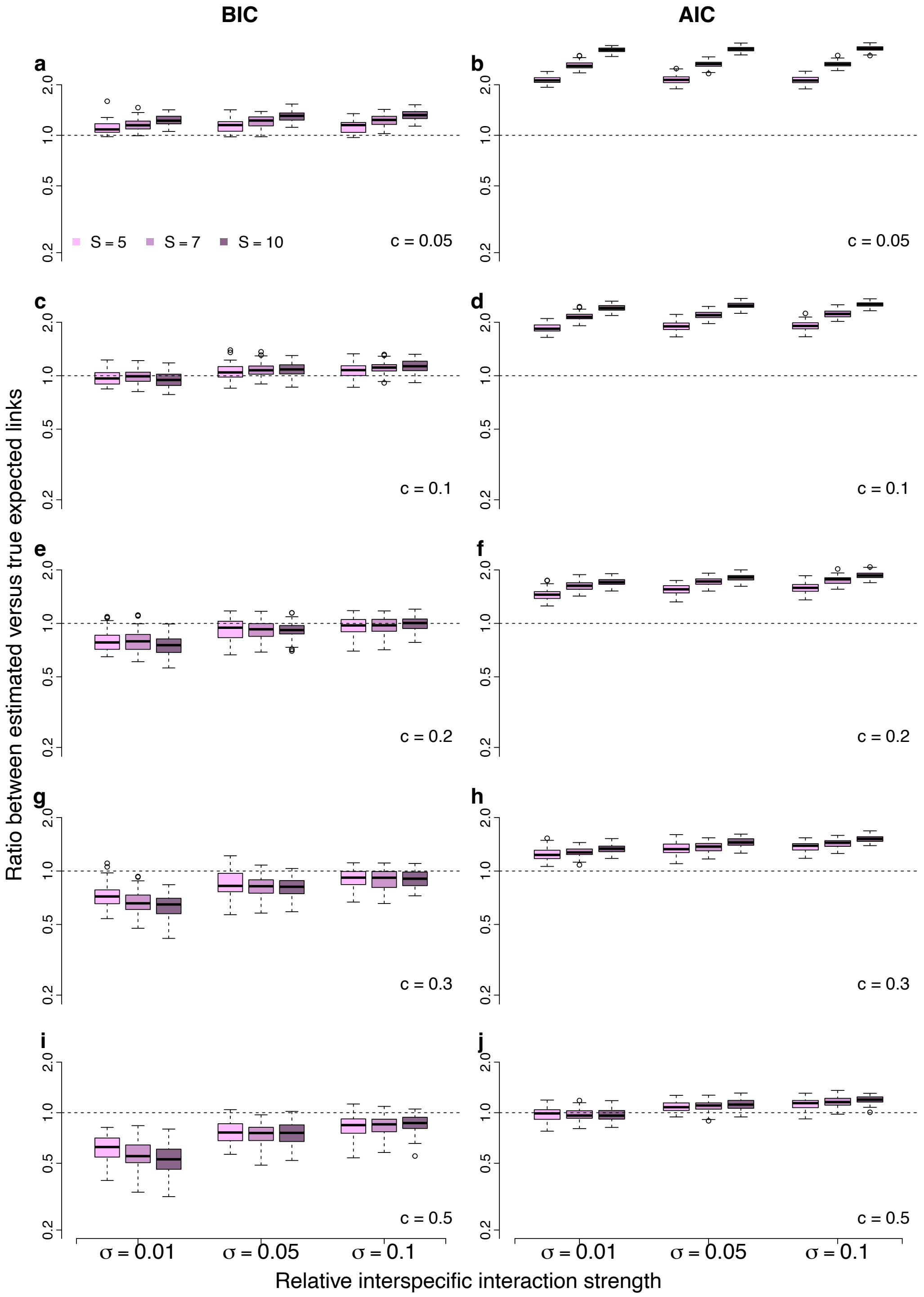

**Supplementary Fig S2.** Comparison between true and estimated expected links for random networks of different sizes ($S = (5,\ 7,\ 10)$), with different connectance ($c = (0.05,\ 0.1,\ 0.2,\ 0.3,\ 0.5)$), and different relative interspecific interaction strength ($\sigma = (0.01,\ 0.05,\ 0.1)$).**a, c, e, g, i**. Results using BIC weights. **b, d, f, h, j**. Results using AIC weights.

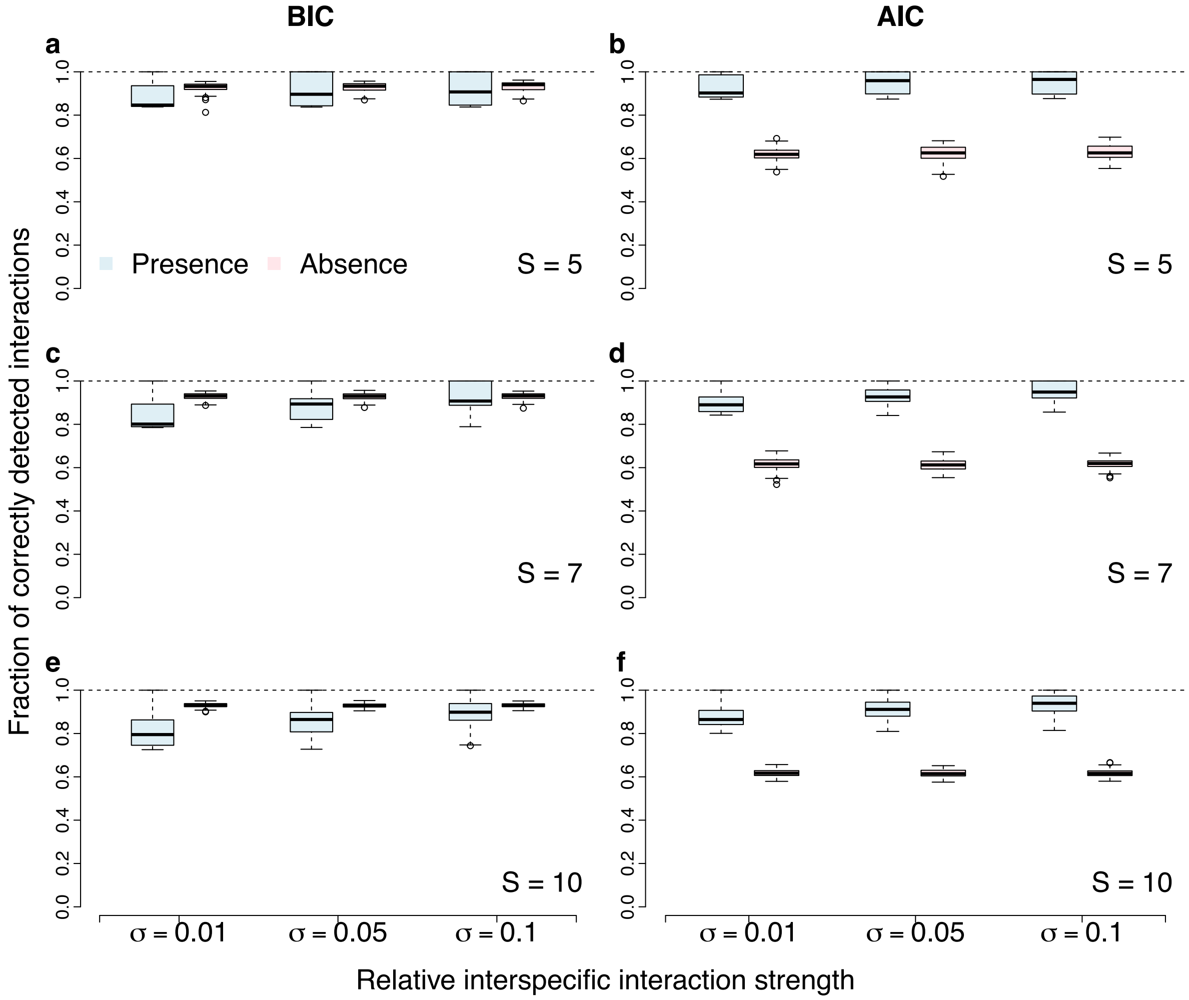

**Supplementary Fig S3.** Fraction of correctly detected presence and absence of interactions of the networks of different sizes ($S = (5,\ 7,\ 10)$) for $c = 0.05$. **a, c, e**. Results using BIC weights. **b, d, f**. Results using AIC weights.

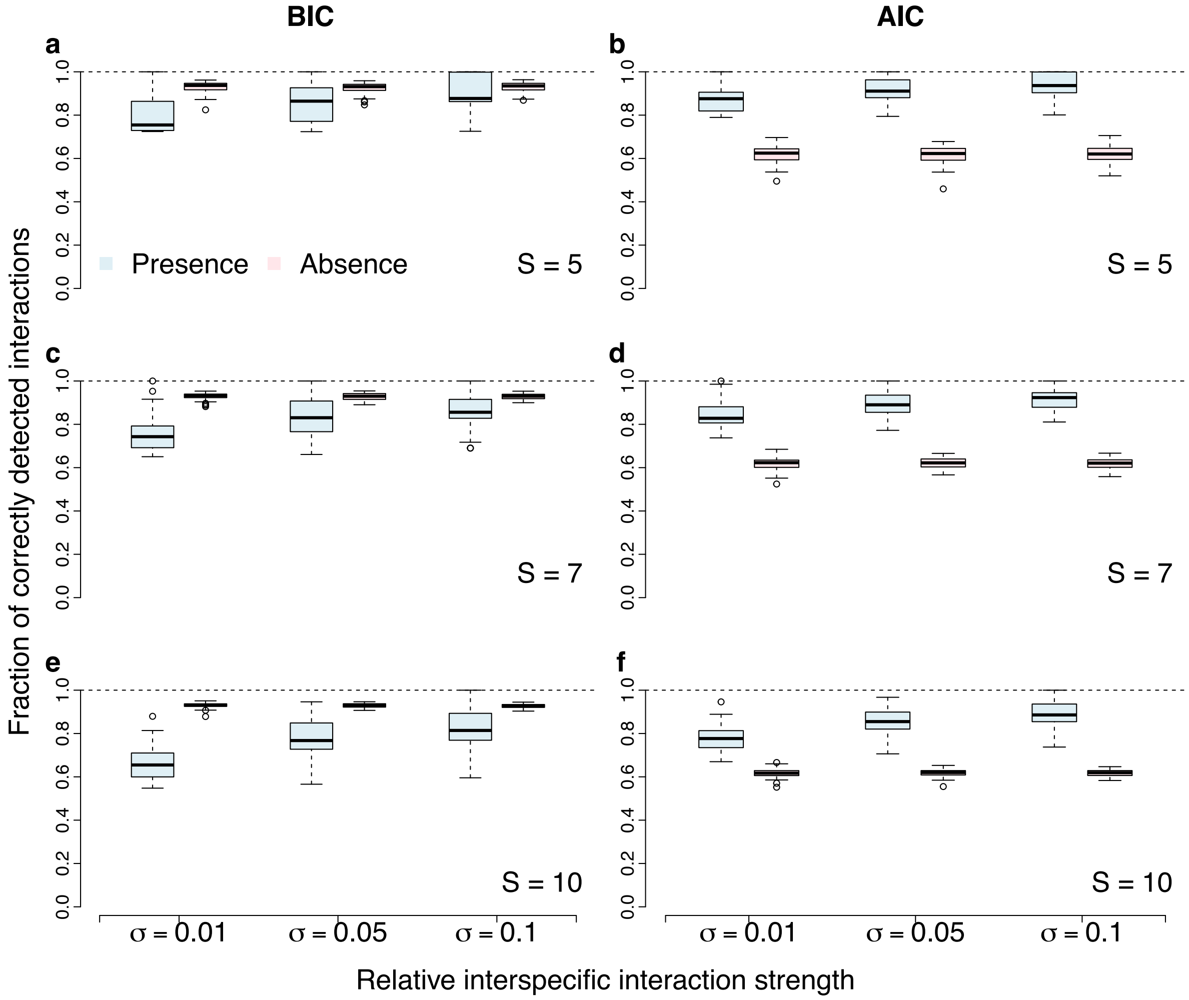

**Supplementary Fig S4.** Fraction of correctly detected presence and absence of interactions of the networks of different sizes ($S = (5,\ 7,\ 10)$) for $c = 0.1$. **a, c, e**. Results using BIC weights. **b, d, f**. Results using AIC weights.

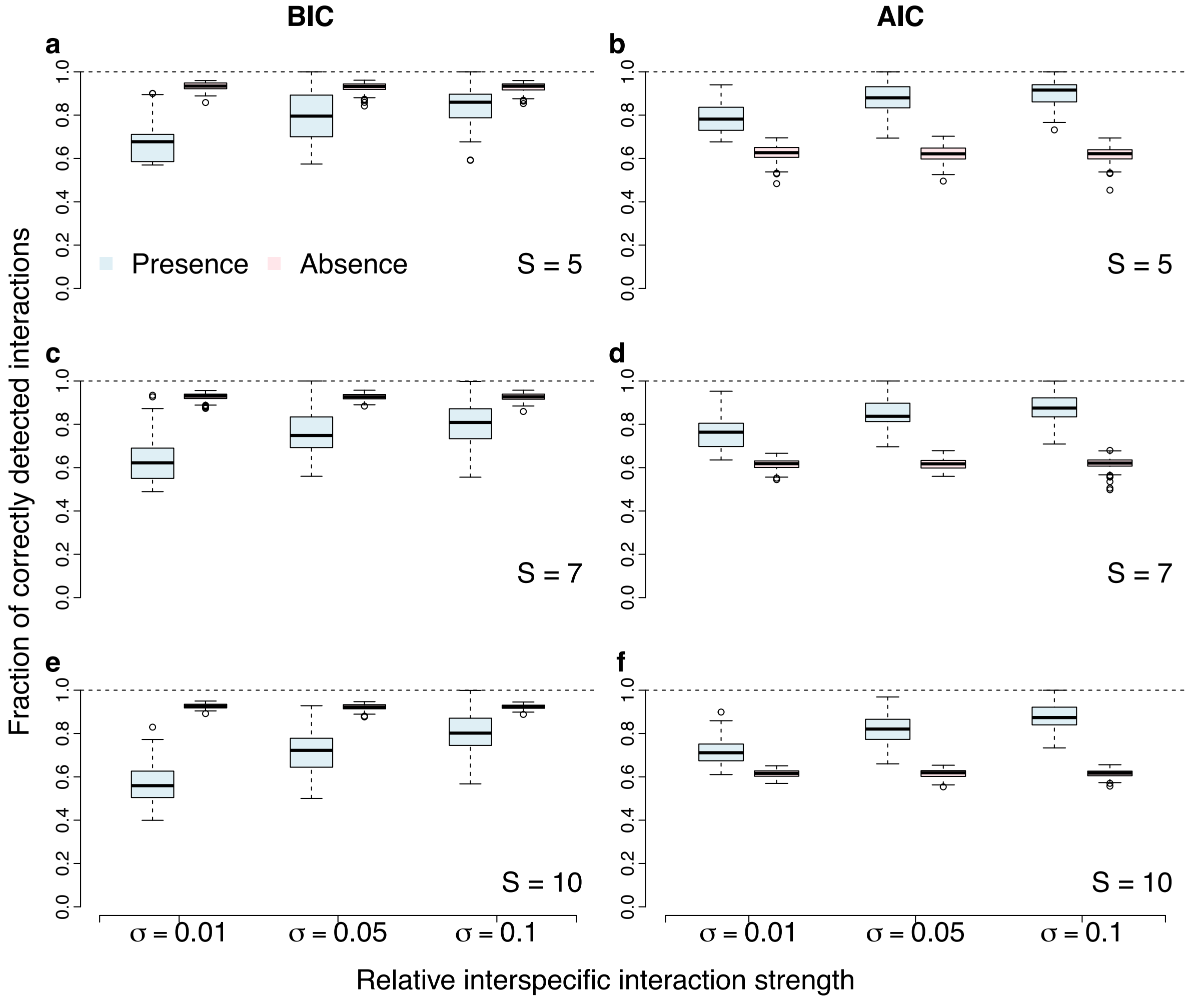

**Supplementary Fig S5.** Fraction of correctly detected presence and absence of interactions of the networks of different sizes ($S = (5,\ 7,\ 10)$) for $c = 0.2$. **a, c, e**. Results using BIC weights. **b, d, f**. Results using AIC weights.

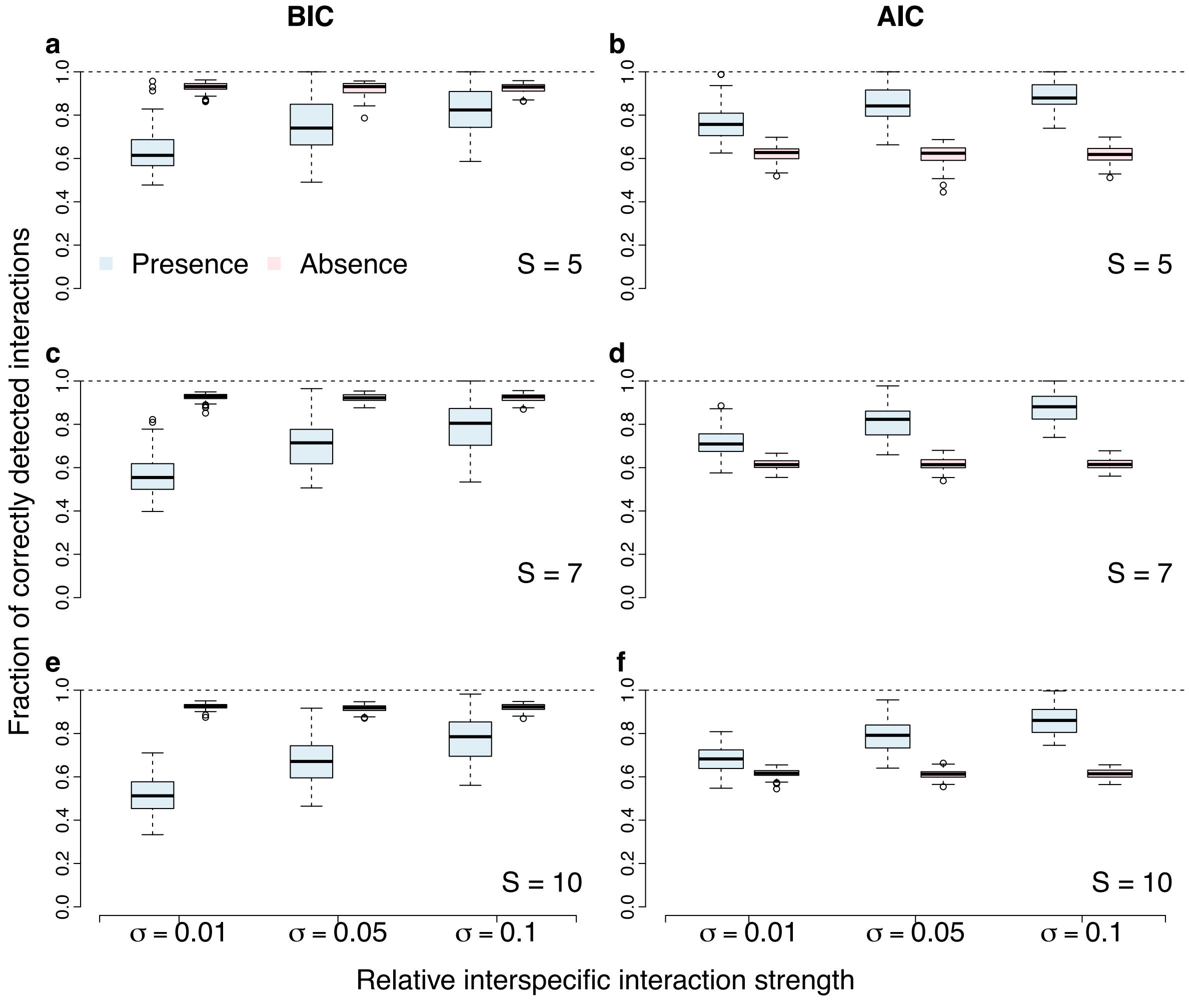

**Supplementary Fig S6.** Fraction of correctly detected presence and absence of interactions of the networks of different sizes ($S = (5,\ 7,\ 10)$) for $c = 0.3$. **a, c, e**. Results using BIC weights. **b, d, f**. Results using AIC weights.

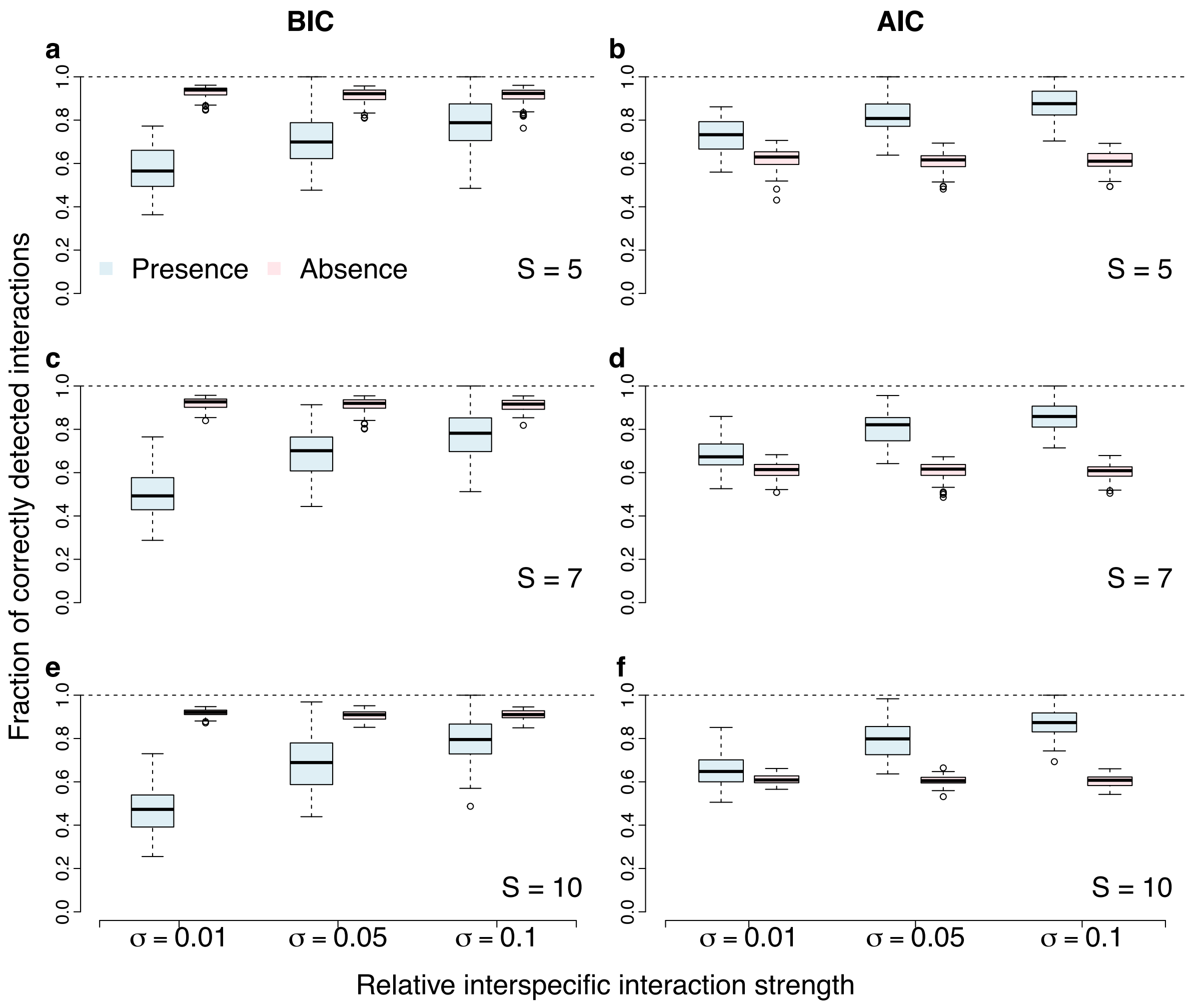

**Supplementary Fig S7.** Fraction of correctly detected presence and absence of interactions of the networks of different sizes ($S = (5,\ 7,\ 10)$) for $c = 0.5$. **a, c, e**. Results using BIC weights. **b, d, f**. Results using AIC weights.

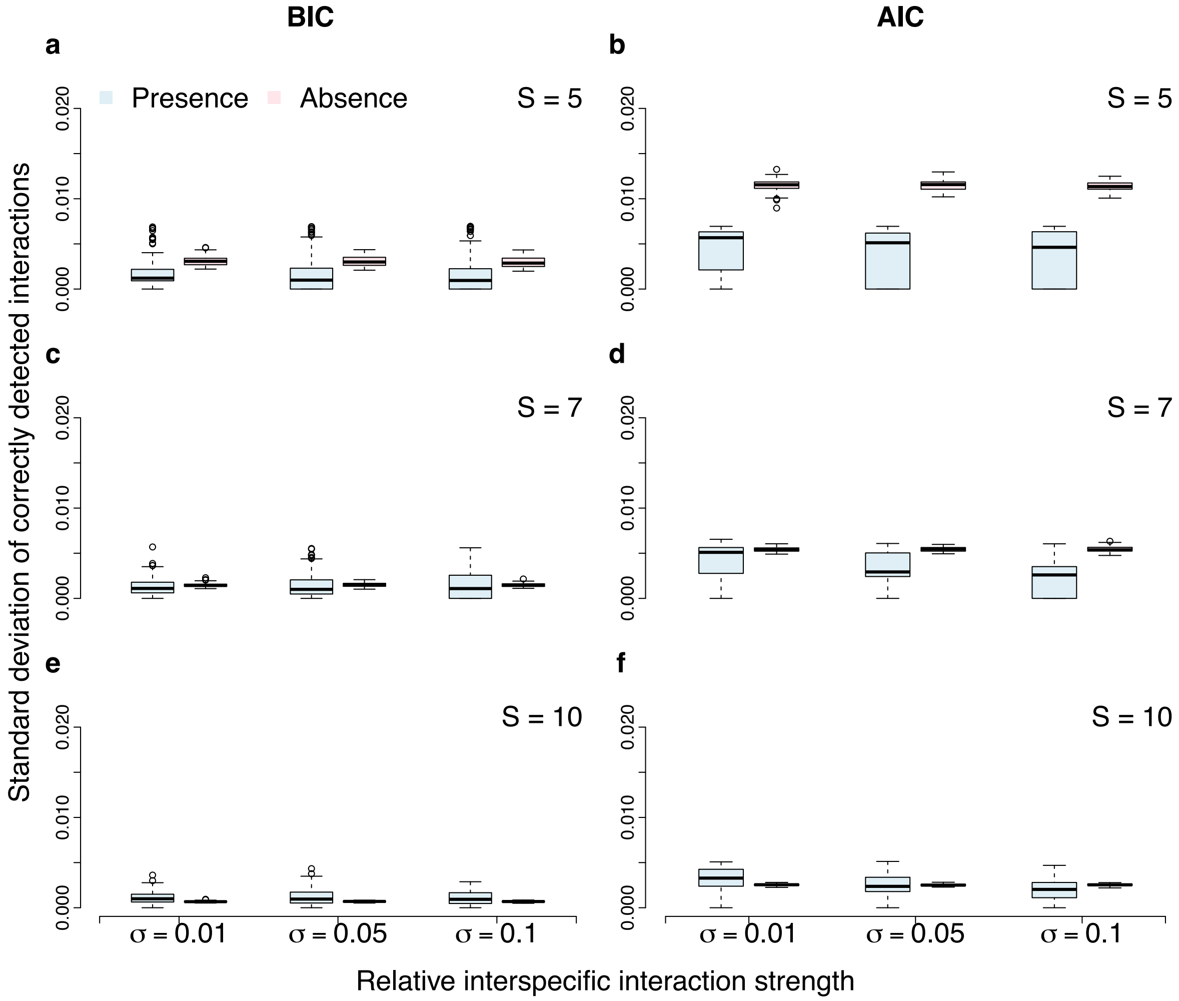

**Supplementary Fig S8.** Standard deviation of the distribution of the correctly detected presence and absence of interactions. Results obtained from 1000 adjacency binary matrices sampled from the inferred interaction probabilities. Networks have different sizes ($S = (5,\ 7,\ 10)$) for $c = 0.05$. **a, c, e**. Results using BIC weights. **b, d, f**. Results using AIC weights.

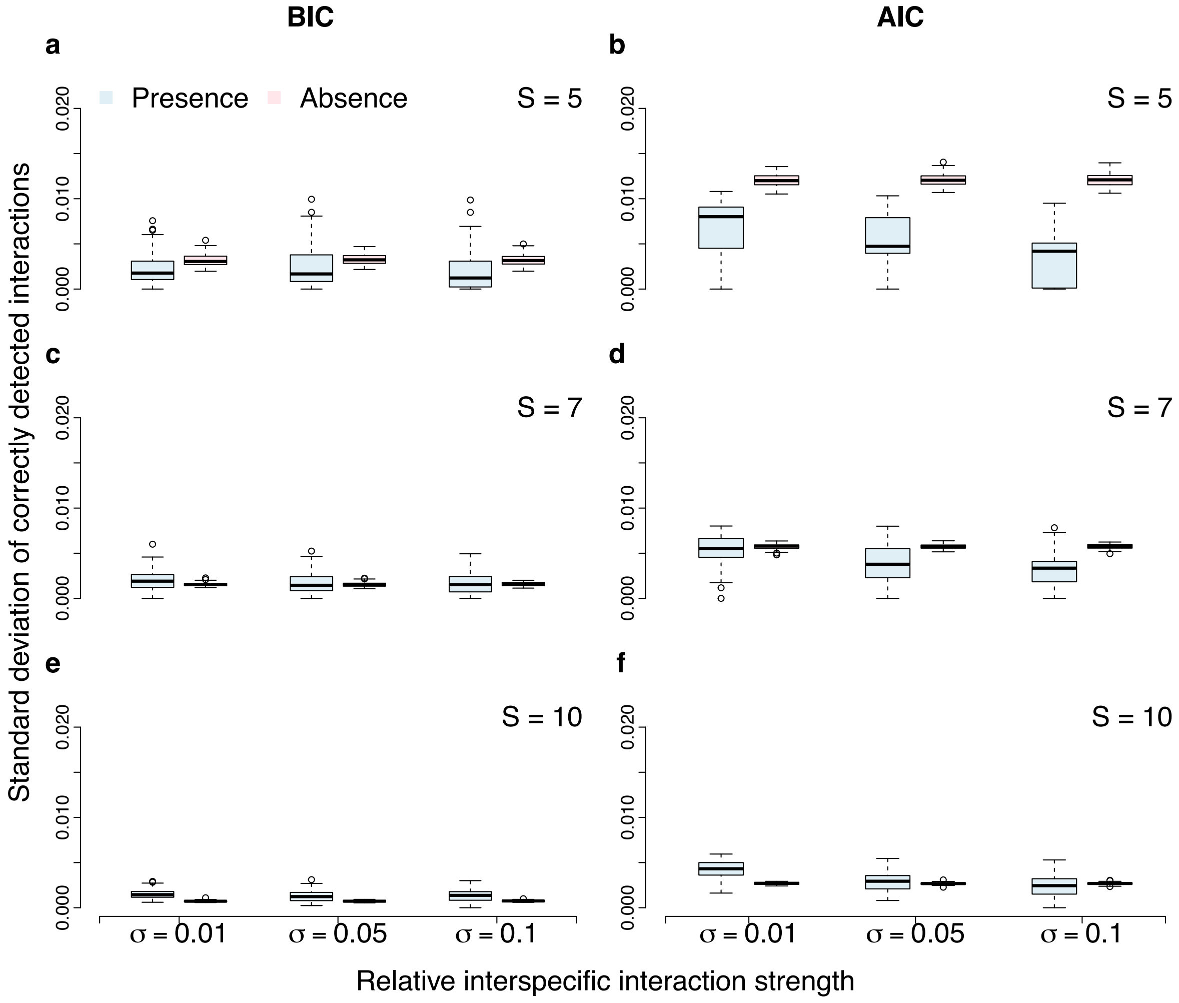

**Supplementary Fig S9.** Standard deviation of the distribution of the correctly detected presence and absence of interactions. Results obtained from 1000 adjacency binary matrices sampled from the inferred interaction probabilities. Networks have different sizes ($S = (5,\ 7,\ 10)$) for $c = 0.1$. **a, c, e**. Results using BIC weights. **b, d, f**. Results using AIC weights.

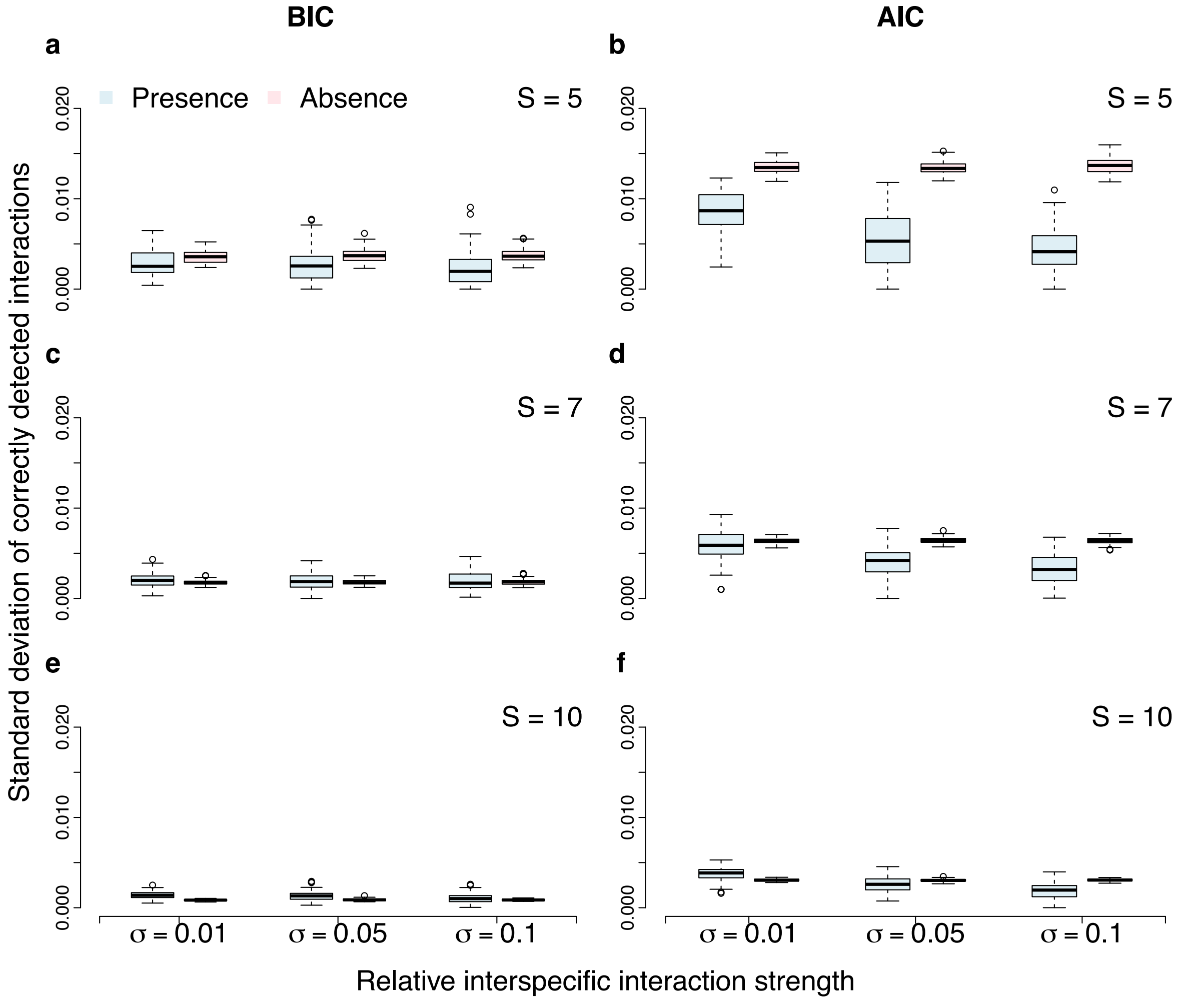

**Supplementary Fig S10.** Standard deviation of the distribution of the correctly detected presence and absence of interactions. Results obtained from 1000 adjacency binary matrices sampled from the inferred interaction probabilities. Networks have different sizes ($S = (5,\ 7,\ 10)$) for $c = 0.2$. **a, c, e**. Results using BIC weights. **b, d, f**. Results using AIC weights.

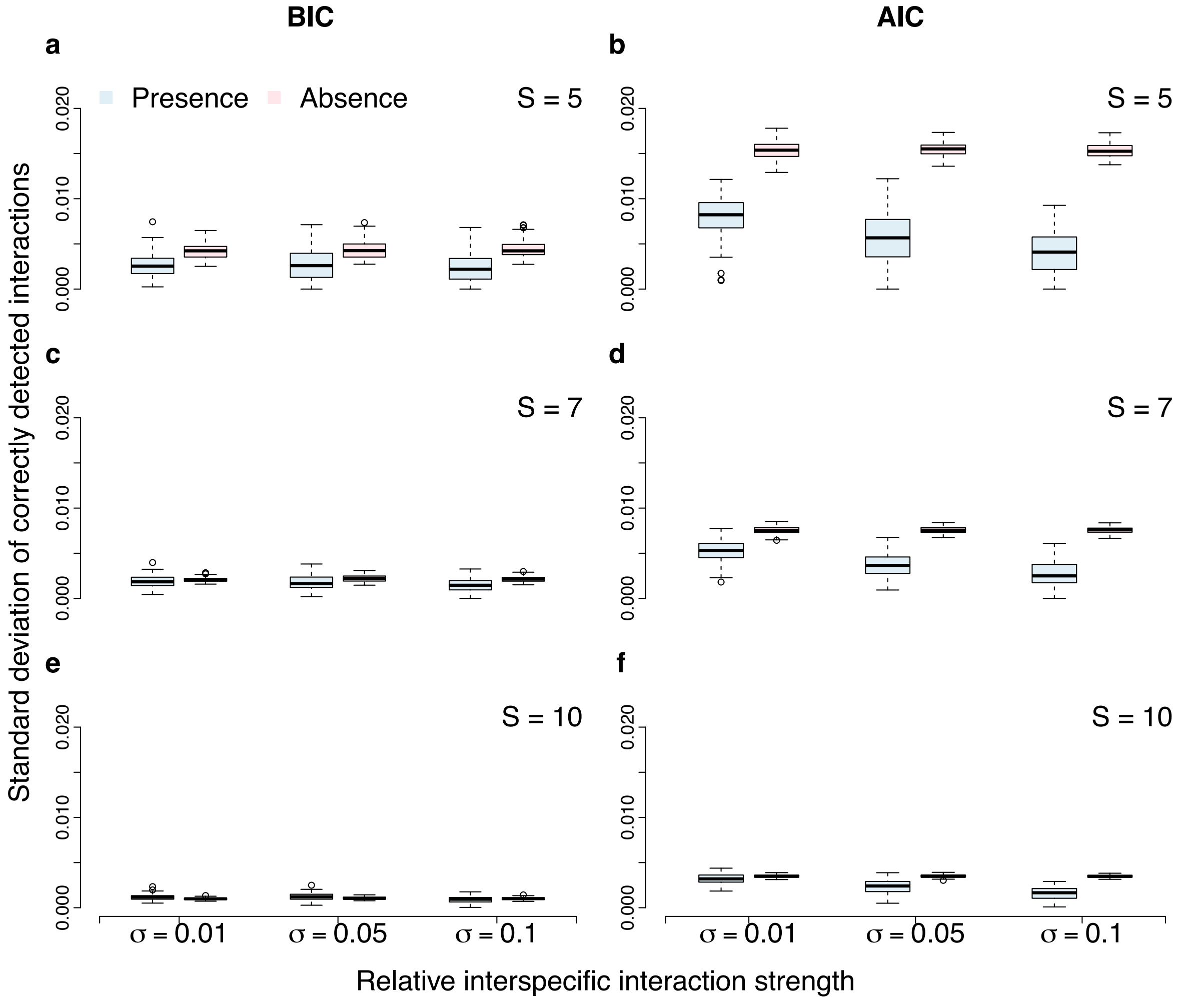

**Supplementary Fig S11.** Standard deviation of the distribution of the correctly detected presence and absence of interactions. Results obtained from 1000 adjacency binary matrices sampled from the inferred interaction probabilities. Networks have different sizes ($S = (5,\ 7,\ 10)$) for $c = 0.3$. **a, c, e**. Results using BIC weights. **b, d, f**. Results using AIC weights.

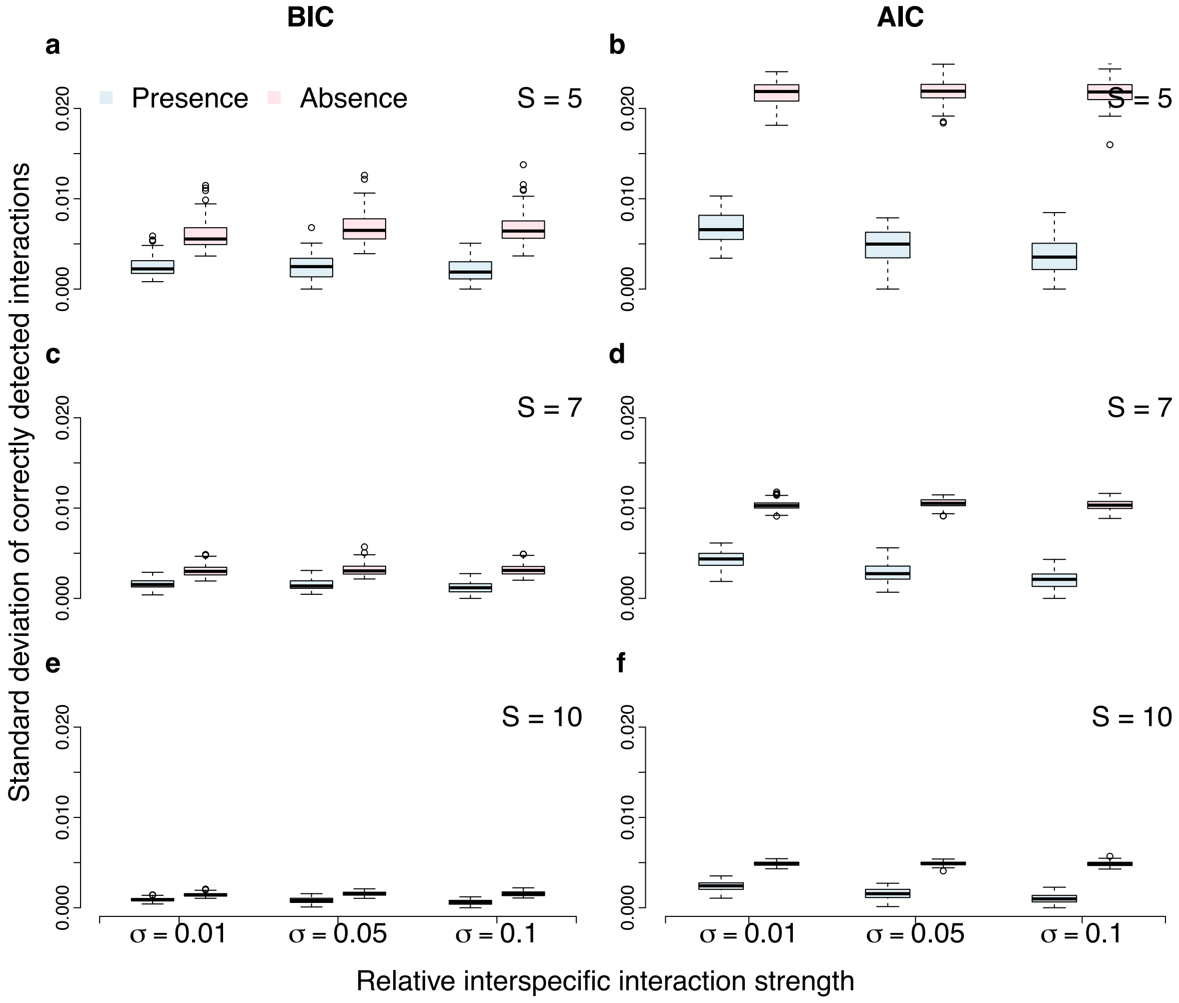

**Supplementary Fig S12.** Standard deviation of the distribution of the correctly detected presence and absence of interactions. Results obtained from 1000 adjacency binary matrices sampled from the inferred interaction probabilities. Networks have different sizes ($S = (5,\ 7,\ 10)$) for $c = 0.5$. **a, c, e**. Results using BIC weights. **b, d, f**. Results using AIC weights.

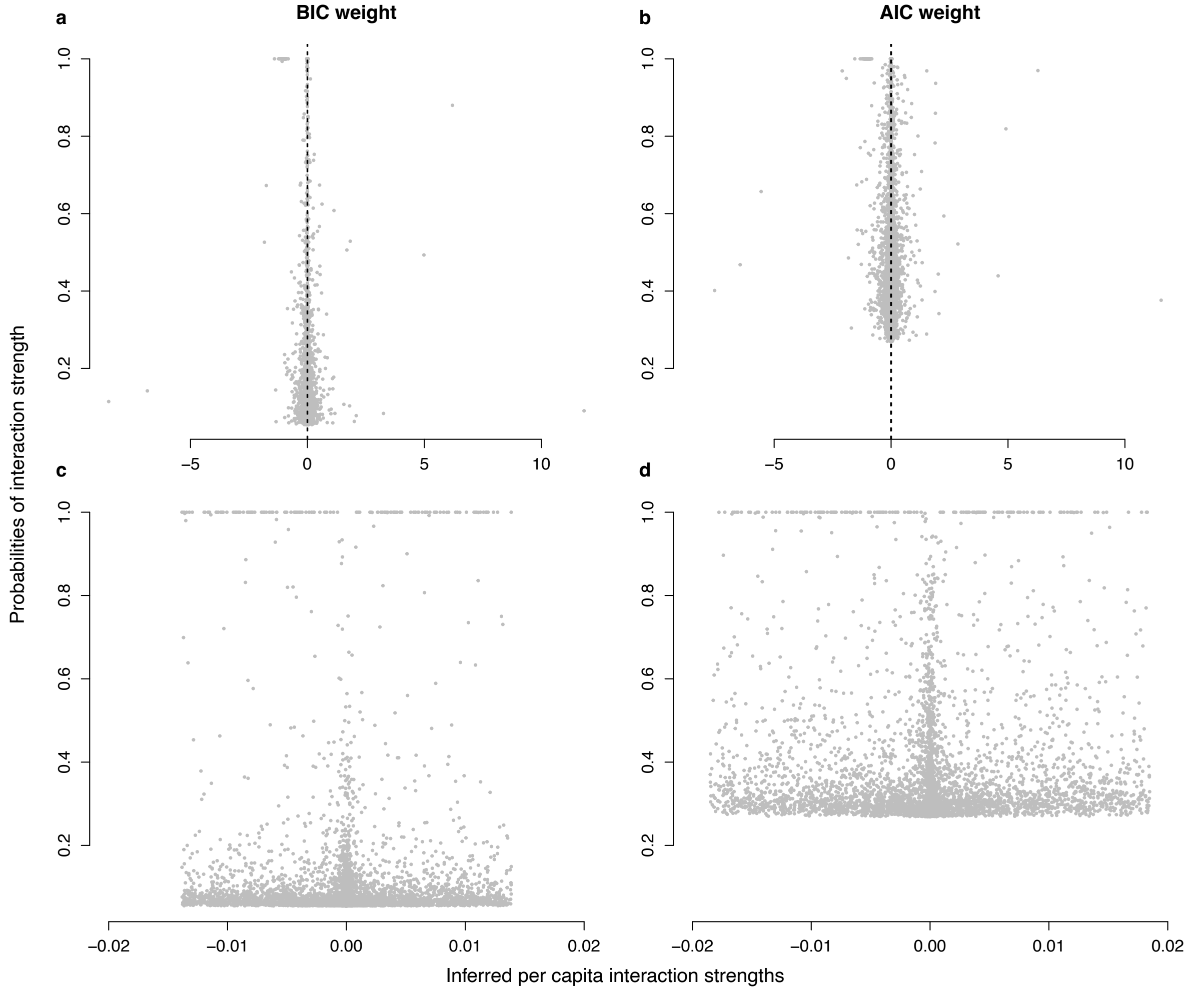

**Supplementary Fig S13.** Correlation between inferred interaction strengths and interaction probabilities for a community of $S = 10$ species, small connectance $c = 0.1$, small interaction strength $\sigma = 0.01$, **a, b**. All inferred *per capita* interaction strengths. **c, d**. Zoom in on the small values of inferred *per capita* interaction strengths.

#### S5.2 Semi-synthetic empirical networks

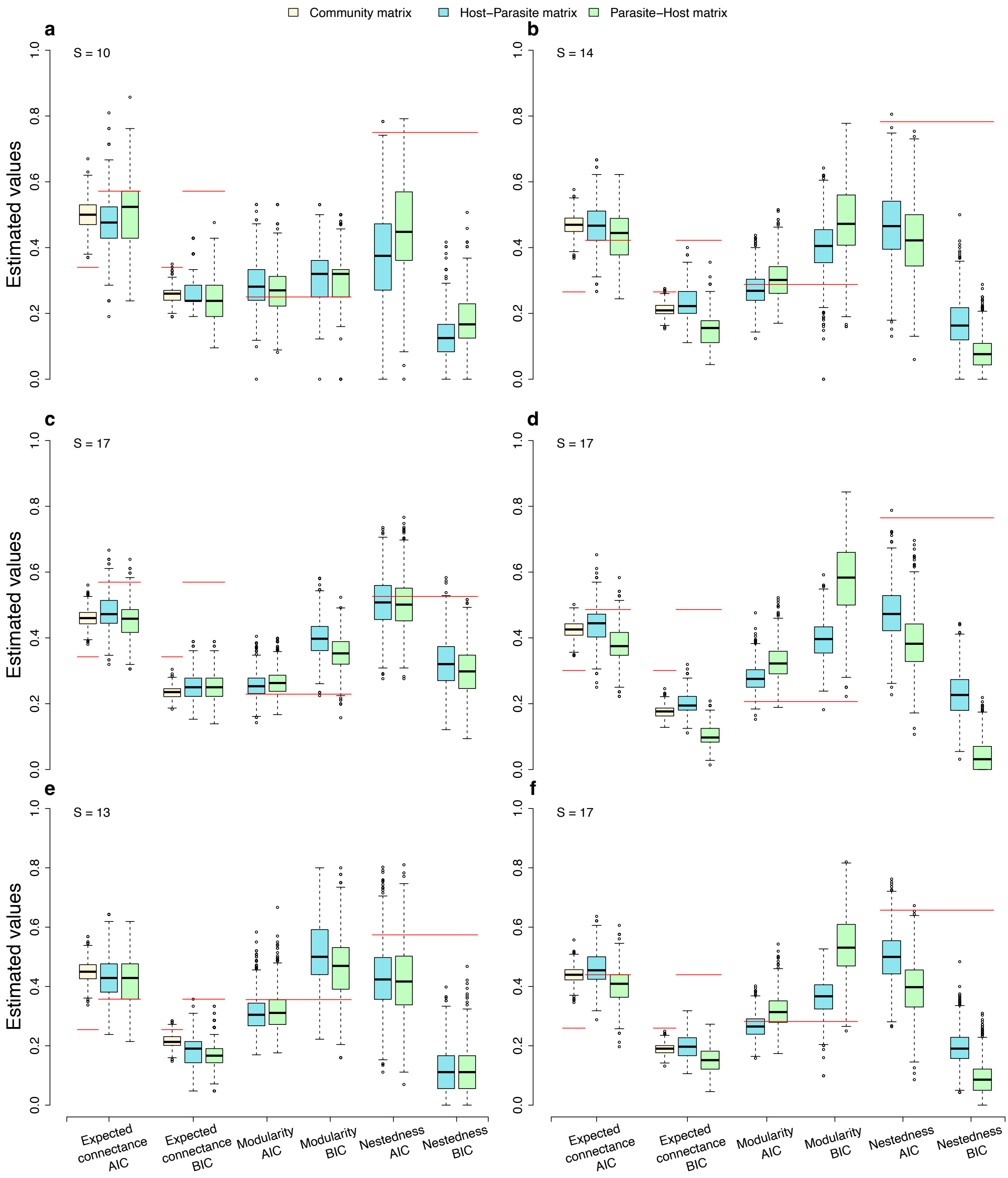

**Supplementary Fig S14. Inference of network architecture for antagonistic networks.** Results are from 1000 randomizations of the inferred matrix for each community. Red lines indicate true values. Other parameters are $\sigma_b = 0.1$ for the interguild interaction strengths. We assume that there are no intraguild interaction strengths among hosts and among parasites.

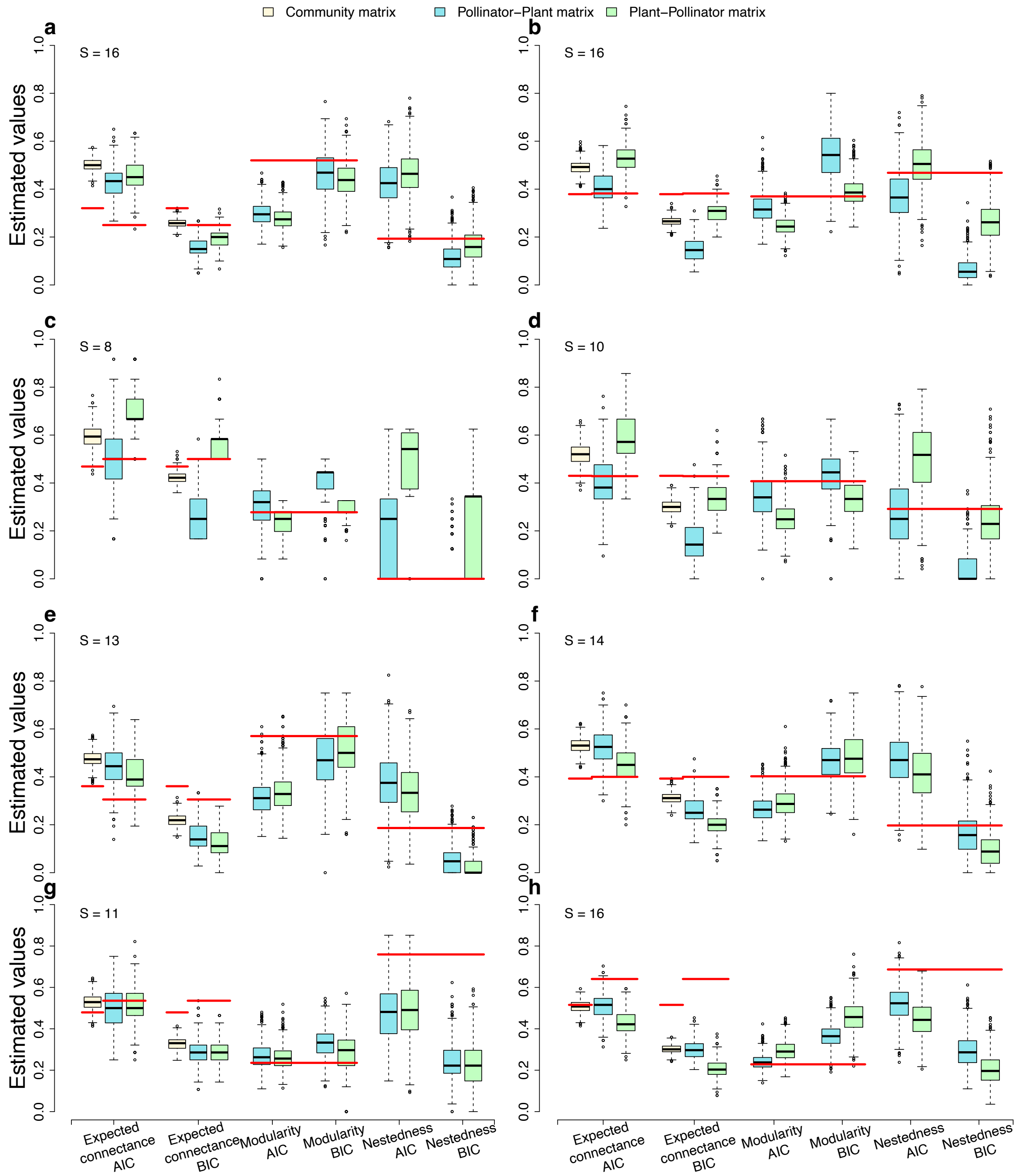

**Supplementary Fig S15. Inference of network architecture for mutualistic networks.** Results are from 1000 randomizations of the inferred matrix for each community. Red lines indicate true values. Other parameters are $\sigma_b = 0.1$ for interguild interaction strengths and $\sigma_w = 0.005$ for intraguild interaction strengths among plants and among pollinators.

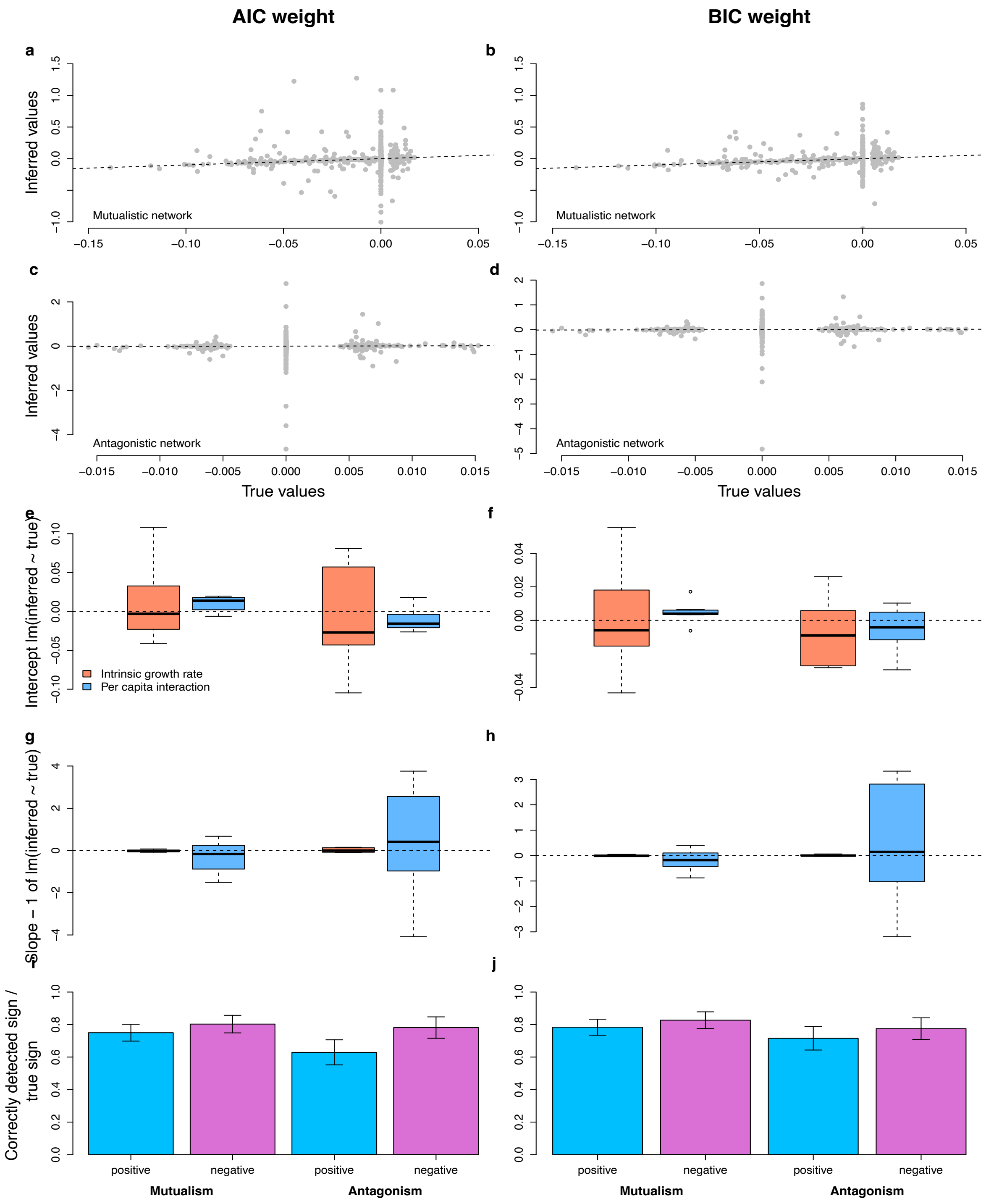

**Supplementary Fig S16. Inference of intrinsic growth rates and *per capita* interaction strengths for both mutualistic and antagonistic networks. a-d.** Correlation between true *per capita* interaction strengths and estimated *per capita* interaction strengths. Dashed lines indicate perfect correlation $y = x$. **e-h.** Intercept and slope of the linear regression on the true values versus estimated values. Horizontal dashed lines indicate zero intercept and zero (slope - 1) of the linear regression. **i, j.** Ratios of correctly detected positive and negative interactions to true positive and negative interactions, respectively. Vertical bars indicate 95% CI.

#### S5.3 Comparison with regularisation techniques

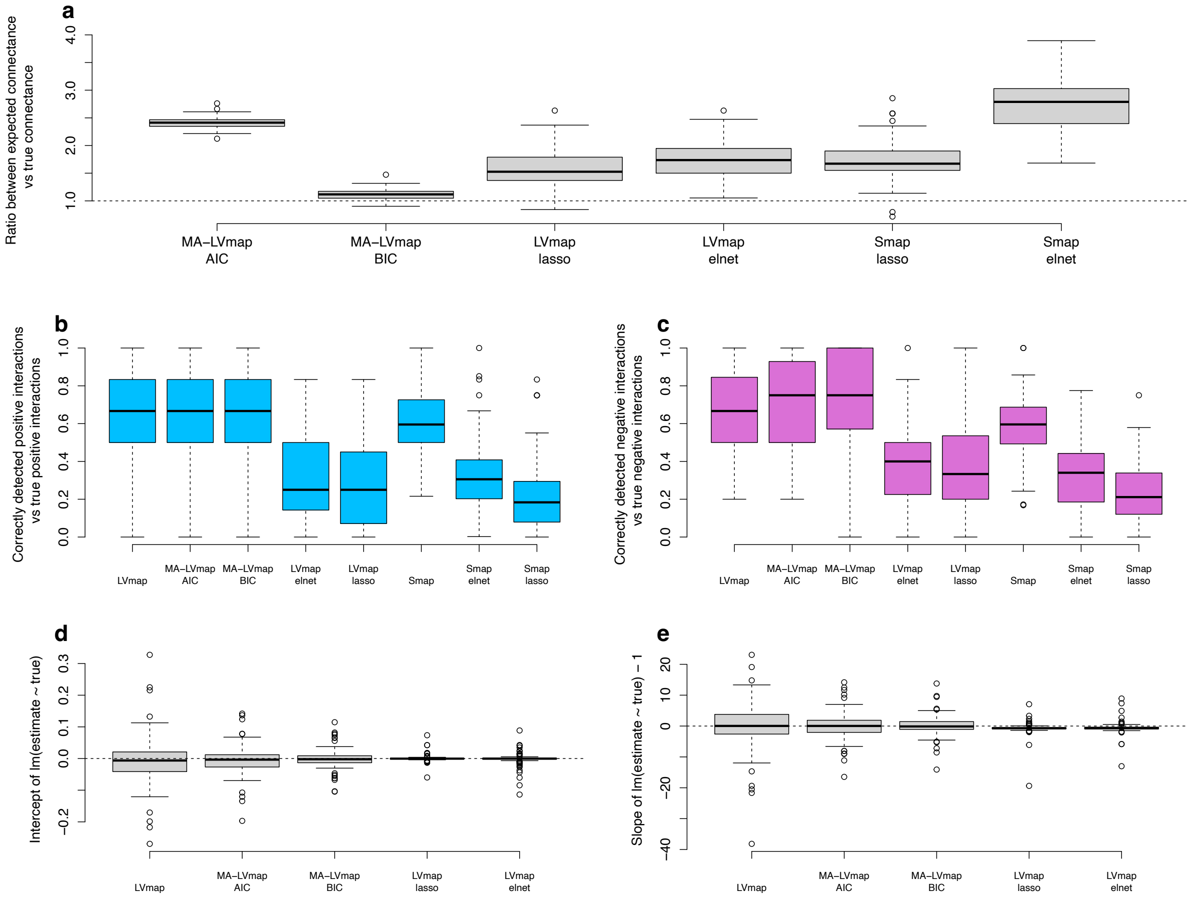

**Supplementary Fig S17. Comparing results using different techniques: MA-LVmap (AIC weight), MA-LVmap (BIC-weight), LV-map lasso, LV-map elasticnet, Smap, Smap lasso, Smap elastic net. a**. Connectance. **b** Positive interactions. **c** Negative interactions. **d, e** Correlation between true and inferred intraspecific interaction strengths reflected through the intercept and the slope of the linear regression between true and inferred values. Simulation for a community of size $S = 10$ species, small connectance $c = 0.1$, and small interaction strength $\sigma = 0.01$

#### S5.4 Synthetic data in changing environmental conditions

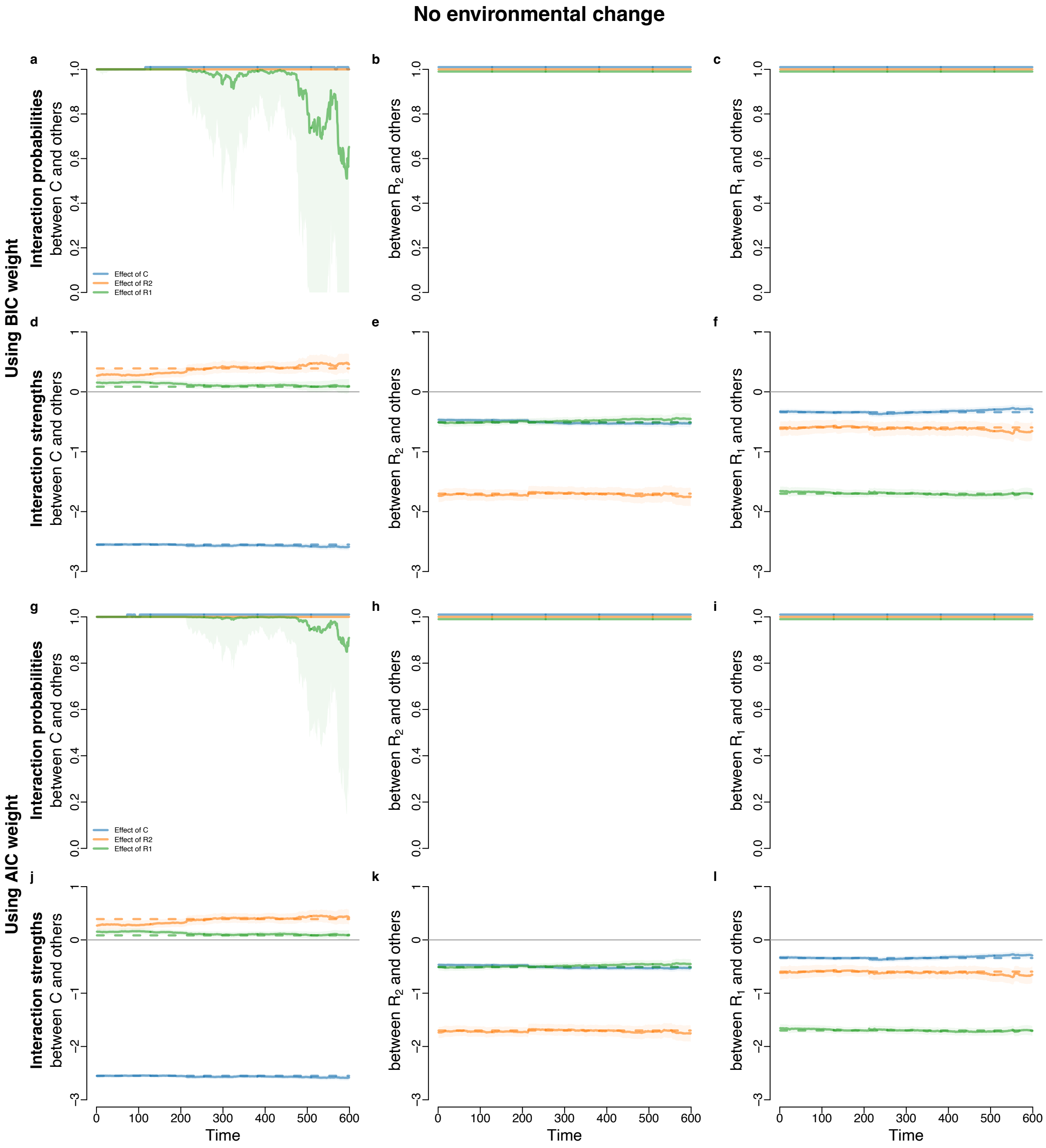

**Supplementary Fig S18.** Inference of interaction probability and interaction strength for the synthetic predator-prey system with no change in environmental condition. Dashed lines indicate true values and solid lines indicate inferred values. Transparent ribbons indicate 95% confidence intervals.

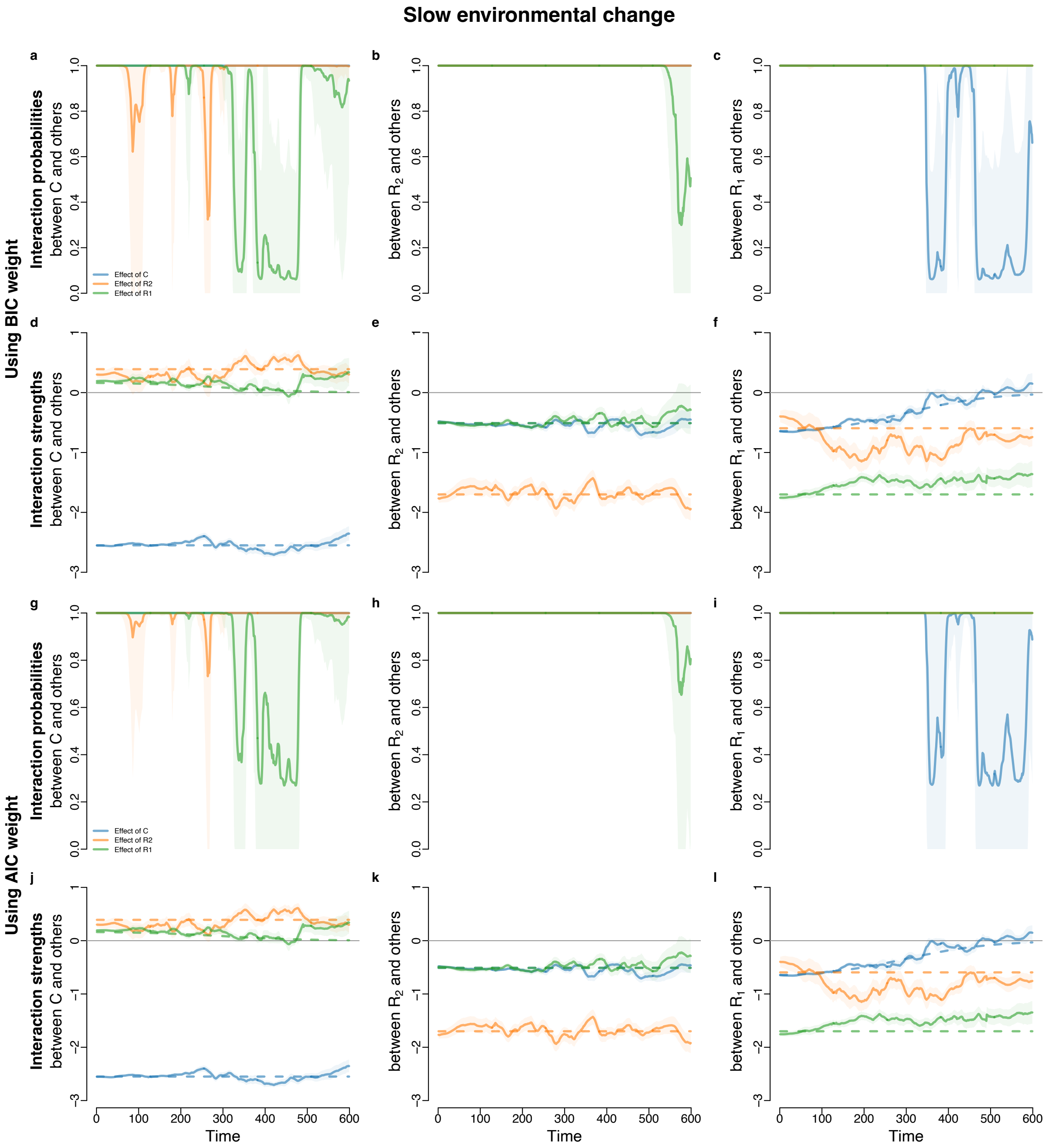

**Supplementary Fig S19.** Inference of interaction probability and interaction strength for the synthetic predator-prey system with slow changes in environmental condition. Dashed lines indicate true values and solid lines indicate inferred values. Transparent ribbons indicate 95% confidence intervals.

#### Intermediate environmental change

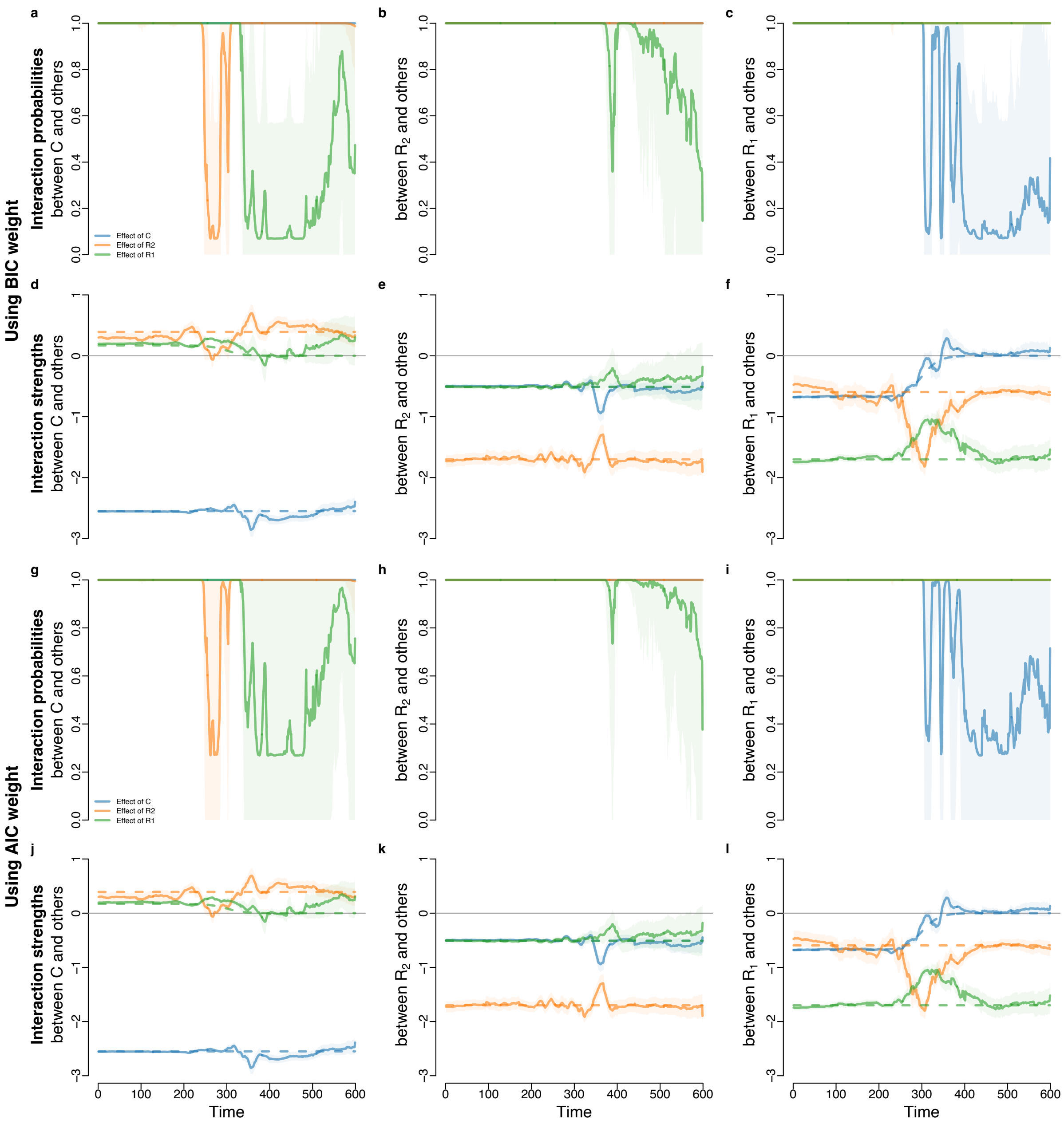

**Supplementary Fig S20.** Inference of interaction probability and interaction strength for the synthetic predator-prey system with intermediately gradual changes in environmental condition. Dashed lines indicate true values and solid lines indicate inferred values. Transparent ribbons indicate 95% confidence intervals.

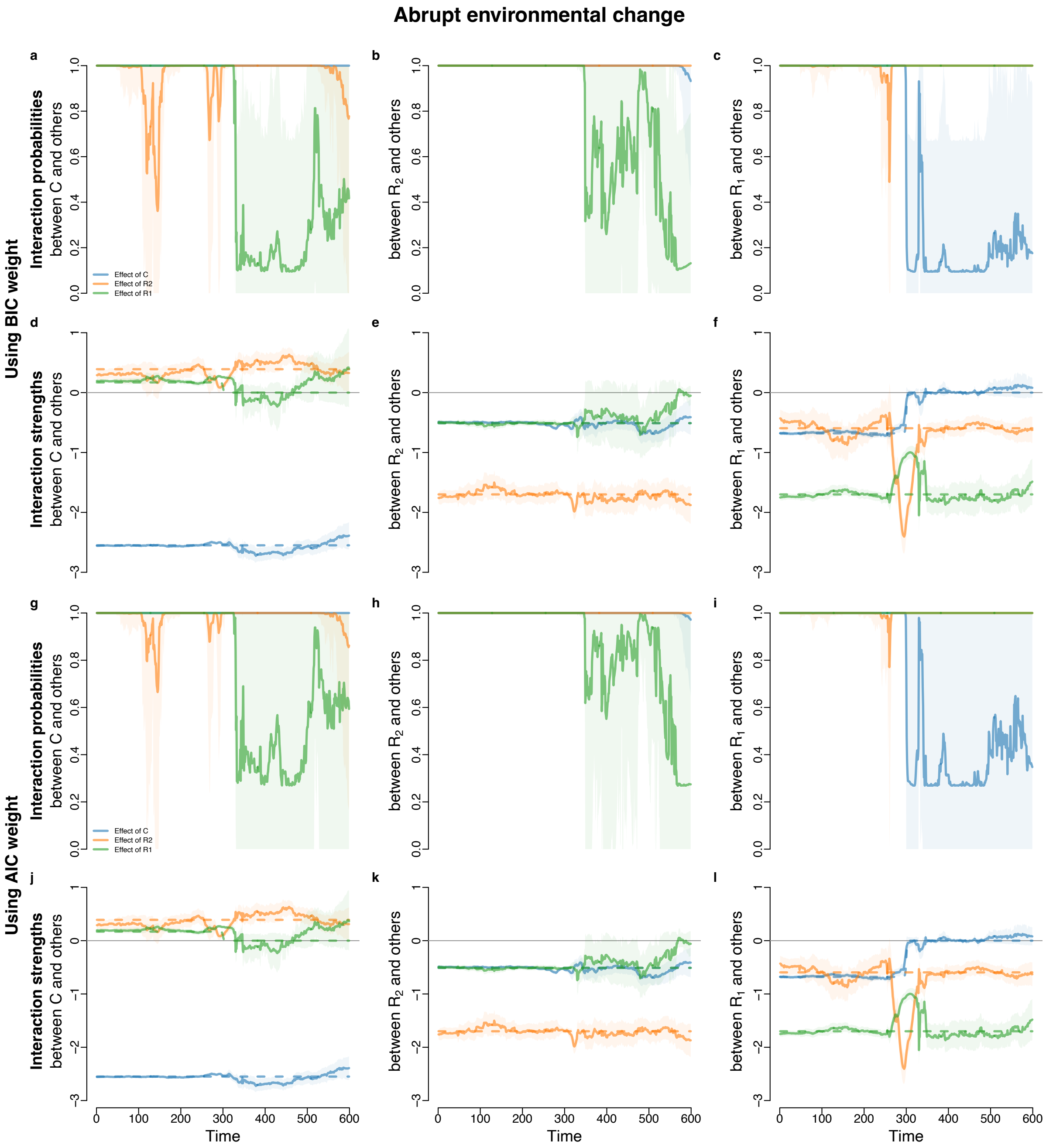

**Supplementary Fig S21.** Inference of interaction probability and interaction strengths for the synthetic predator-prey system with abrupt changes in environmental condition. Dashed lines indicate true values and solid lines indicate inferred values. Transparent ribbons indicate 95% confidence intervals.

#### S5.5 Microcosm data

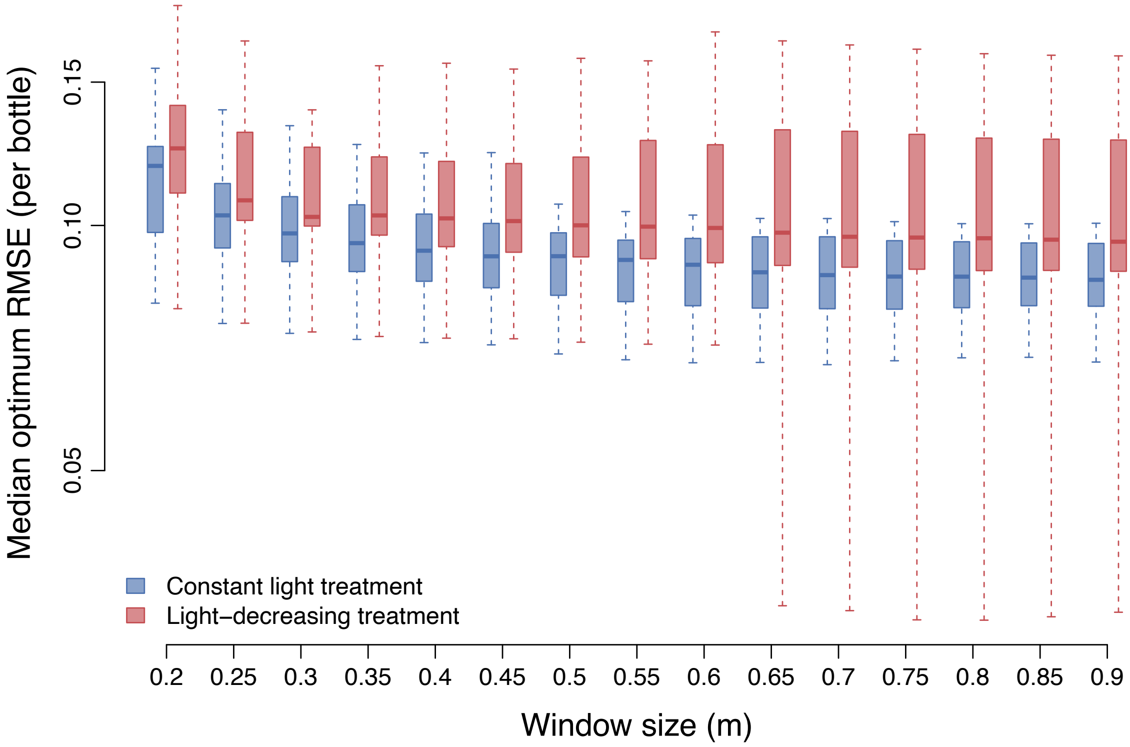

**Supplementary Fig S22.** Comparison of median RMSE for bottles under constant light treatment (blue boxes) versus median RMSE for bottles under light-decreasing treatment (red boxes).

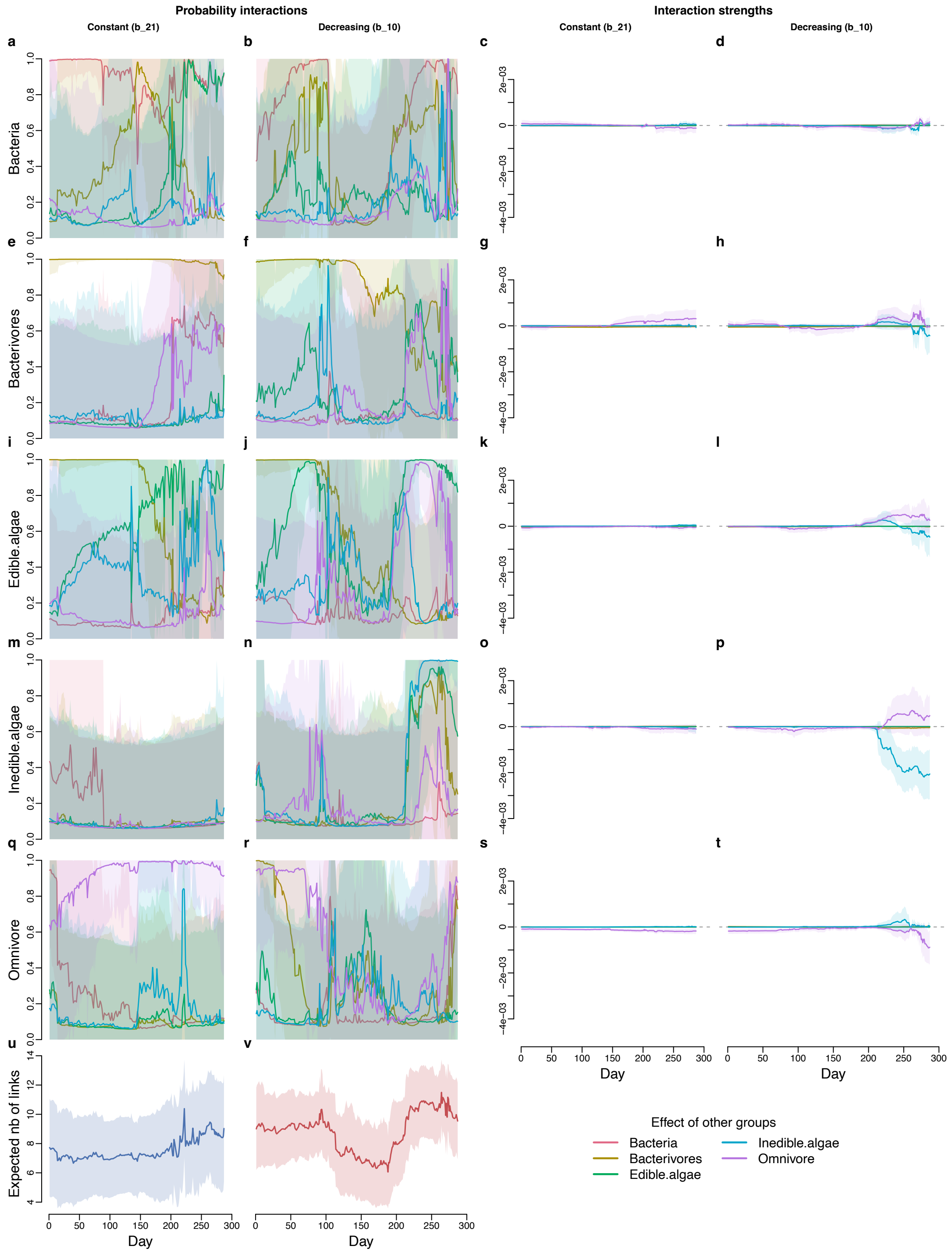

**Supplementary Fig S23.** Inferred interaction probabilities and interaction strengths for bottle 21 and bottle 10 using the chosen window size. These results complement those in figure 4. **a − d** Effects of other groups on bacteria. **e − h** Effects of other groups on bacterivores. **i − l** Effects of other groups on edible algae. **m − p** Effects of other groups on inedible algae. **q − t** Effects of other groups on omnivore. **u** Expected number of links for bottle 21 (constant light treatment). **v** Expected number of links for bottle 10 (light-decreasing treatment).

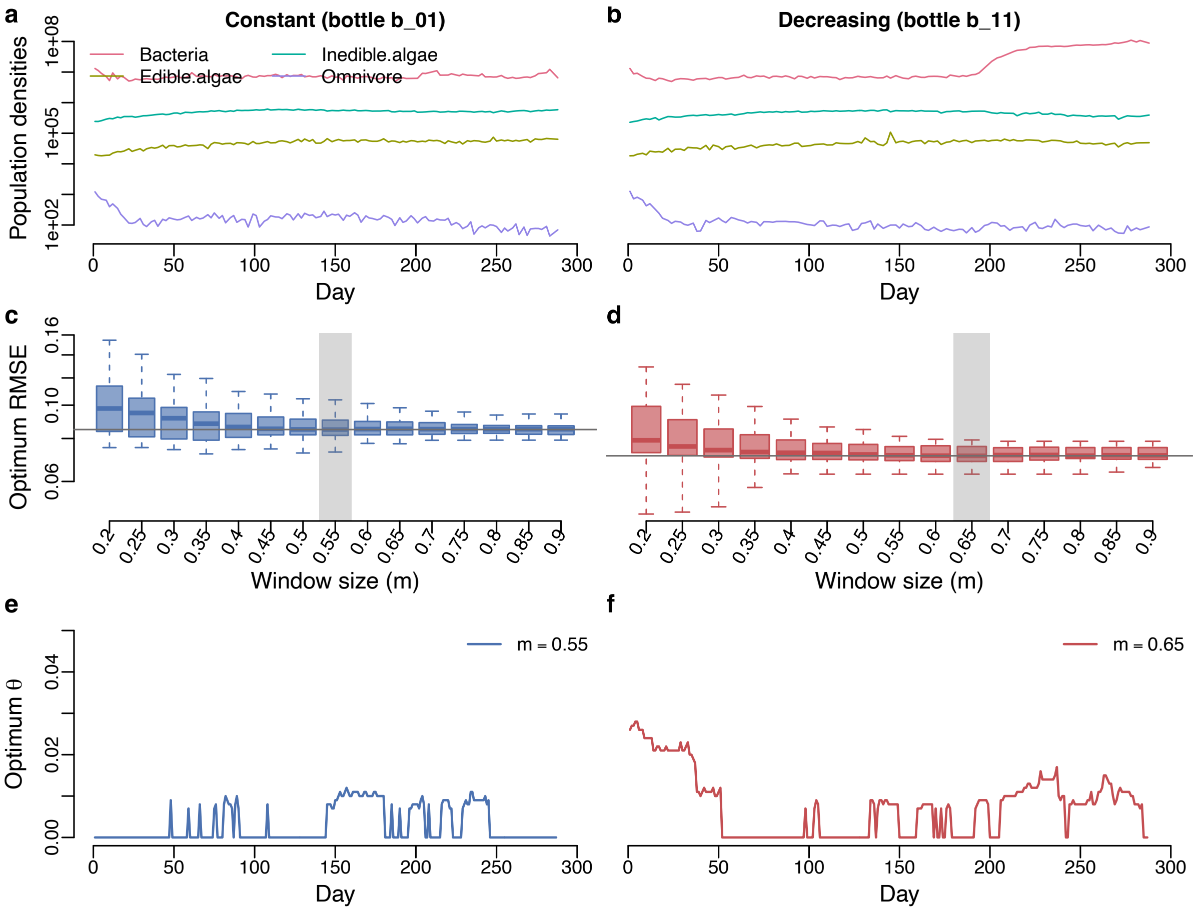

**Supplementary Fig S24.** Results for optimum RMSE and optimum $\theta$ for the microcosm experiment comparing constant light treatment (bottle 01) versus light-decreasing treatment (bottle 11) (both bottles were inoculated with the same community of 10 species). **a, b** Population densities. **c, d**. Optimum RMSE as a function of window size (m). **e, f** Time series of optimum $\theta$ that corresponds to the chosen window size (which is marked by the transparent gray rectangle).

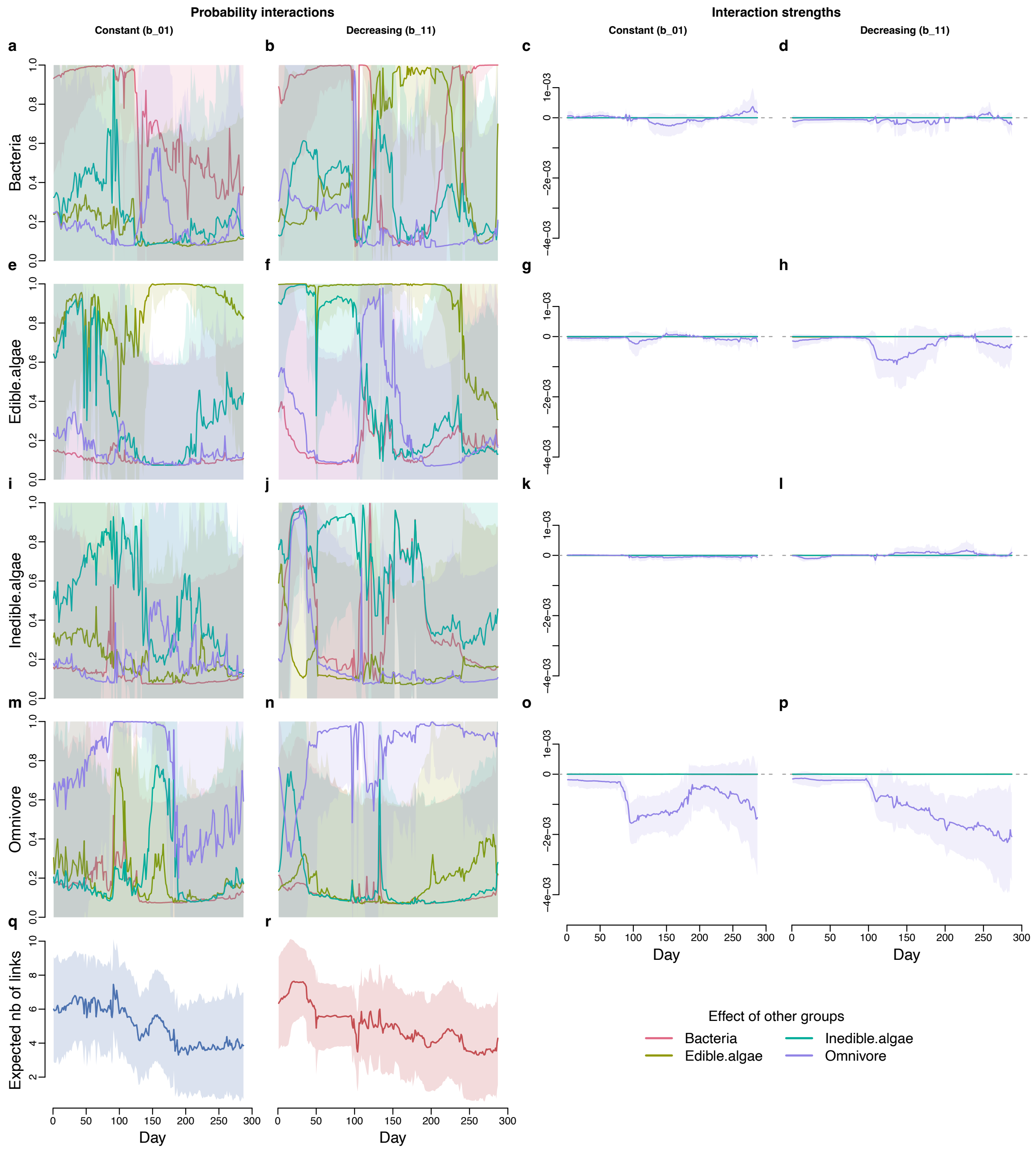

**Supplementary Fig S25.** Inferred interaction probabilities and interaction strengths using the chosen window size for the microcosm experiment comparing constant light treatment (bottle 01) versus light-decreasing treatment (bottle 11) (both bottles include the same community of 10 species). **a** – **d** Effects of other groups on bacteria. **e** – **h** Effects of other groups on edible algae. **i** – **l** Effects of other groups on inedible algae. **m** – **p** Effects of other groups on omnivore. **q** Expected number of links for constant light treatment bottle. **r** Expected number of links for light-decreasing treatment bottle.

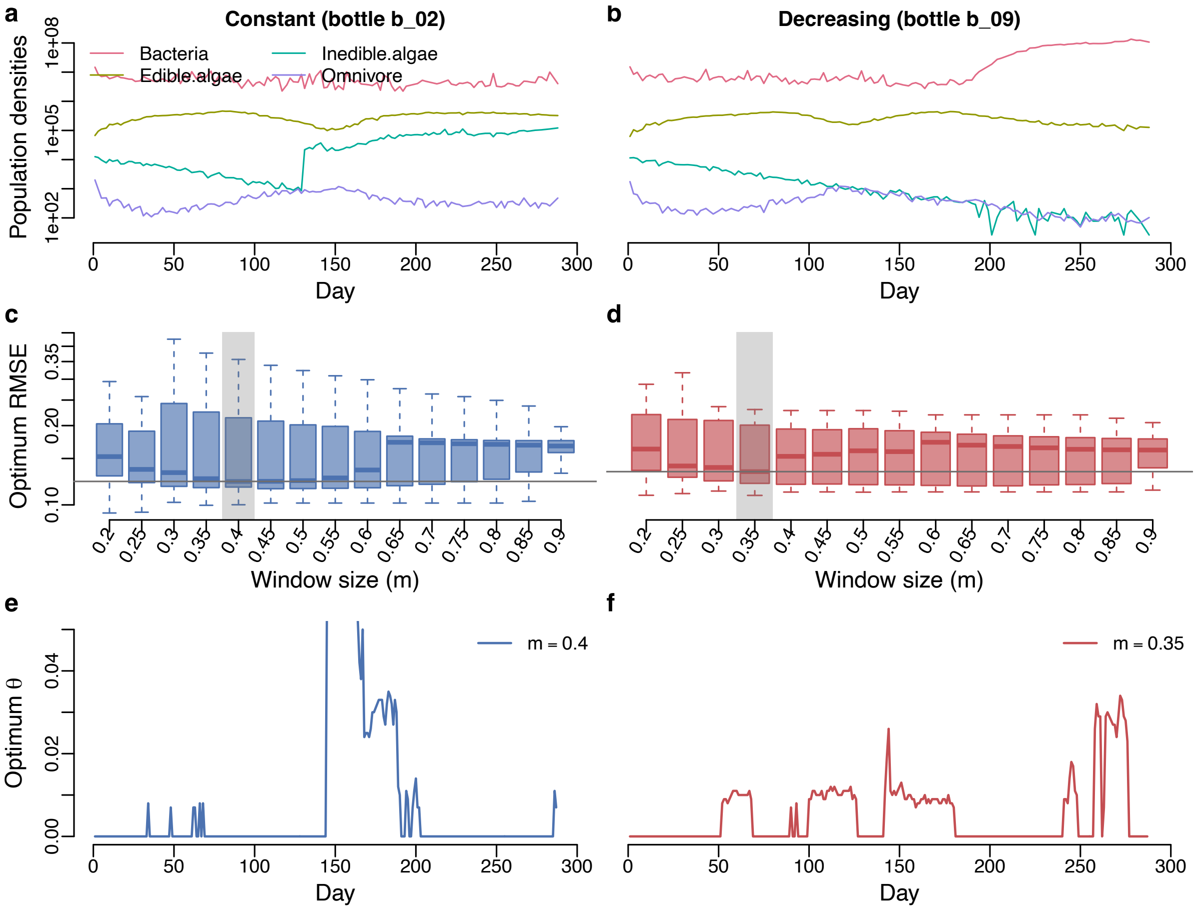

**Supplementary Fig S26.** Results for optimum RMSE and optimum $\theta$ for the microcosm experiment comparing constant light treatment (bottle 02) versus light-decreasing treatment (bottle 09) (both bottles were inoculated with the same community of 7 species). **a, b** Population densities. **c, d**. Optimum RMSE as a function of window size (m). **e, f** Time series of optimum $\theta$ that corresponds to the chosen window size (which is marked by the transparent gray rectangle).

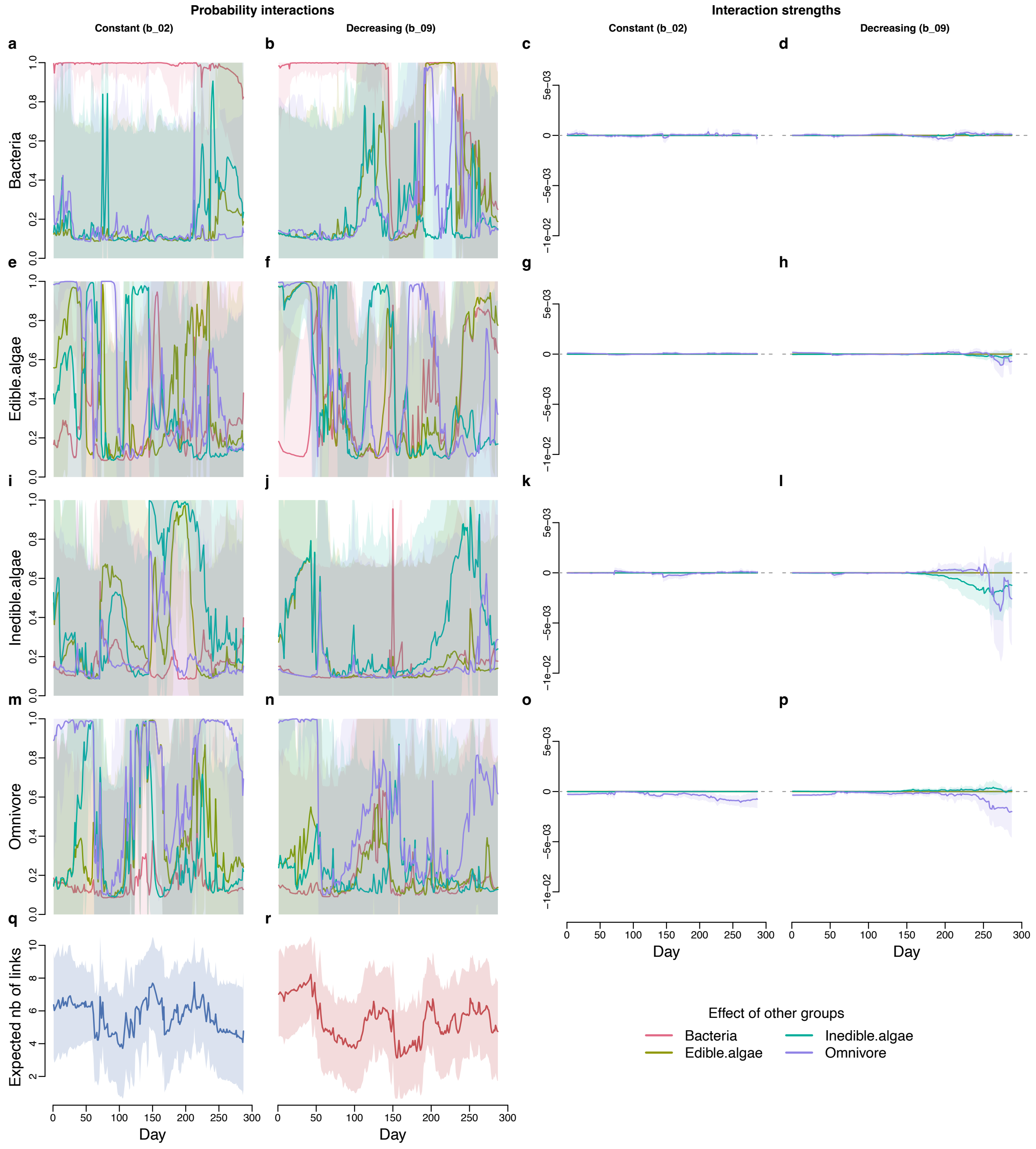

**Supplementary Fig S27.** Inferred interaction probabilities and interaction strengths using the chosen window size for the microcosm experiment comparing constant light treatment (bottle 02) versus light-decreasing treatment (bottle 09) (both bottles include the same community of 7 species). **a** – **d** Effects of other groups on bacteria. **e** – **h** Effects of other groups on edible algae. **i** – **l** Effects of other groups on inedible algae. **m** – **p** Effects of other groups on omnivore. **q** Expected number of links for constant light treatment. **r** Expected number of links for light-decreasing treatment.

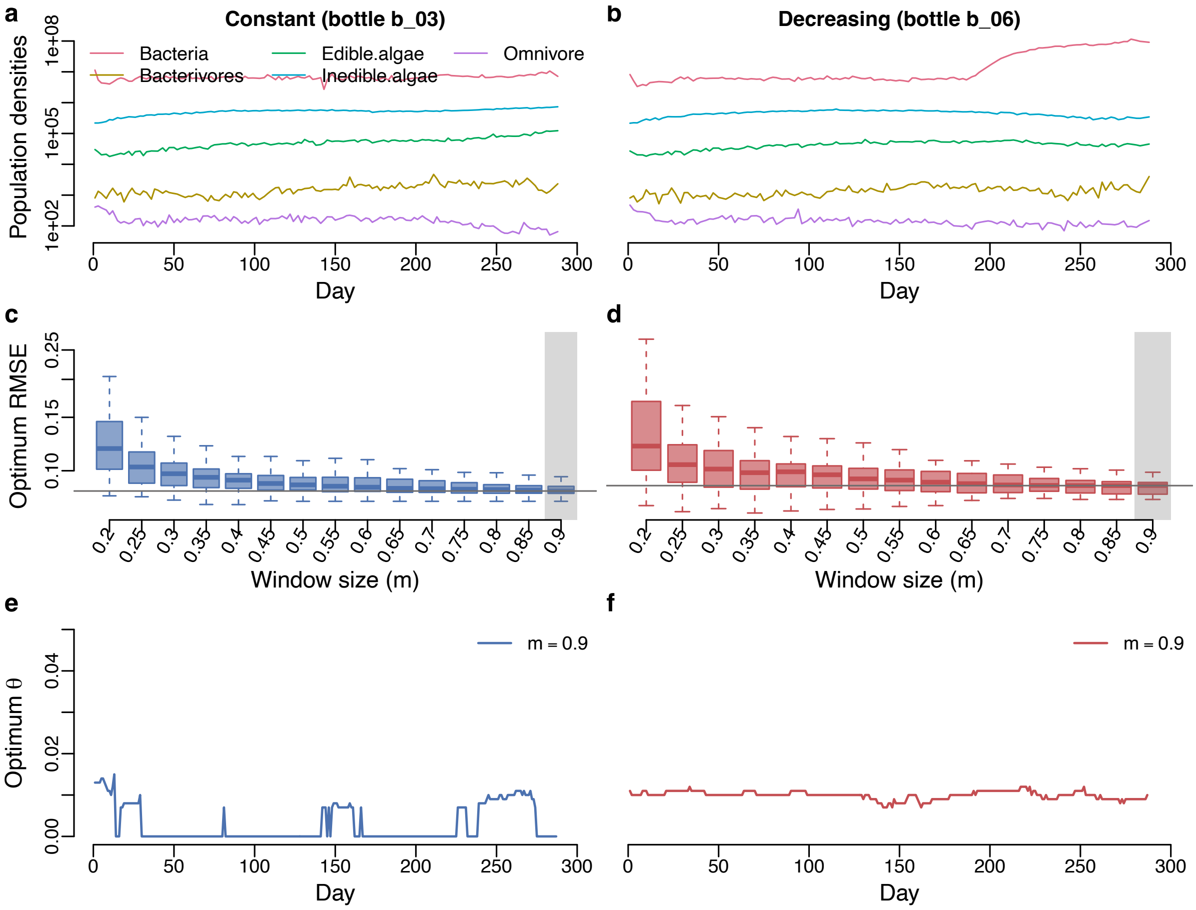

**Supplementary Fig S28.** Results for optimum RMSE and optimum $\theta$ for the microcosm experiment comparing constant light treatment (bottle 03) versus light-decreasing treatment (bottle 06) (both bottles were inoculated with the same community of 14 species). **a, b** Population densities. **c, d**. Optimum RMSE as a function of window size (m). **e, f** Time series of optimum $\theta$ that corresponds to the chosen window size (which is marked by the transparent gray rectangle).

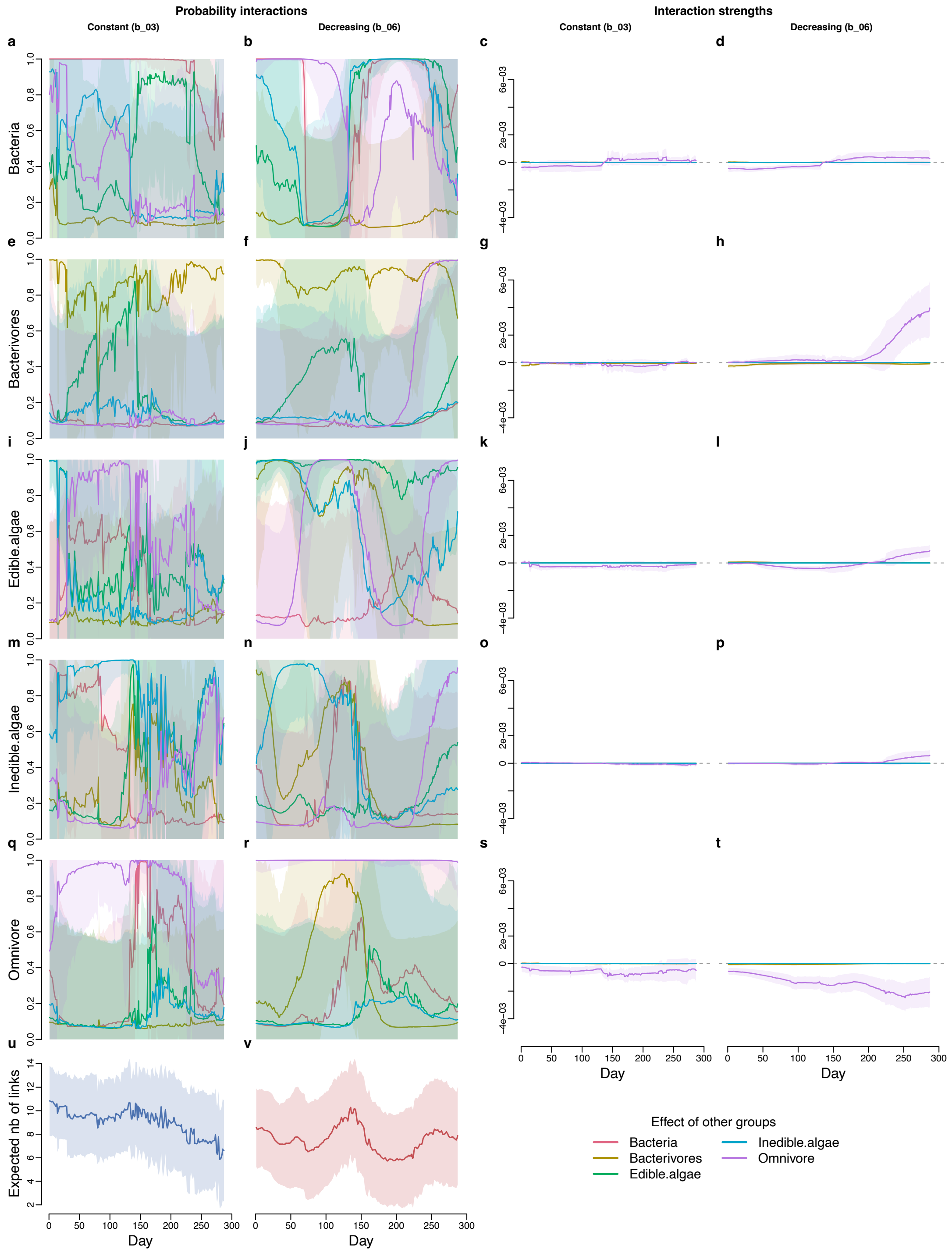

**Supplementary Fig S29.** Inferred interaction probabilities and interaction strengths using the chosen window size for the microcosm experiment comparing constant light treatment (bottle 03) versus light-decreasing treatment (bottle 06) (both bottles were inoculated with the same community of 14 species). **a–d** Effects of other groups on bacteria. **e–h** Effects of other groups on bacterivores. **i –l** Effects of other groups on edible algae. **m–p** Effects of other groups on inedible algae. **q–t** Effects of other groups on omnivore. **u** Expected number of links for constant light treatment. **v** Expected number of links for light-decreasing treatment.

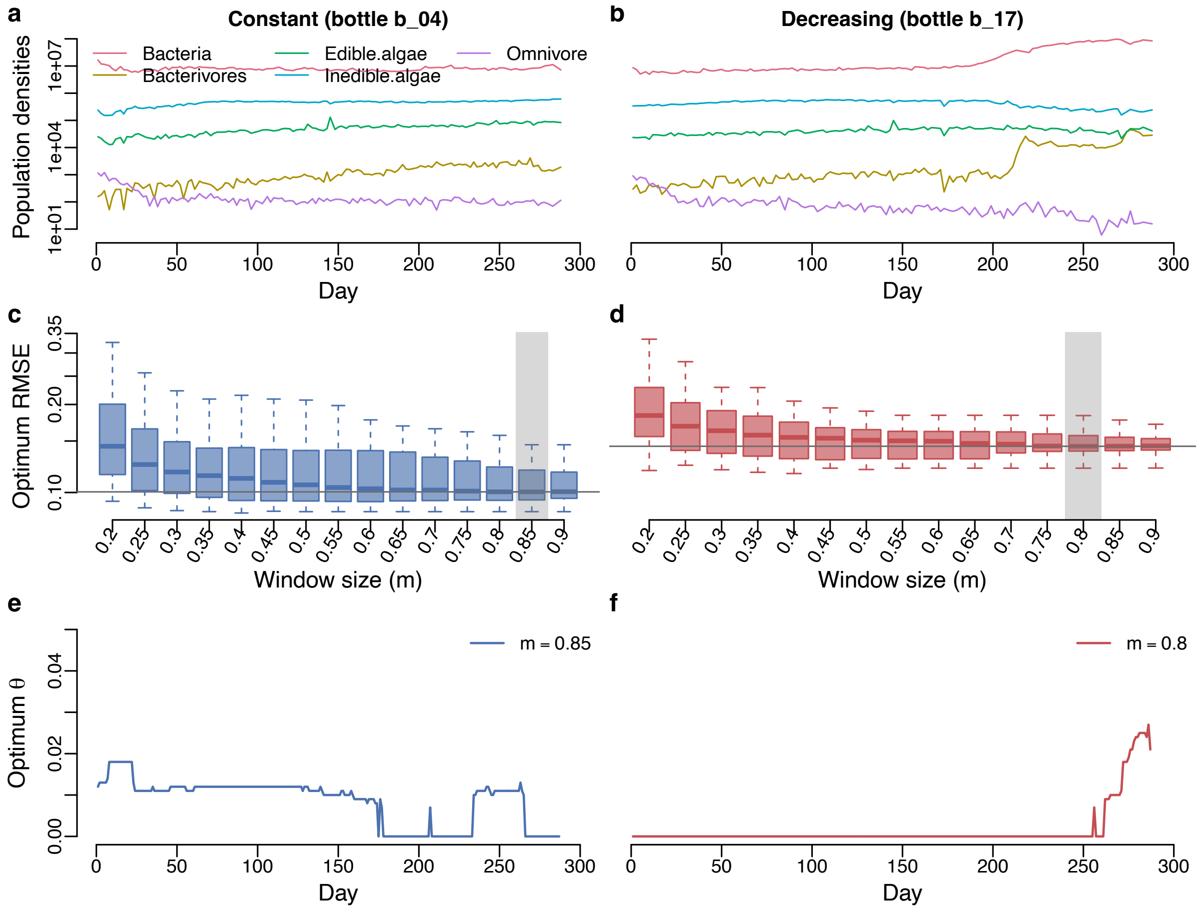

**Supplementary Fig S30.** Results for optimum RMSE and optimum $\theta$ for the microcosm experiment comparing constant light treatment (bottle 04) versus light-decreasing treatment (bottle 17) (both bottles were inoculated with the same community of 7 species). **a, b** Population densities. **c, d**. Optimum RMSE as a function of window size (m). **e, f** Time series of optimum $\theta$ that corresponds to the chosen window size (which is marked by the transparent gray rectangle).

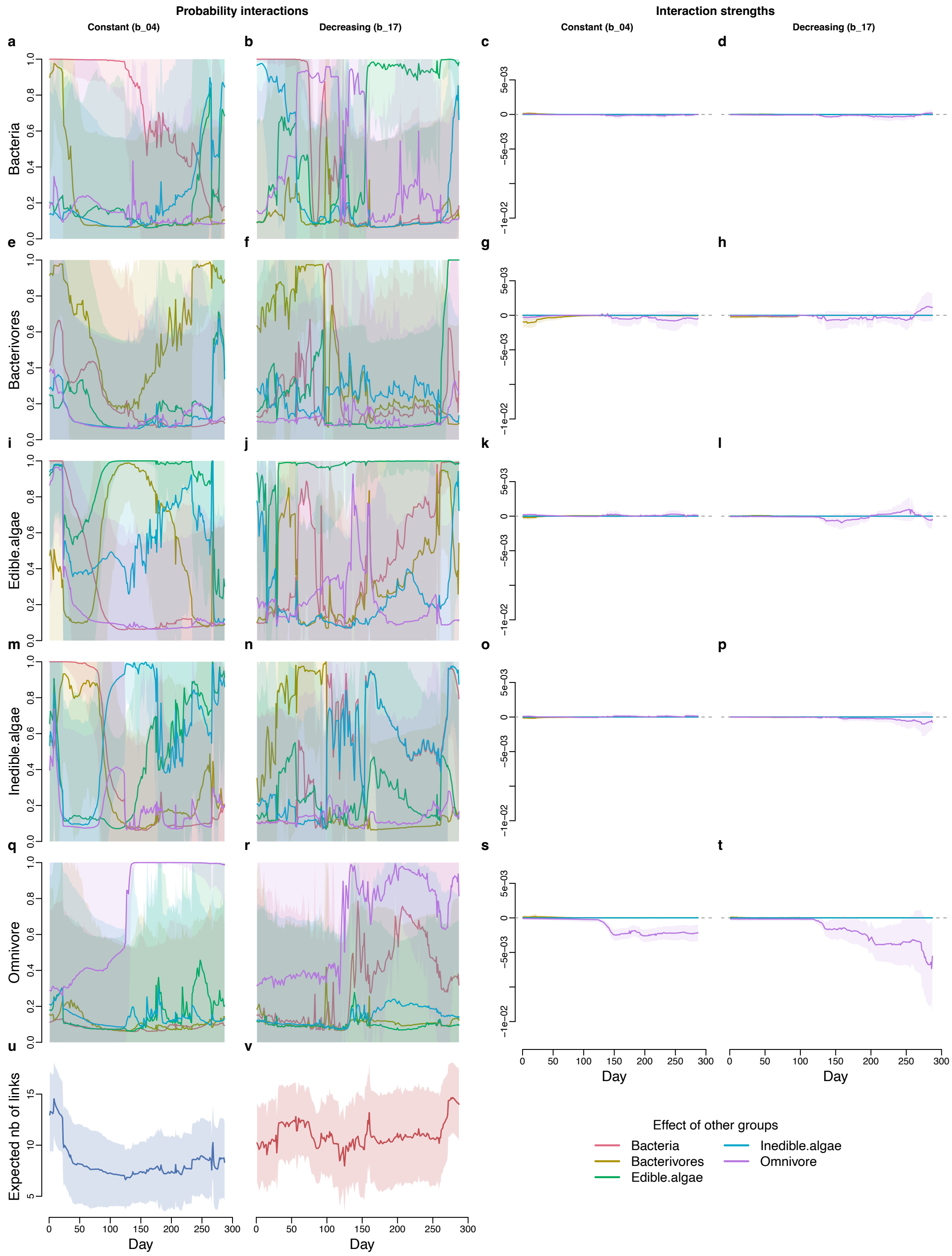

**Supplementary Fig S31.** Inferred interaction probabilities and interaction strengths using the chosen window size for the microcosm experiment comparing constant light treatment (bottle 04) versus light-decreasing treatment (bottle 17) (both bottles were inoculated with the same community of 7 species). **a–d** Effects of other groups on bacteria. **e–h** Effects of other groups on bacterivores. **i–l** Effects of other groups on edible algae. **m–p** Effects of other groups on inedible algae. **q–t** Effects of other groups on omnivore. **u** Expected number of links in the bottle with constant light treatment. **v** Expected number of links in the bottle with light-decreasing treatment.

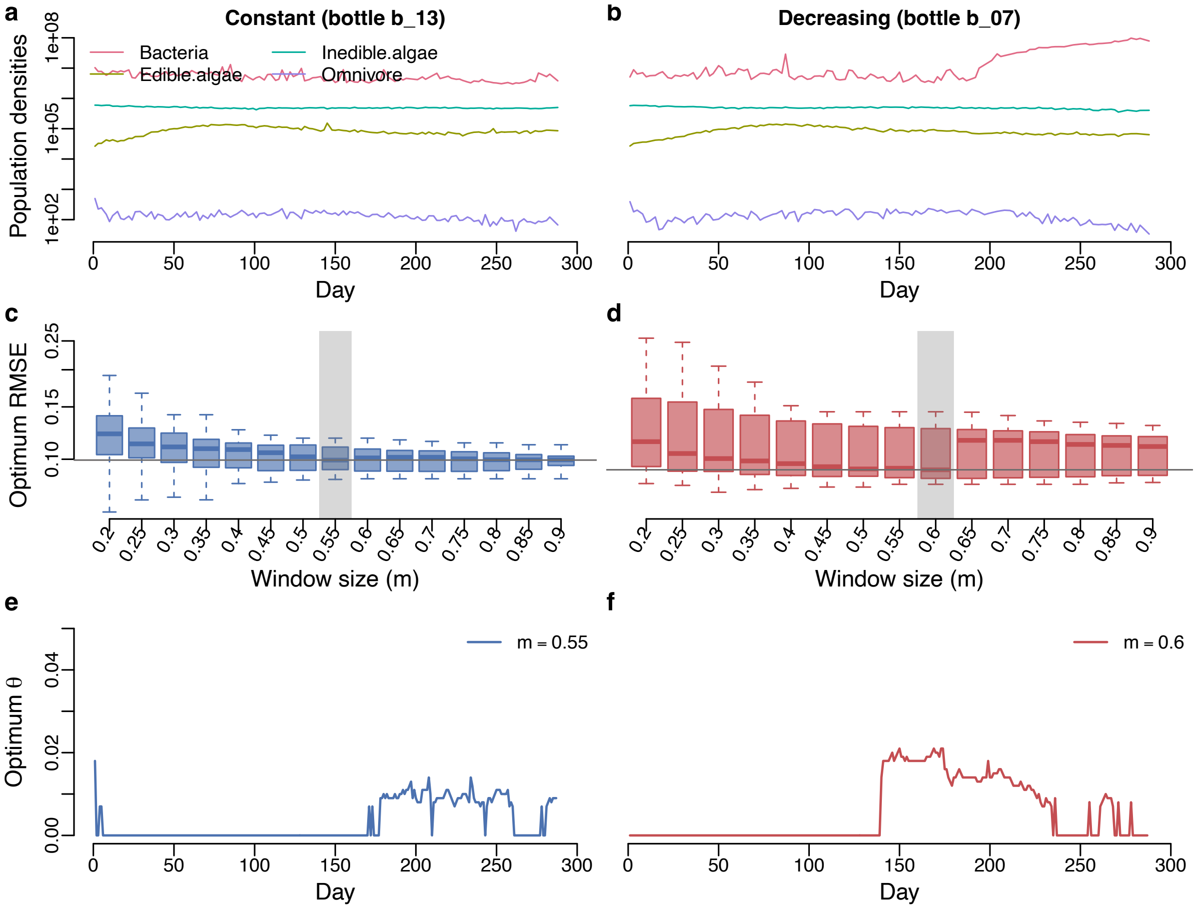

**Supplementary Fig S32.** Results for optimum RMSE and optimum $\theta$ for the microcosm experiment comparing constant light treatment (bottle 13) versus light-decreasing treatment (bottle 07) (both bottles were inoculated with the same community of 10 species). **a, b** Population densities. **c, d**. Optimum RMSE as a function of window size (m). **e, f** Time series of optimum $\theta$ that corresponds to the chosen window size (which is marked by the transparent gray rectangle).

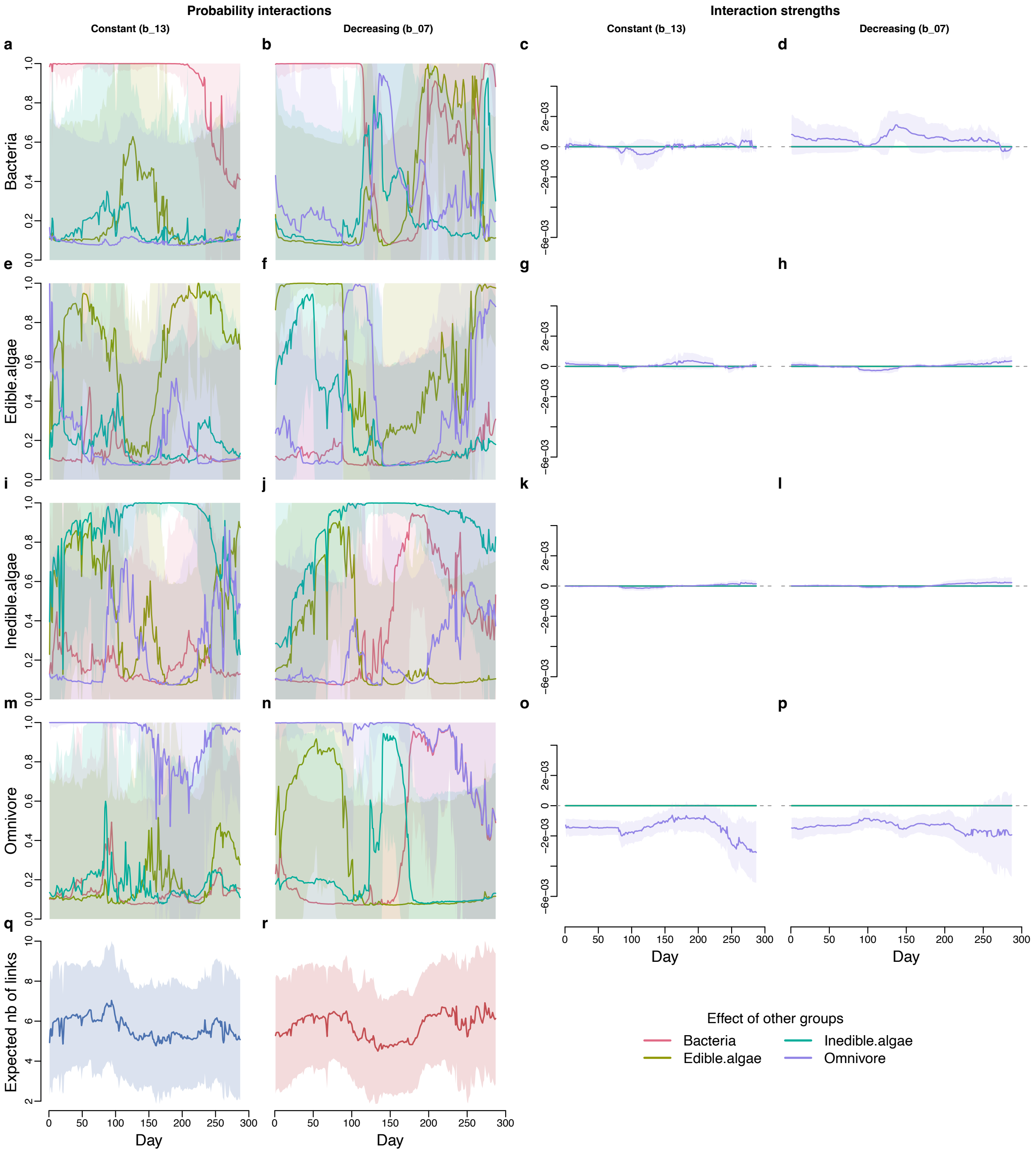

**Supplementary Fig S33.** Inferred interaction probabilities and interaction strengths using the chosen window size for the microcosm experiment comparing constant light treatment (bottle 13) versus light-decreasing treatment (bottle 07) (both bottles were inoculated with the same community of 10 species). **a–d** Effects of other functional groups on bacteria. **e–h** Effects of other functional groups on edible algae. **i–l** Effects of other functional groups on inedible algae. **m–p** Effects of other functional groups on omnivore. **q** Expected number of links in the bottle with constant light treatment. **r** Expected number of links in the bottle with light-decreasing treatment.

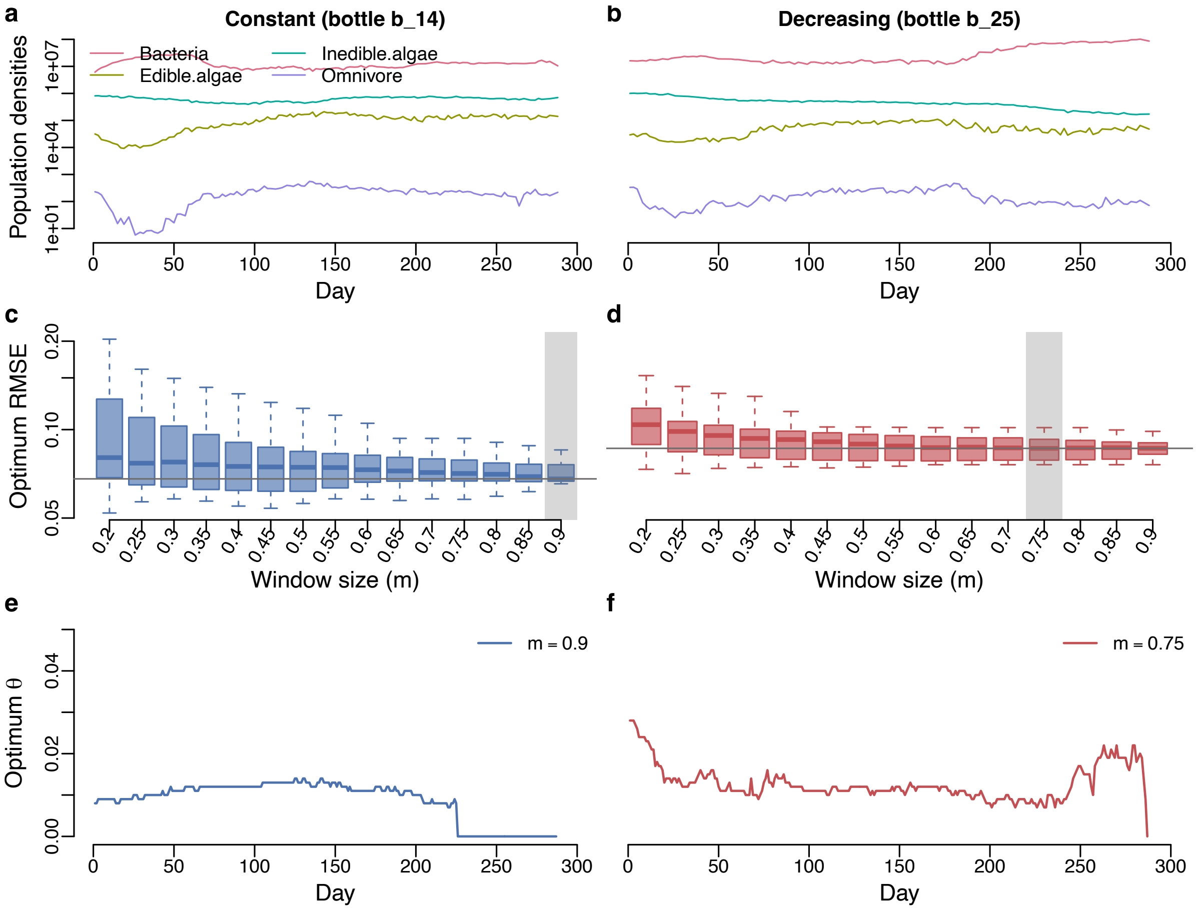

**Supplementary Fig S34.** Results for optimum RMSE and optimum $\theta$ for the microcosm experiment comparing constant light treatment (bottle 14) versus light-decreasing treatment (bottle 25) (both bottles were inoculated with the same community of 7 species). **a, b** Population densities. **c, d**. Optimum RMSE as a function of window size (m). **e, f** Time series of optimum $\theta$ that corresponds to the chosen window size (which is marked by the transparent gray rectangle).

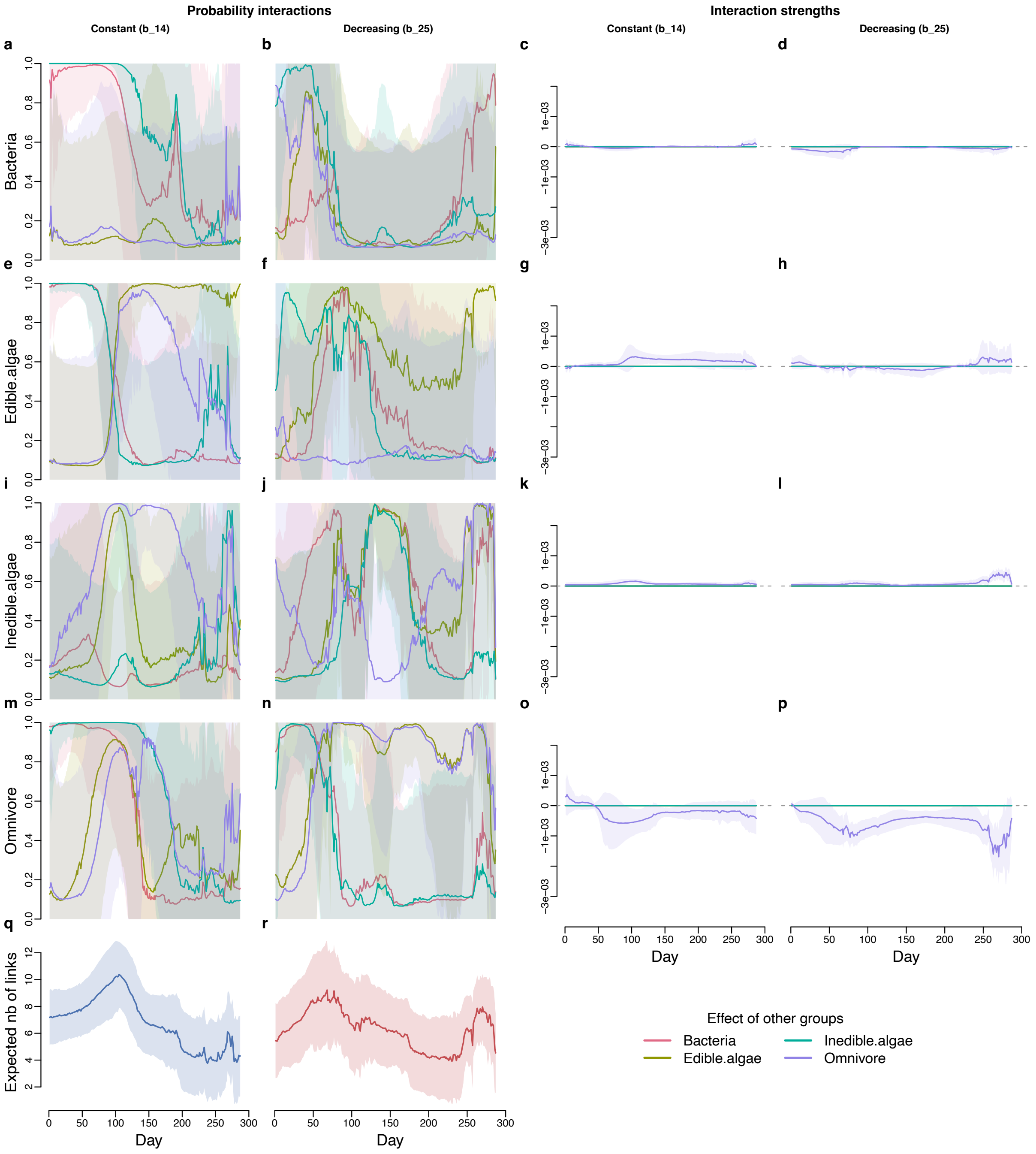

**Supplementary Fig S35.** Inferred interaction probabilities and interaction strengths using the chosen window size for the microcosm experiment comparing constant light treatment (bottle 14) versus light-decreasing treatment (bottle 25) (both bottles were inoculated with the same community of 7 species). **a–d** Effects of other groups on bacteria. **e–h** Effects of other groups on edible algae. **i–l** Effects of other groups on inedible algae. **m–p** Effects of other groups on omnivore. **q** Expected number of links in the bottle with constant light treatment. **r** Expected number of links in the bottle with light-decreasing treatment.

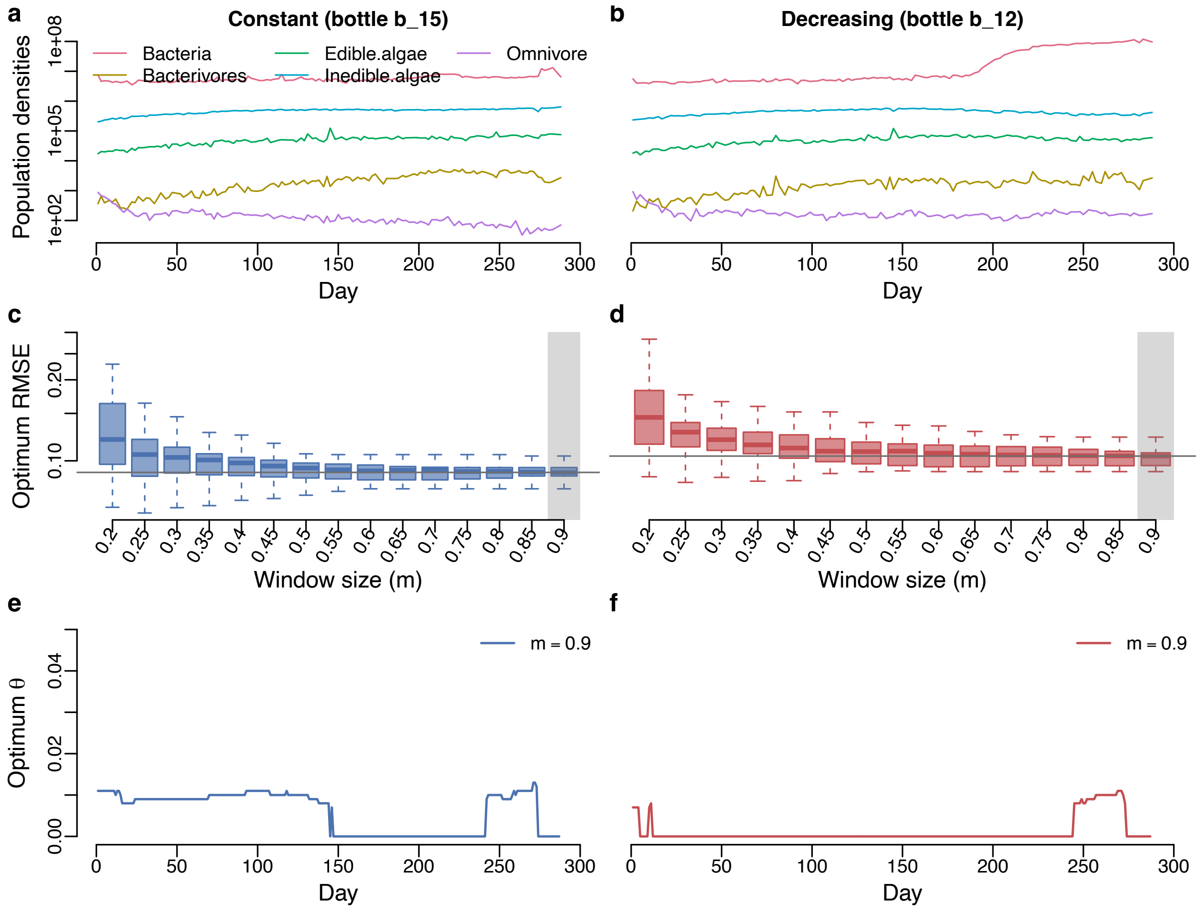

**Supplementary Fig S36.** Results for optimum RMSE and optimum $\theta$ for the microcosm experiment comparing constant light treatment (bottle 15) versus light-decreasing treatment (bottle 12) (both bottles were inoculated with the same community of 7 species). **a, b** Population densities. **c, d**. Optimum RMSE as a function of window size (m). **e, f** Time series of optimum $\theta$ that corresponds to the chosen window size (which is marked by the transparent gray rectangle).

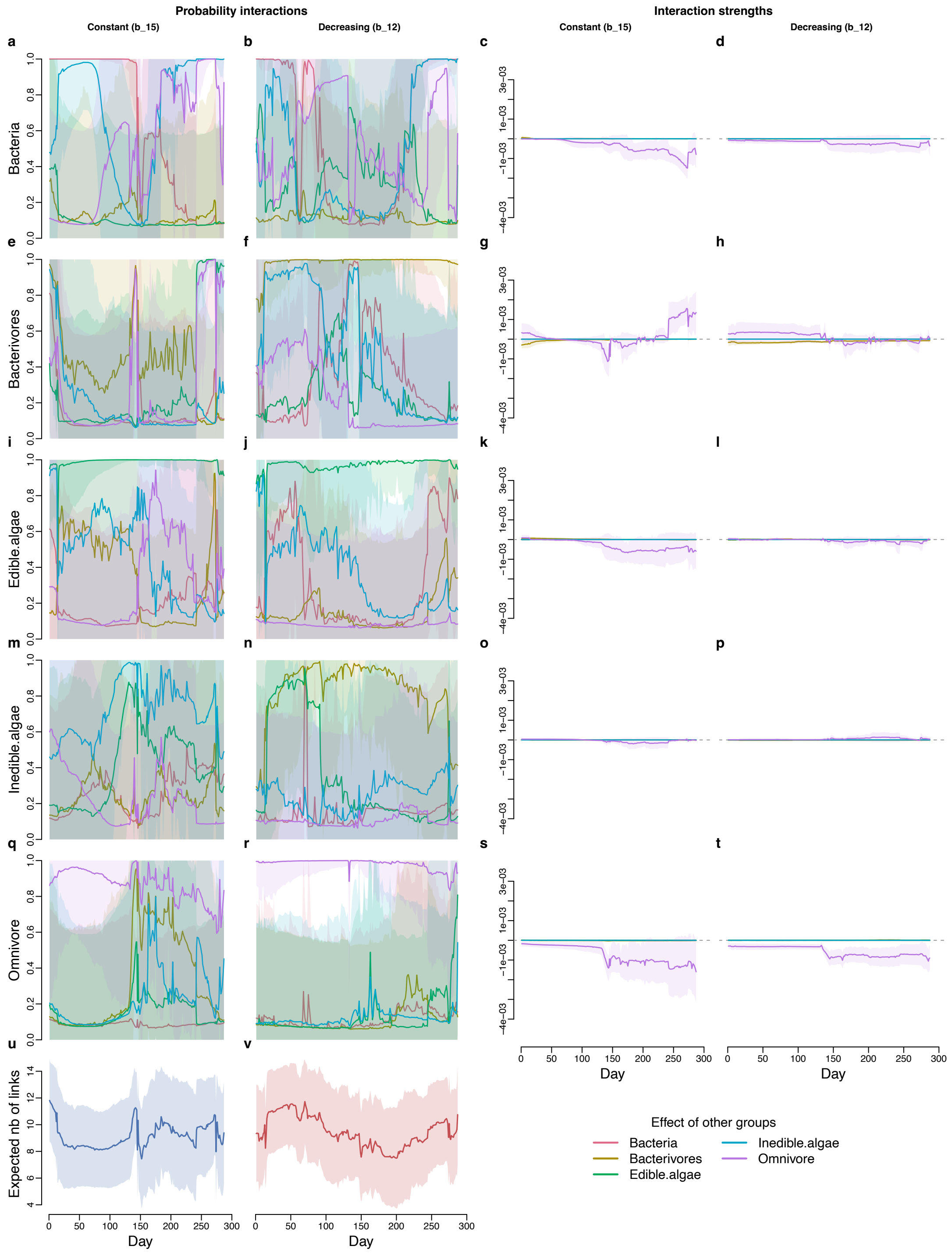

**Supplementary Fig S37.** Inferred interaction probabilities and interaction strengths using the chosen window size for the microcosm experiment comparing constant light treatment (bottle 15) versus light-decreasing treatment (bottle 12) (both bottles were inoculated with the same community of 7 species). **a–d** Effects of other groups on bacteria. **e–h** Effects of other groups on bacterivores. **i–l** Effects of other groups on edible algae. **m–p** Effects of other groups on inedible algae. **q–t** Effects of other groups on omnivore. **u** Expected number of links in the bottle with constant light treatment. **v** Expected number of links in the bottle with light-decreasing treatment.

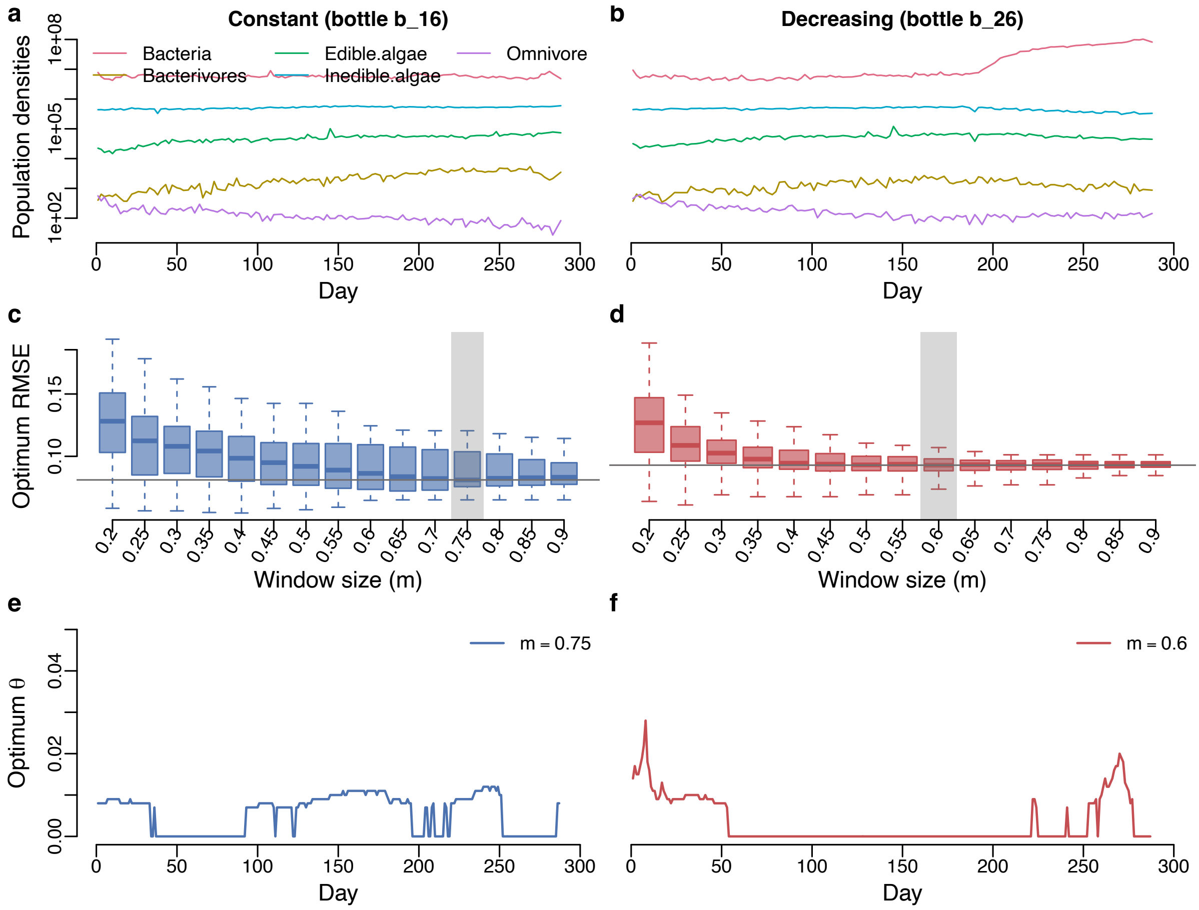

**Supplementary Fig S38.** Results for optimum RMSE and optimum $\theta$ for the microcosm experiment comparing constant light treatment (bottle 16) versus light-decreasing treatment (bottle 26) (both bottles were inoculated with the same community of 14 species). **a, b** Population densities. **c, d**. Optimum RMSE as a function of window size (m). **e, f** Time series of optimum $\theta$ that corresponds to the chosen window size (which is marked by the transparent gray rectangle).

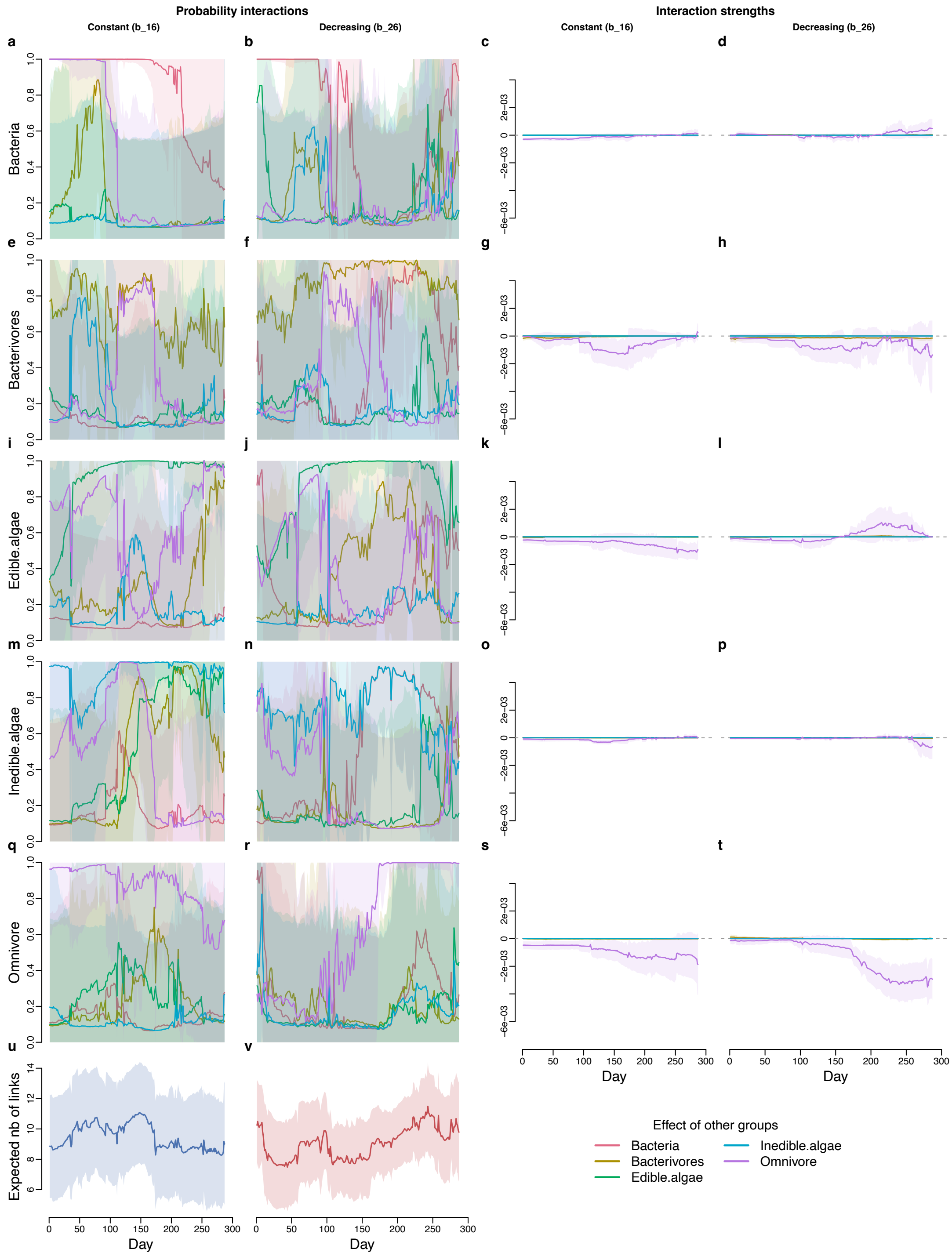

**Supplementary Fig S39.** Inferred interaction probabilities and interaction strengths using the chosen window size for the microcosm experiment comparing constant light treatment (bottle 16) versus light-decreasing treatment (bottle 26) (both bottles were inoculated with the same community of 14 species). **a–d**. Effects of other groups on bacteria. **e–h**. Effects of other groups on bacterivores. **i–l**. Effects of other groups on edible algae. **m–p**. Effects of other groups on inedible algae. **q–t**. Effects of other groups on omnivore. **u** Expected number of links in the bottle with constant light treatment. **v**. Expected number of links in the bottle with light-decreasing treatment.

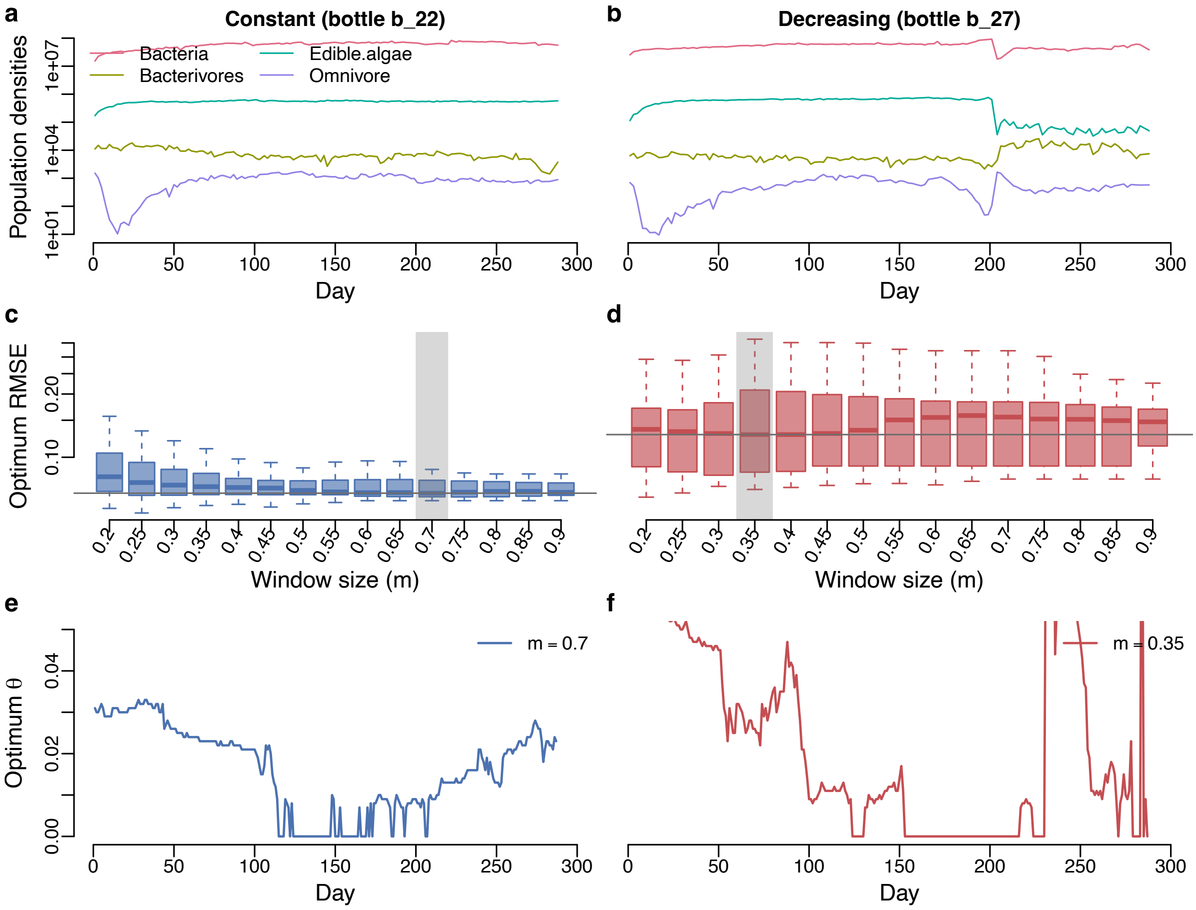

**Supplementary Fig S40.** Results for optimum RMSE and optimum $\theta$ for the microcosm experiment comparing constant light treatment (bottle 22) versus light-decreasing treatment (bottle 27) (both bottles were inoculated with the same community of 10 species). **a, b**. Population densities. **c, d**. Optimum RMSE as a function of window size (m). **e, f**. Time series of optimum $\theta$ that corresponds to the chosen window size (which is marked by the transparent gray rectangle).

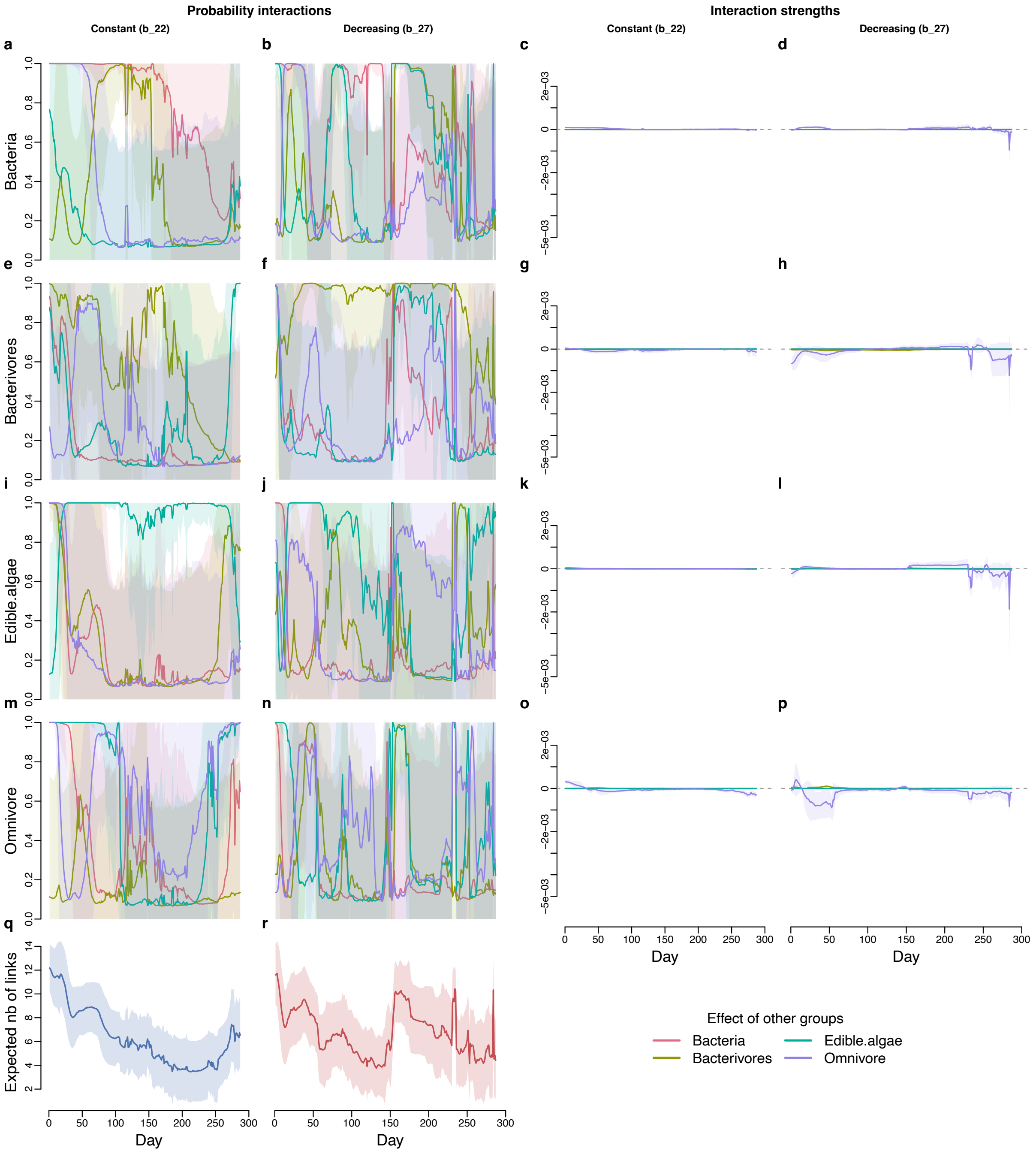

**Supplementary Fig S41.** Inferred interaction probabilities and interaction strengths using the chosen window size for the microcosm experiment comparing constant light treatment (bottle 22) versus light-decreasing treatment (bottle 27) (both bottles were inoculated with the same community of 10 species). **a–d**. Effects of other groups on bacteria. **e–h**. Effects of other groups on bacterivores. **i–l**. Effects of other groups on edible algae. **m–p**. Effects of other groups on omnivore. **q**. Expected number of links in the bottle with constant light treatment. **r**. Expected number of links in the bottle with light-decreasing treatment.

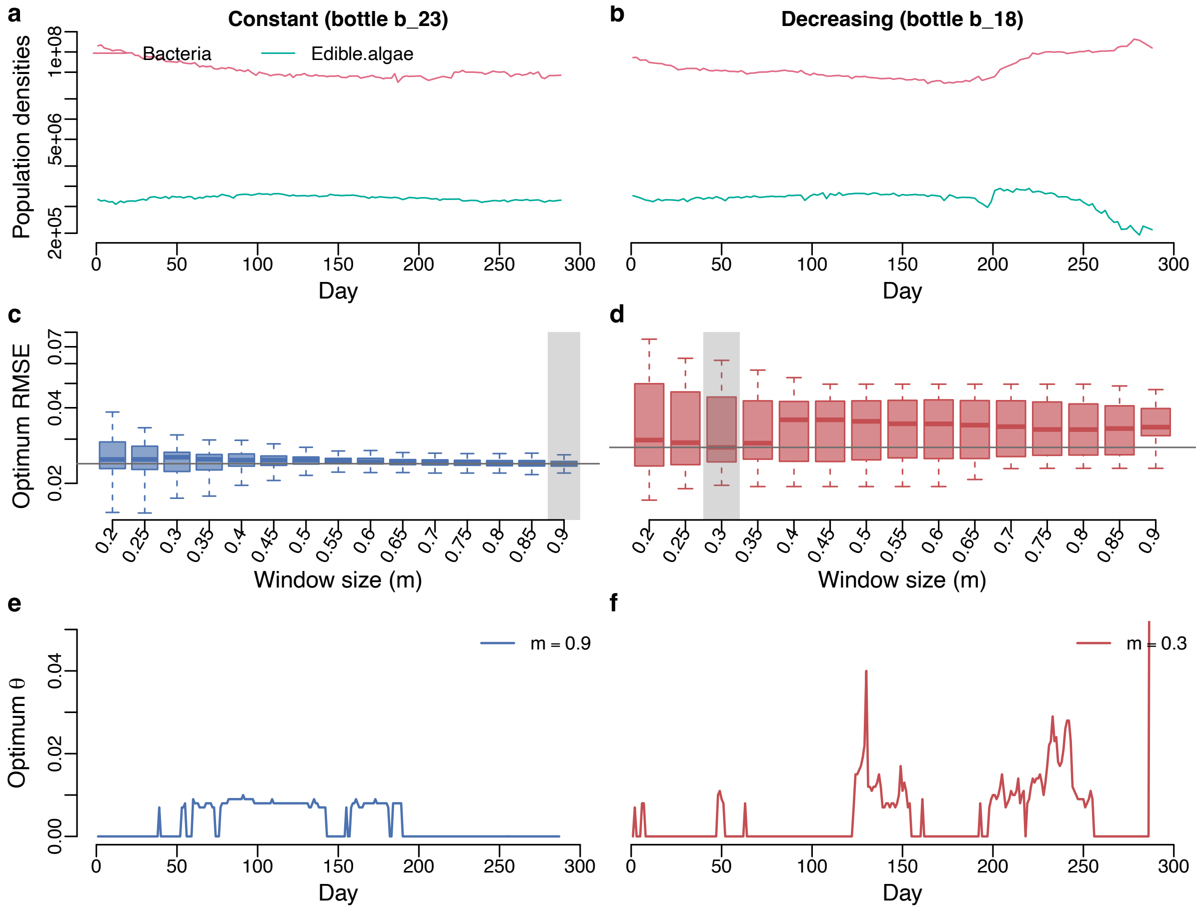

**Supplementary Fig S42.** Results for optimum RMSE and optimum $\theta$ for the microcosm experiment comparing constant light treatment (bottle 23) versus light-decreasing treatment (bottle 18) (both bottles were inoculated with the same community of 7 species). **a, b**. Population densities. **c, d**. Optimum RMSE as a function of window size (m). **e, f**. Time series of optimum $\theta$ that corresponds to the chosen window size (which is marked by the transparent gray rectangle).

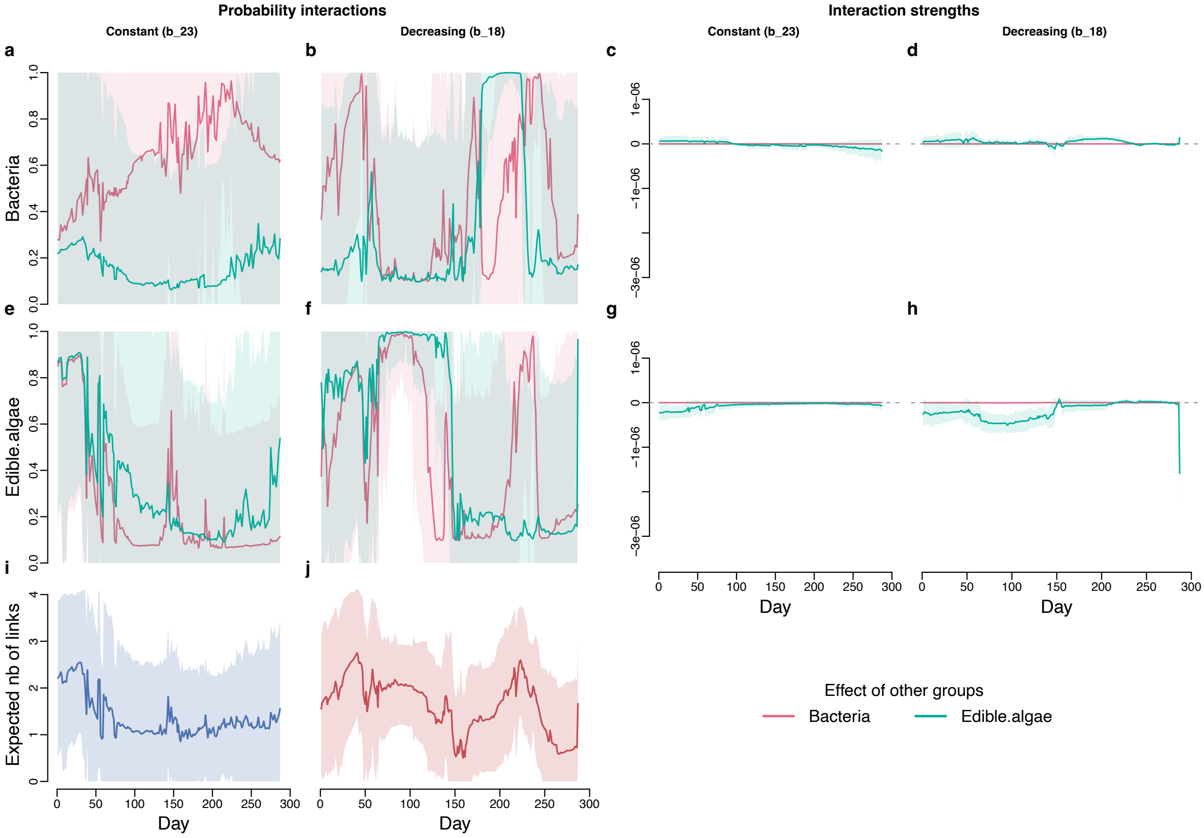

**Supplementary Fig S43.** Inferred interaction probabilities and interaction strengths using the chosen window size for the microcosm experiment comparing constant light treatment (bottle 23) versus light-decreasing treatment (bottle 18) (both bottles were inoculated with the same community of 7 species). **a–d**. Effects of edible algae and bacteria on bacteria. **e–h**. Effect of edible algae and bacteria on edible algae. **i**. Expected number of links in the bottle with constant light treatment. **j**. Expected number of links in the bottle with light-decreasing treatment.

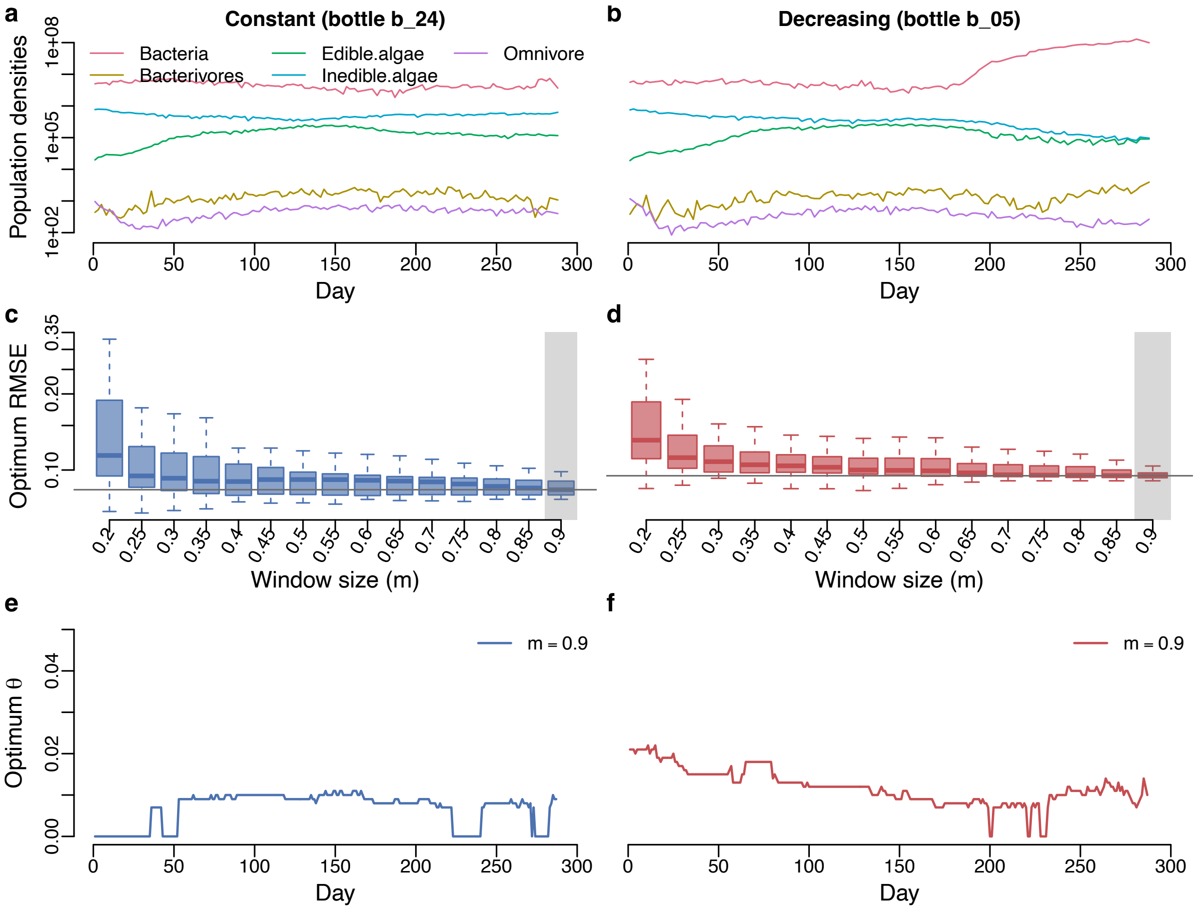

**Supplementary Fig S44.** Results for optimum RMSE and optimum $\theta$ for the microcosm experiment comparing constant light treatment (bottle 24) versus light-decreasing treatment (bottle 05) (both bottles were inoculated with the same community of 14 species). **a, b**. Population densities. **c, d**. Optimum RMSE as a function of window size (m). **e, f**. Time series of optimum $\theta$ that corresponds to the chosen window size (which is marked by the transparent gray rectangle).

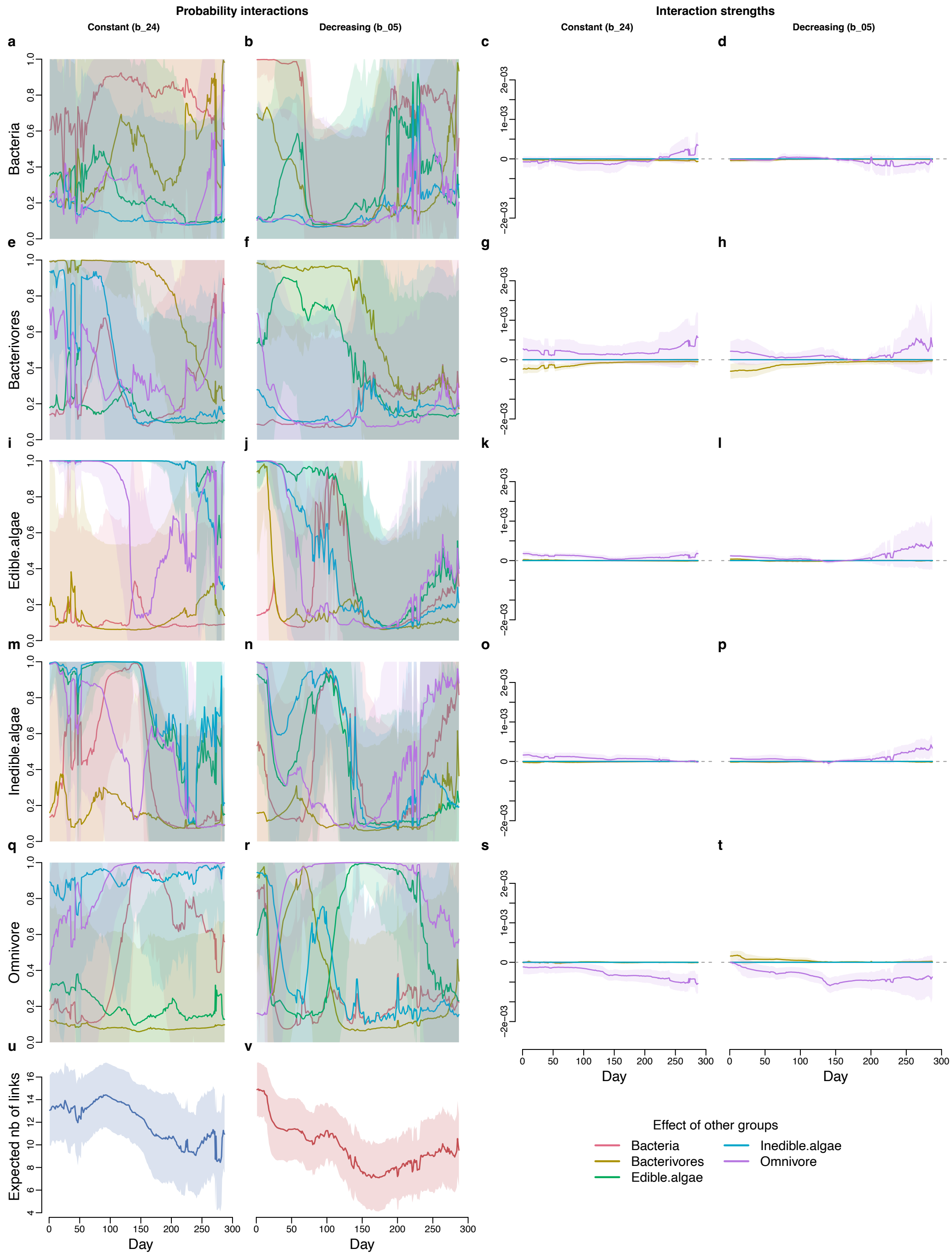

**Supplementary Fig S45.** Inferred interaction probabilities and interaction strengths using the chosen window size for the microcosm experiment comparing constant light treatment (bottle 24) versus light-decreasing treatment (bottle 05) (both bottles were inoculated with the same community composition of 14 species). **a–d**. Effects of other groups on bacteria. **e–h** Effects of other groups on bacterivores. **i–l**. Effects of other groups on edible algae. **m–p**. Effects of other groups on inedible algae. **q–t**. Effects of other groups on omnivore. **u** Expected number of links in the bottle with constant light treatment. **v** Expected number of links in the bottle with light-decreasing treatment.

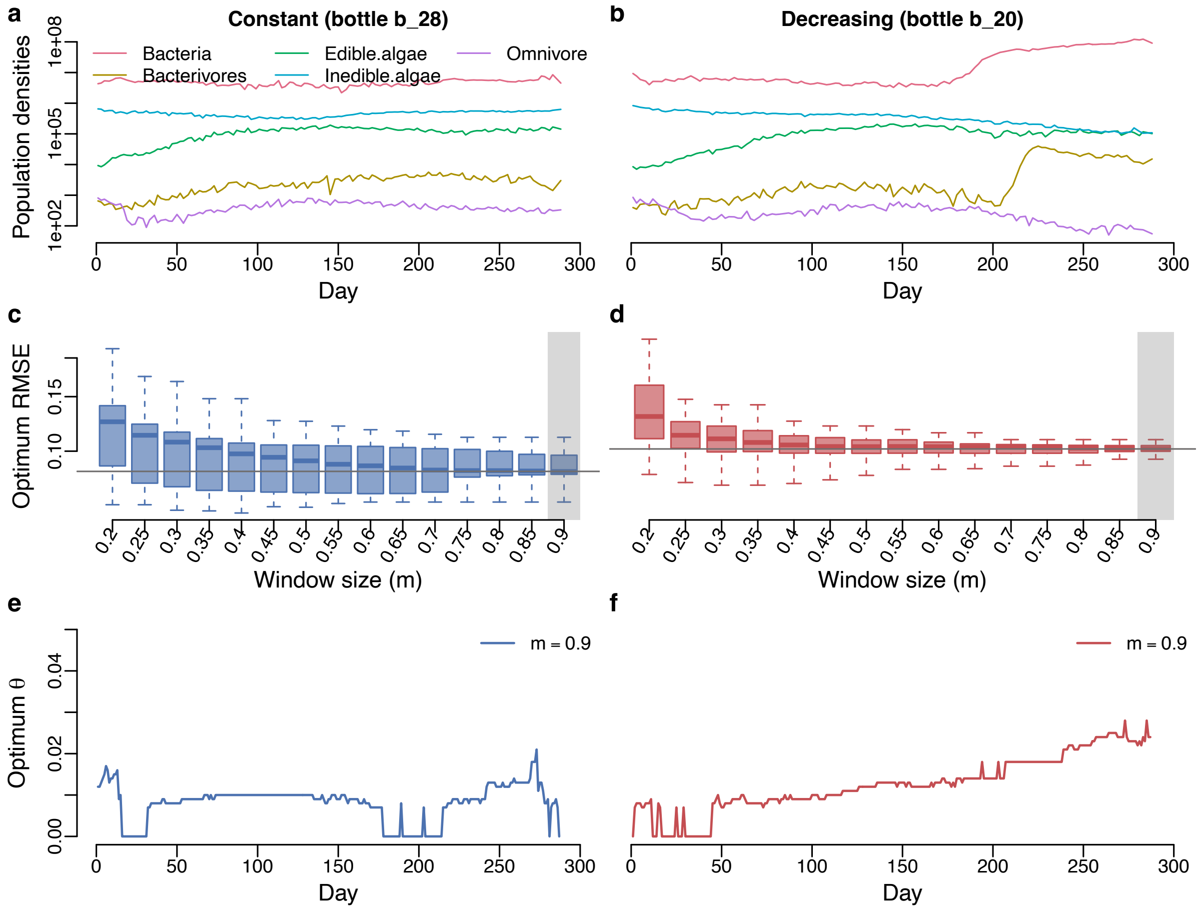

**Supplementary Fig S46.** Results for optimum RMSE and optimum $\theta$ for the microcosm experiment comparing constant light treatment (bottle 28) versus light-decreasing treatment (bottle 20) (both bottles were inoculated with the same community composition of 10 species). **a, b**. Population densities. **c, d**. Optimum RMSE as a function of window size (m). **e, f**. Time series of optimum $\theta$ that corresponds to the chosen window size (which is marked by the transparent gray rectangle).

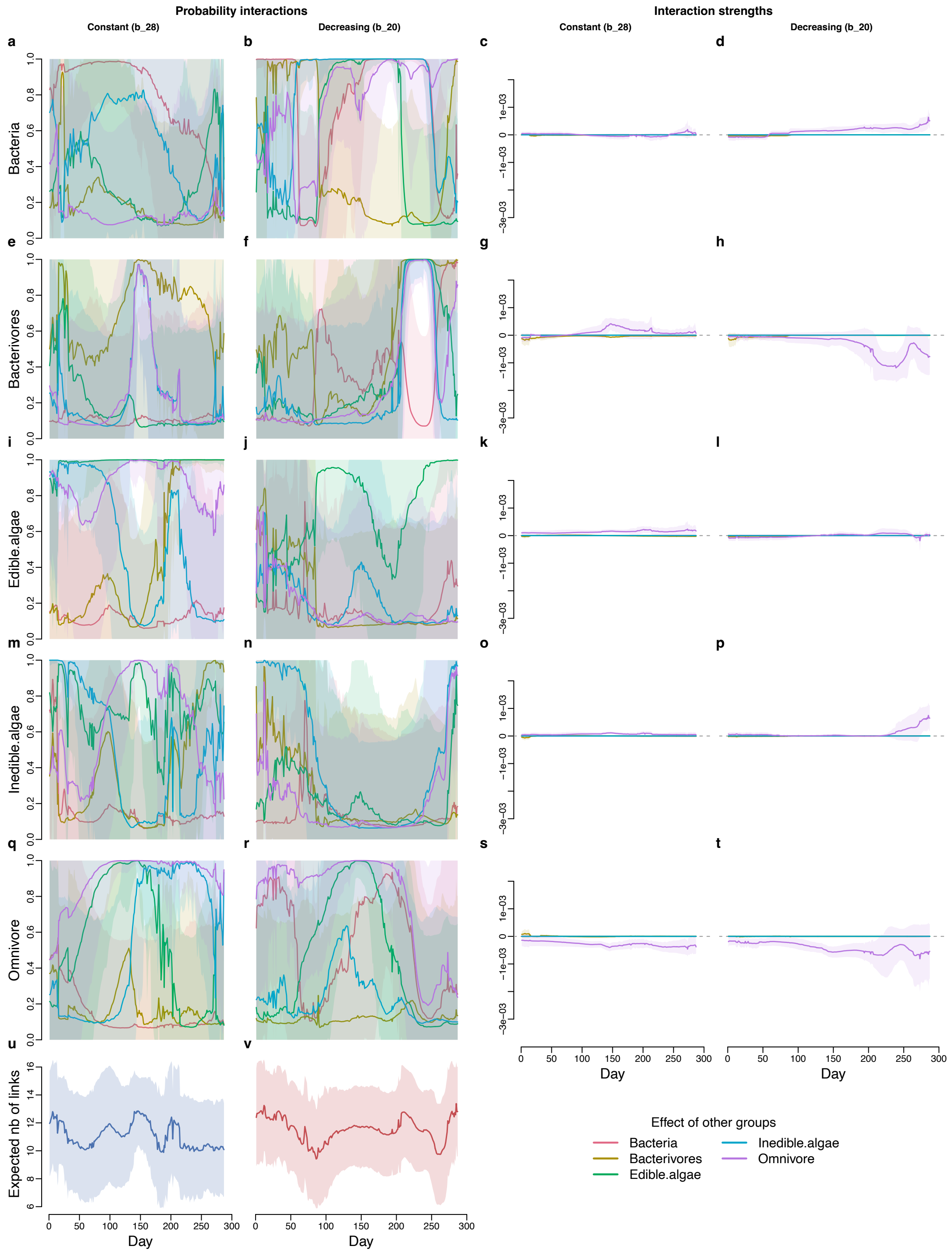

**Supplementary Fig S47.** Inferred interaction probabilities and interaction strengths using the chosen window size for the microcosm experiment comparing constant light treatment (bottle 28) versus light-decreasing treatment (bottle 20) (both bottles were inoculated with the same community composition of 10 species). **a–d**. Effect of other groups on bacteria. **e–h**. Effects of other groups on bacterivores. **i–l**. Effects of other groups on edible algae. **m–p**. Effect of other groups on inedible algae. **q–t**. Effects of other groups on omnivore. **u**. Expected number of links in the bottle with constant light treatment. **v**. Expected number of links in the bottle with light-decreasing treatment.

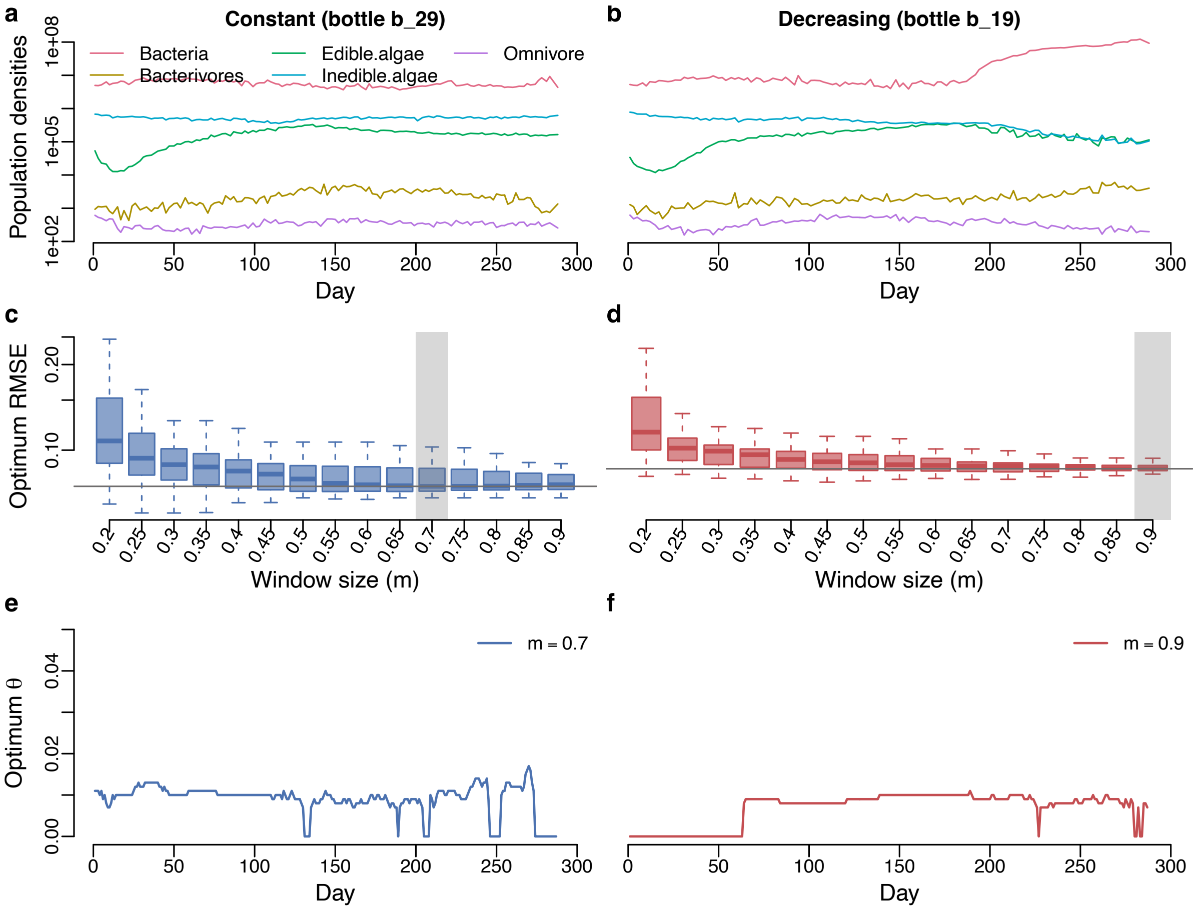

**Supplementary Fig S48.** Results for optimum RMSE and optimum $\theta$ for the microcosm experiment comparing constant light treatment (bottle 29) versus light-decreasing treatment (bottle 19) (both bottles were inoculated with the same community composition of 14 species). **a, b**. Population densities. **c, d**. Optimum RMSE as a function of window size (m). **e, f**. Time series of optimum $\theta$ that corresponds to the chosen window size (which is marked by the transparent gray rectangle).

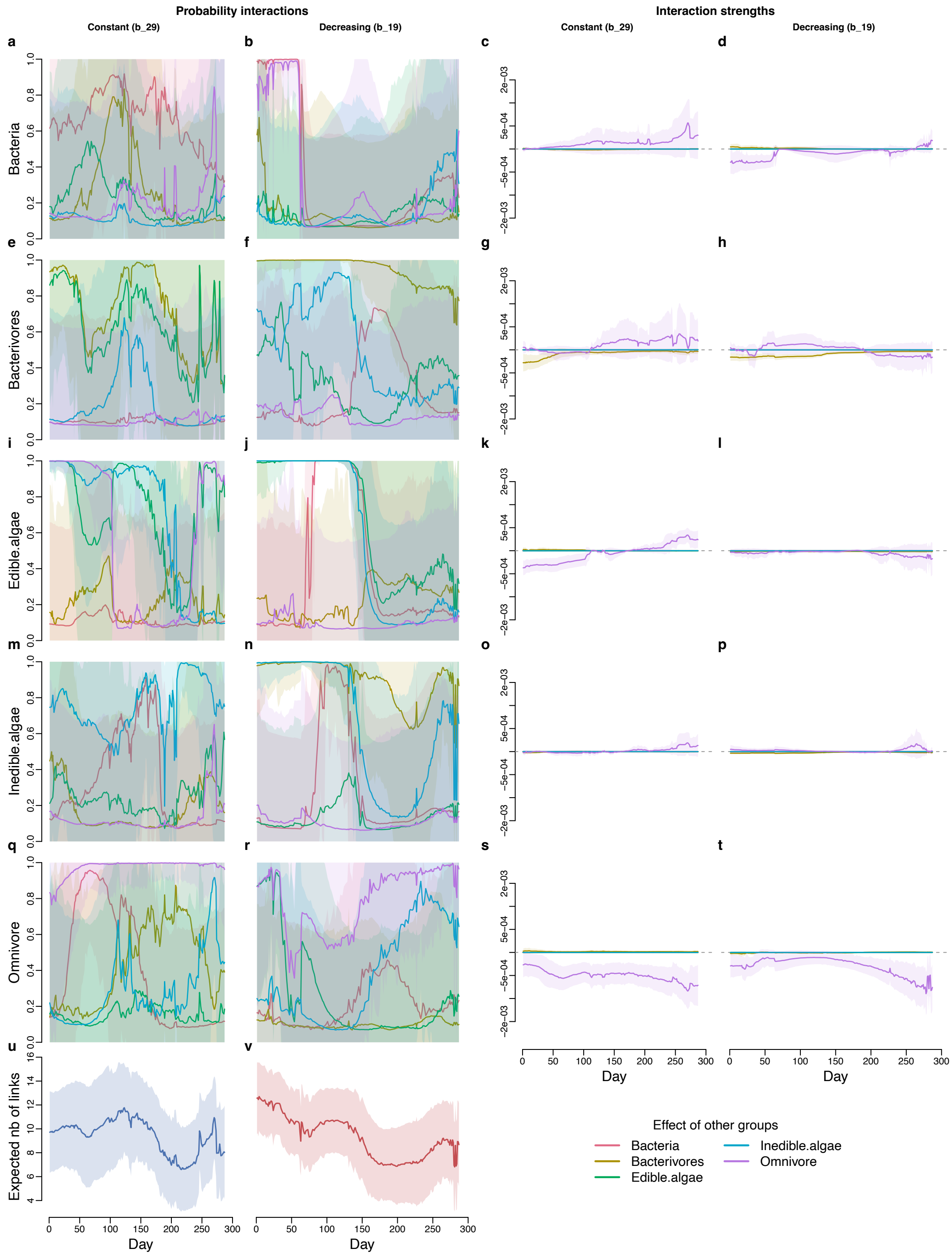

**Supplementary Fig S49.** Inferred interaction probabilities and interaction strengths using the chosen window size for the microcosm experiment comparing constant light treatment (bottle 29) versus light-decreasing treatment (bottle 19) (both bottles were inoculated with the same community composition of 14 species). **a–d**. Effects of other groups on bacteria. **e–h** Effects of other groups on bacterivores. **i–l**. Effects of other groups on edible algae. **m–p**. Effects of other groups on inedible algae. **q–t**. Effects of other groups on omnivore. **u**. Expected number of links in the bottle with constant light treatment. **v**. Expected number of links in the bottle with light-decreasing treatment.

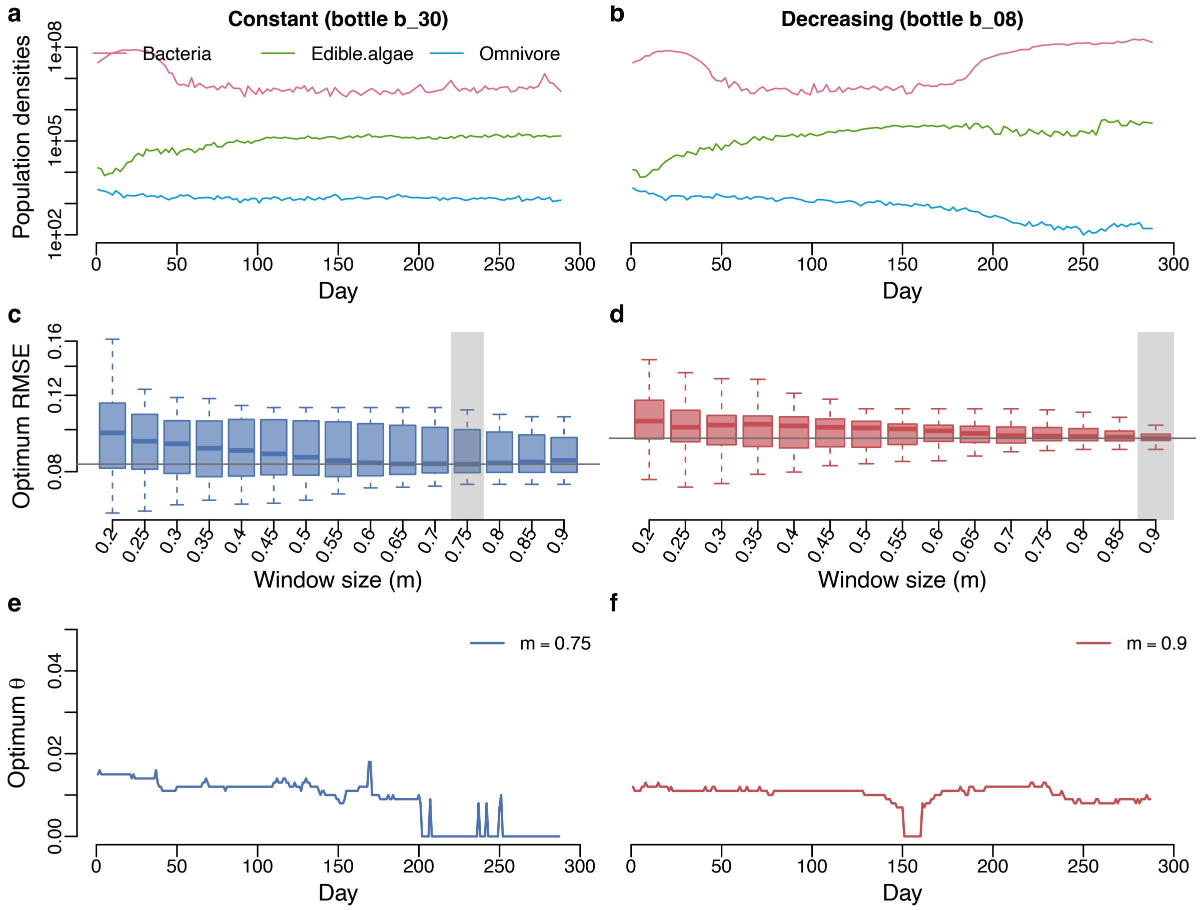

**Supplementary Fig S50.** Results for optimum RMSE and optimum $\theta$ for the microcosm experiment comparing constant light treatment (bottle 30) versus light-decreasing treatment (bottle 08) (both bottles were inoculated with the same community composition of 7 species). **a, b**. Population densities. **c, d**. Optimum RMSE as a function of window size (m). **e, f**. Time series of optimum $\theta$ that corresponds to the chosen window size (which is marked by the transparent gray rectangle).

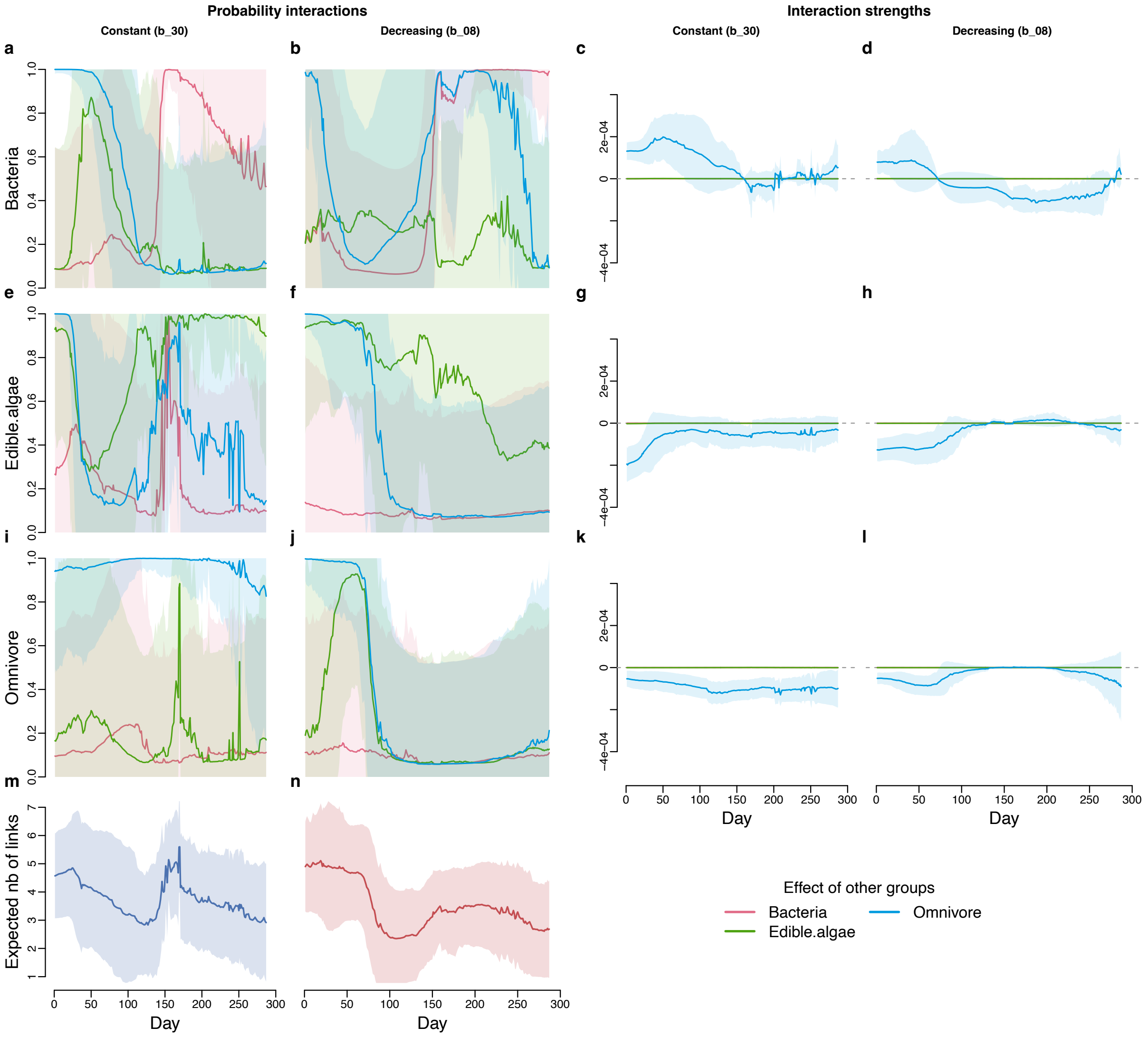

**Supplementary Fig S51.** Inferred interaction probabilities and interaction strengths using the chosen window size for the microcosm experiment comparing constant light treatment (bottle 30) versus light-decreasing treatment (bottle 08) (both bottles were inoculated with the same community composition of 7 species). **a–d**. Effects of other groups on bacteria. **e–h**. Effects of other groups on edible algae. **i–l**. Effects of other groups on omnivore. **m**. Expected number of links in the bottle with constant light treatment. **n**. Expected number of links in the bottle with light-decreasing treatment.

#### S5.6 Mesocosm data

##### S5.6.1 Population densities and nutrient concentration

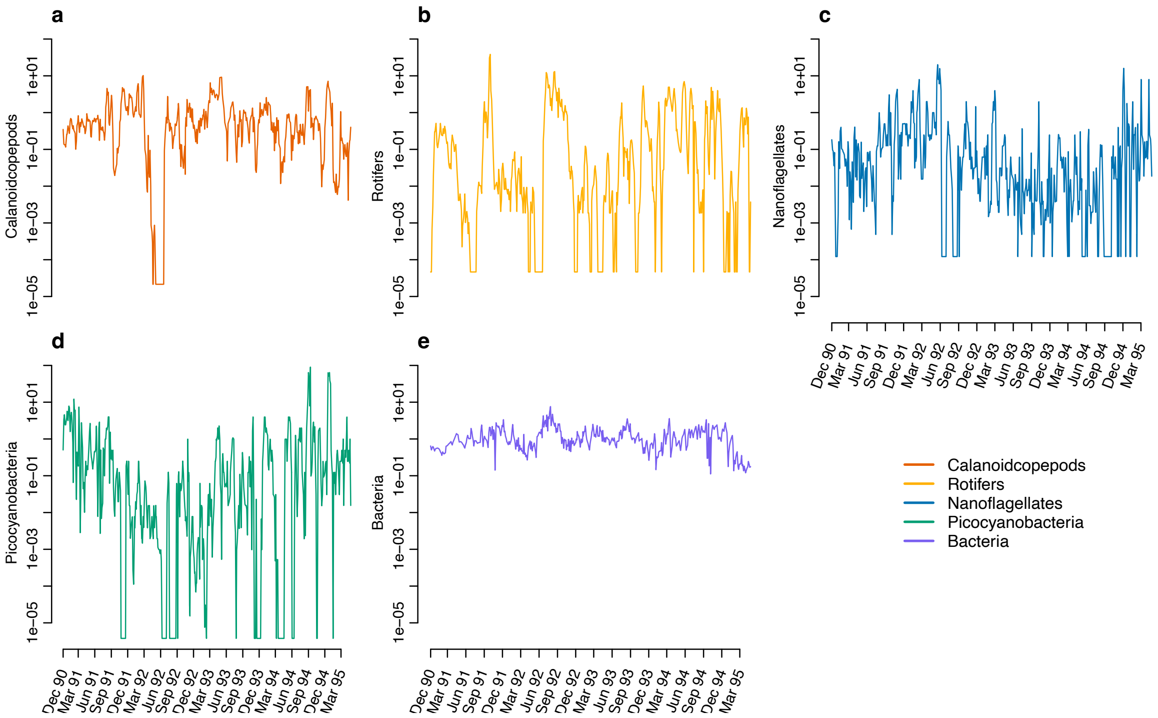

**Supplementary Fig S52.** Population density of the functional groups from the mesocosm experiment, including calanoid copepods, rotifers, and nanoflagellates.

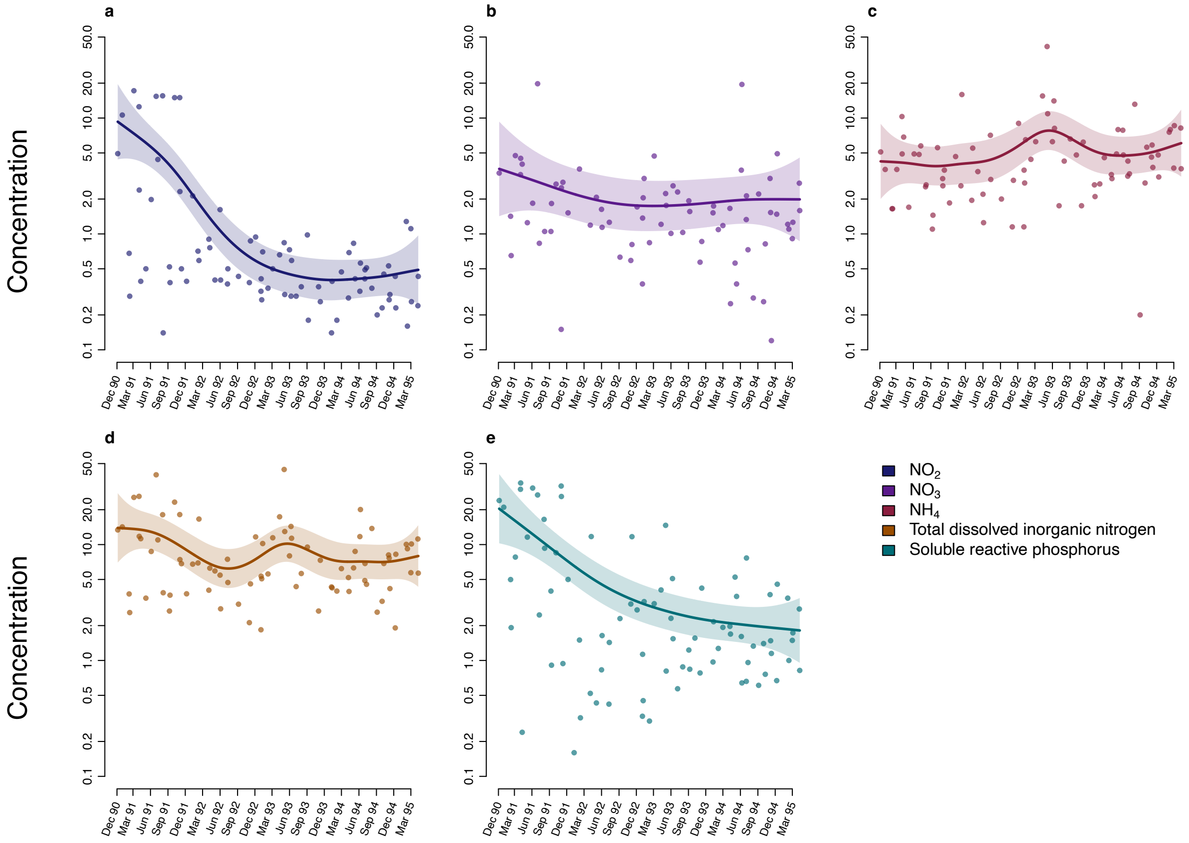

**Supplementary Fig S53.** Nutrient concentration from the mesocosm experiment. The points indicate real data values. The lines indicate the fitted values using the generalised additive mixed model. The transparent ribbons indicate the 95 % confidence interval.

#### S5.6.2 Window sizes and optimum $\theta$

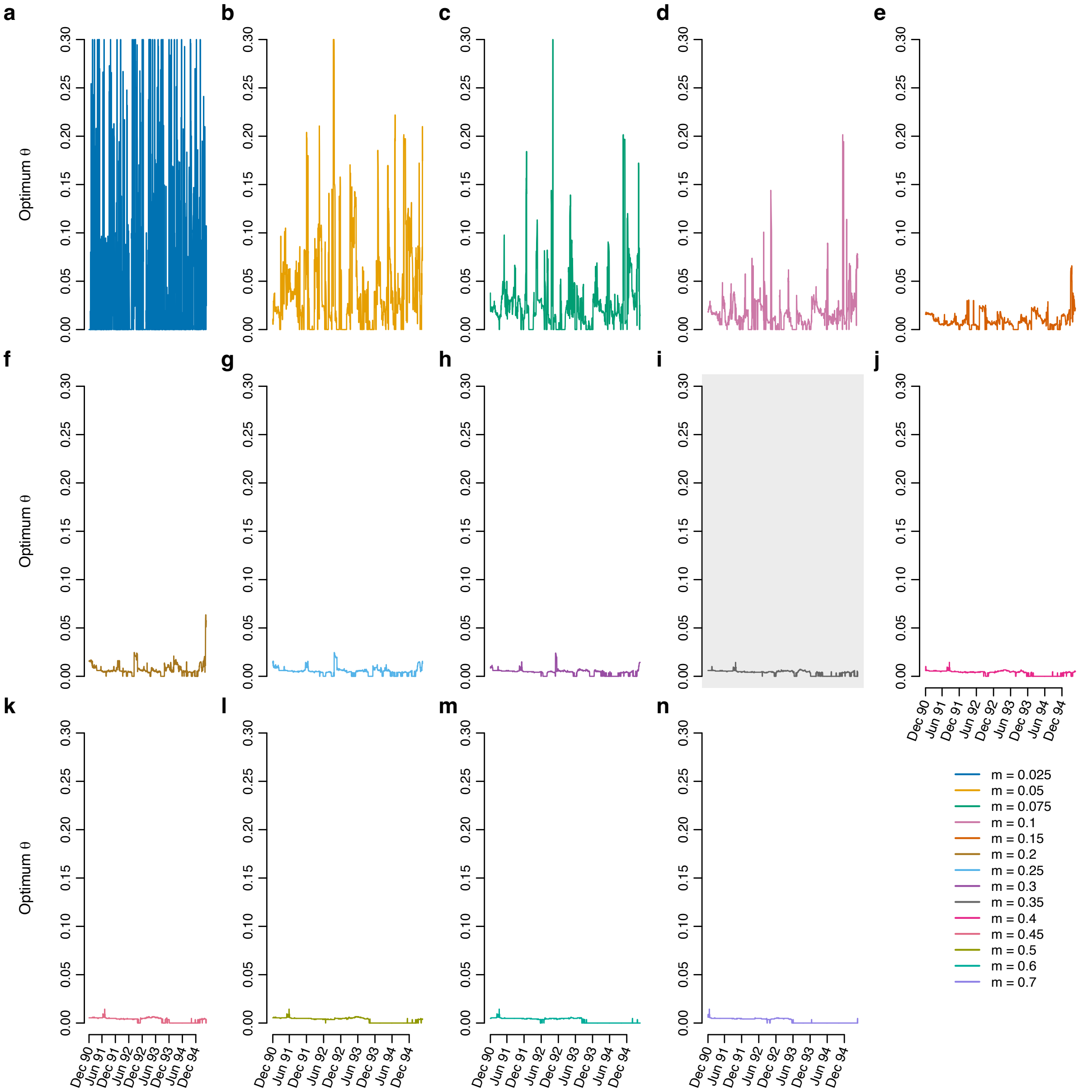

**Supplementary Fig S54.** Time series of the optimum localization values $\theta$ with respect to different window sizes ($m$).

##### S5.6.3 Inference of probability of interactions and interaction strength – BIC

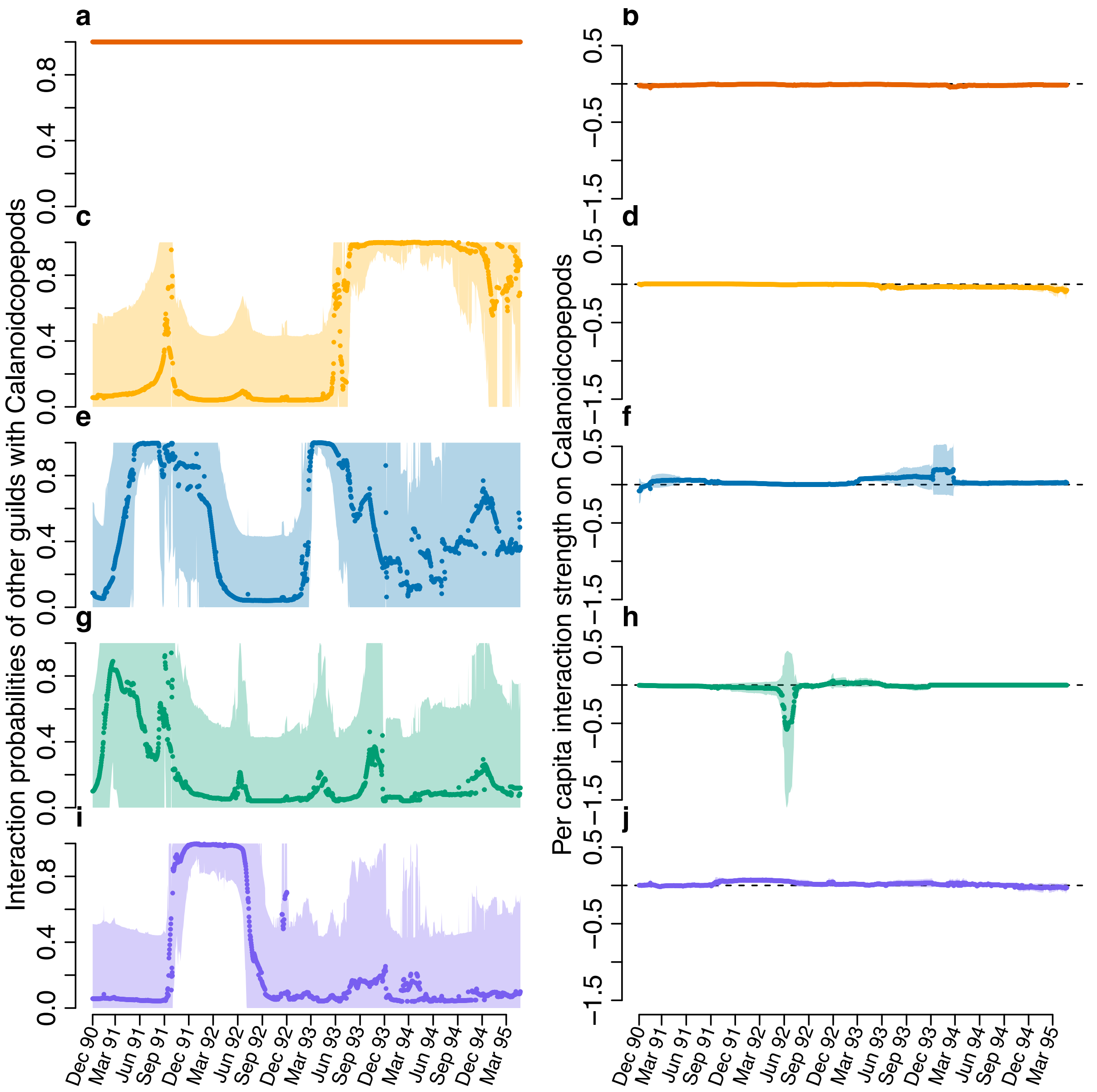

**Supplementary Fig S55.** Inference of probabilities of interactions and interaction strengths for the effects of other groups on calanoid copepods.

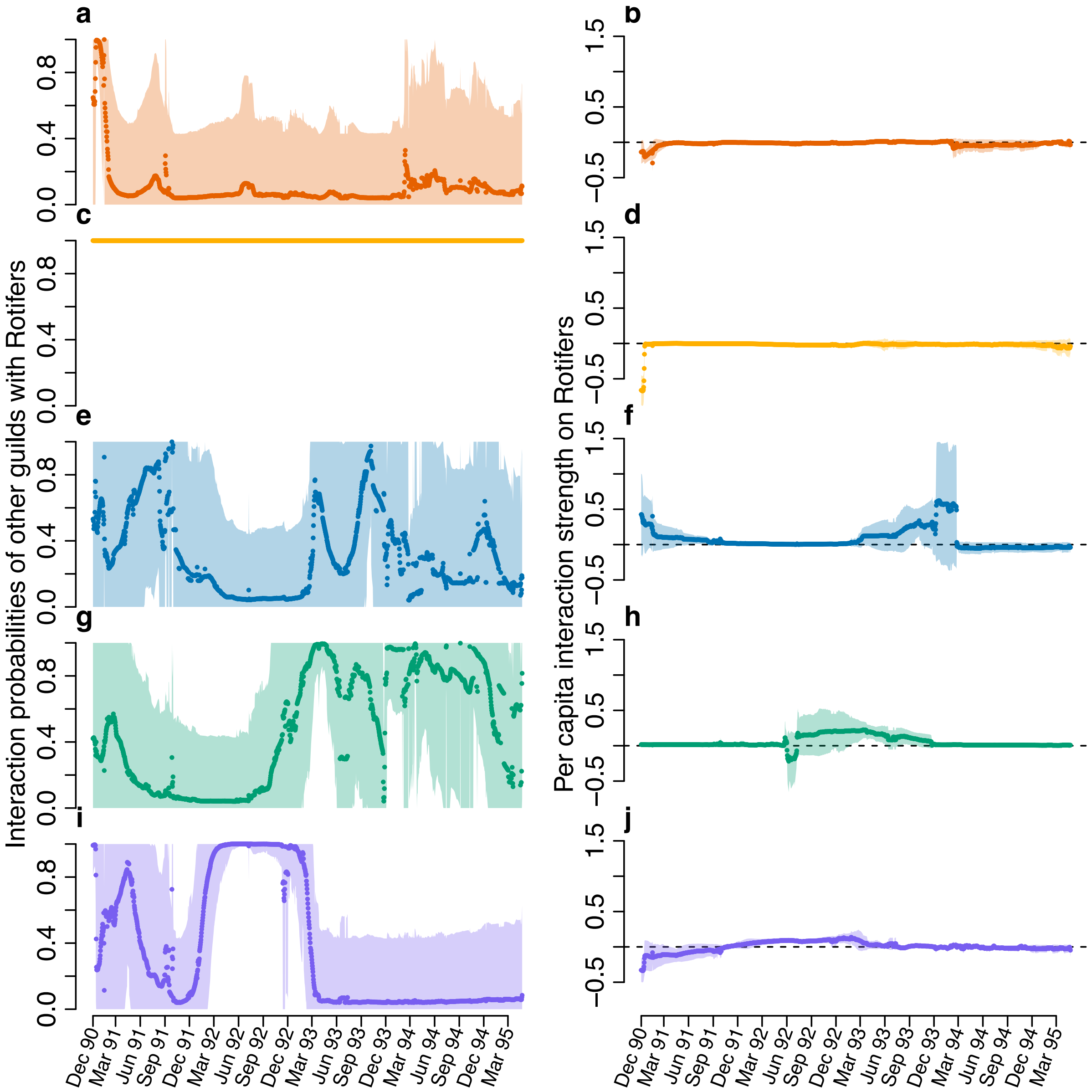

**Supplementary Fig S56.** Inference of probabilities of interactions and interaction strengths for the effects of other groups on rotifers.

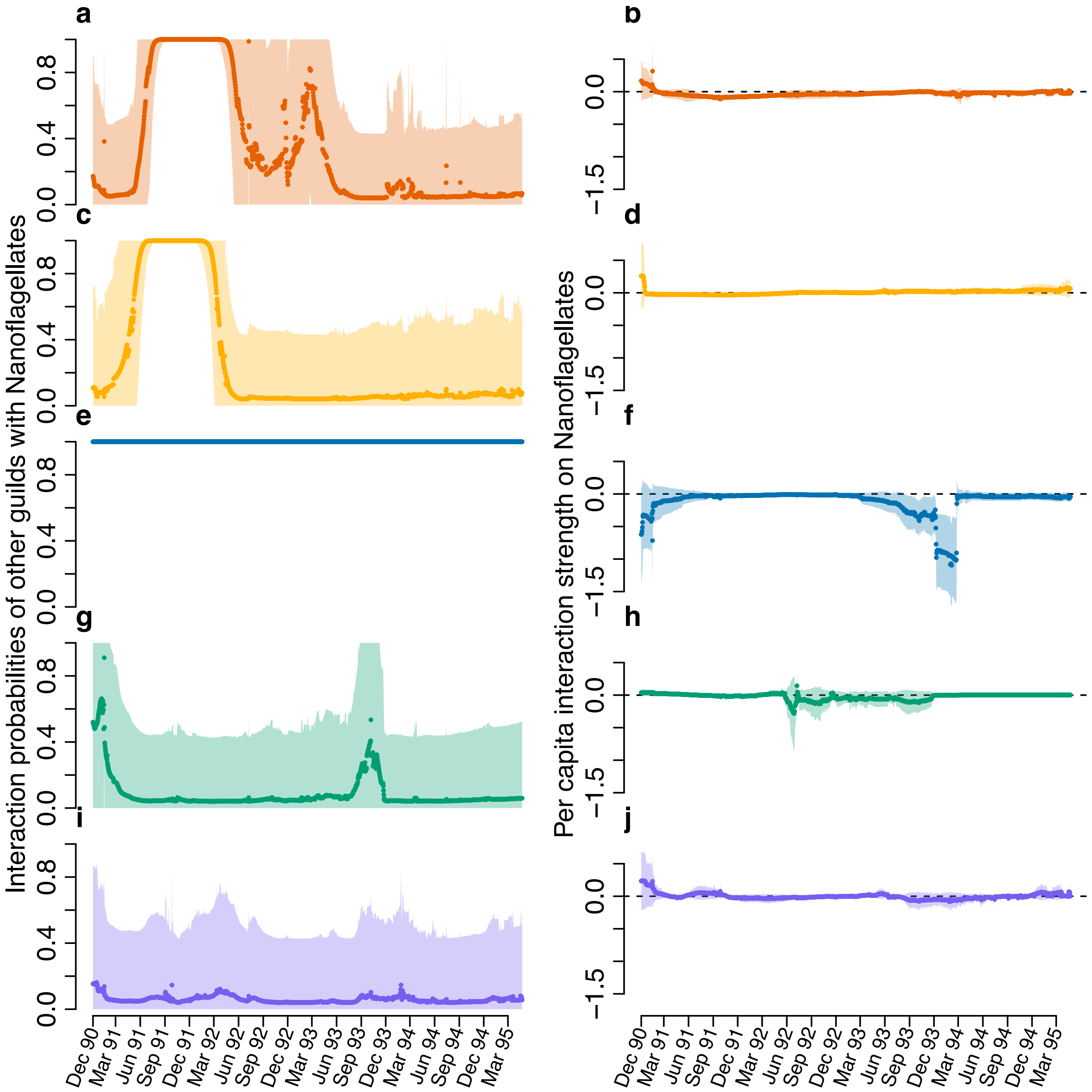

**Supplementary Fig S57. Inference of probabilities of interactions and interaction strengths of other groups on nanoflagellates.**

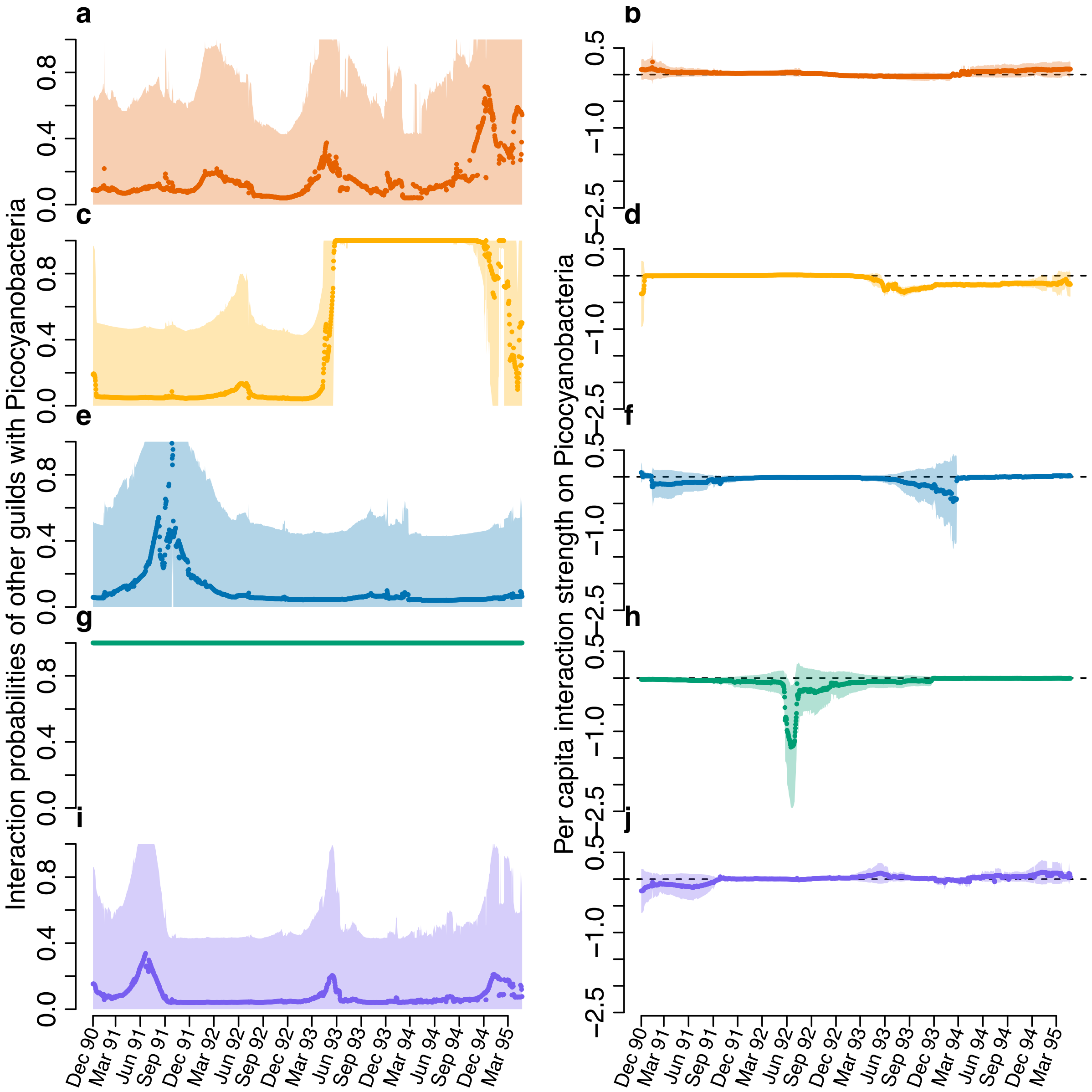

**Supplementary Fig S58. Inference of probabilities of interactions and interaction strengths of other groups on picocyanobacteria.**

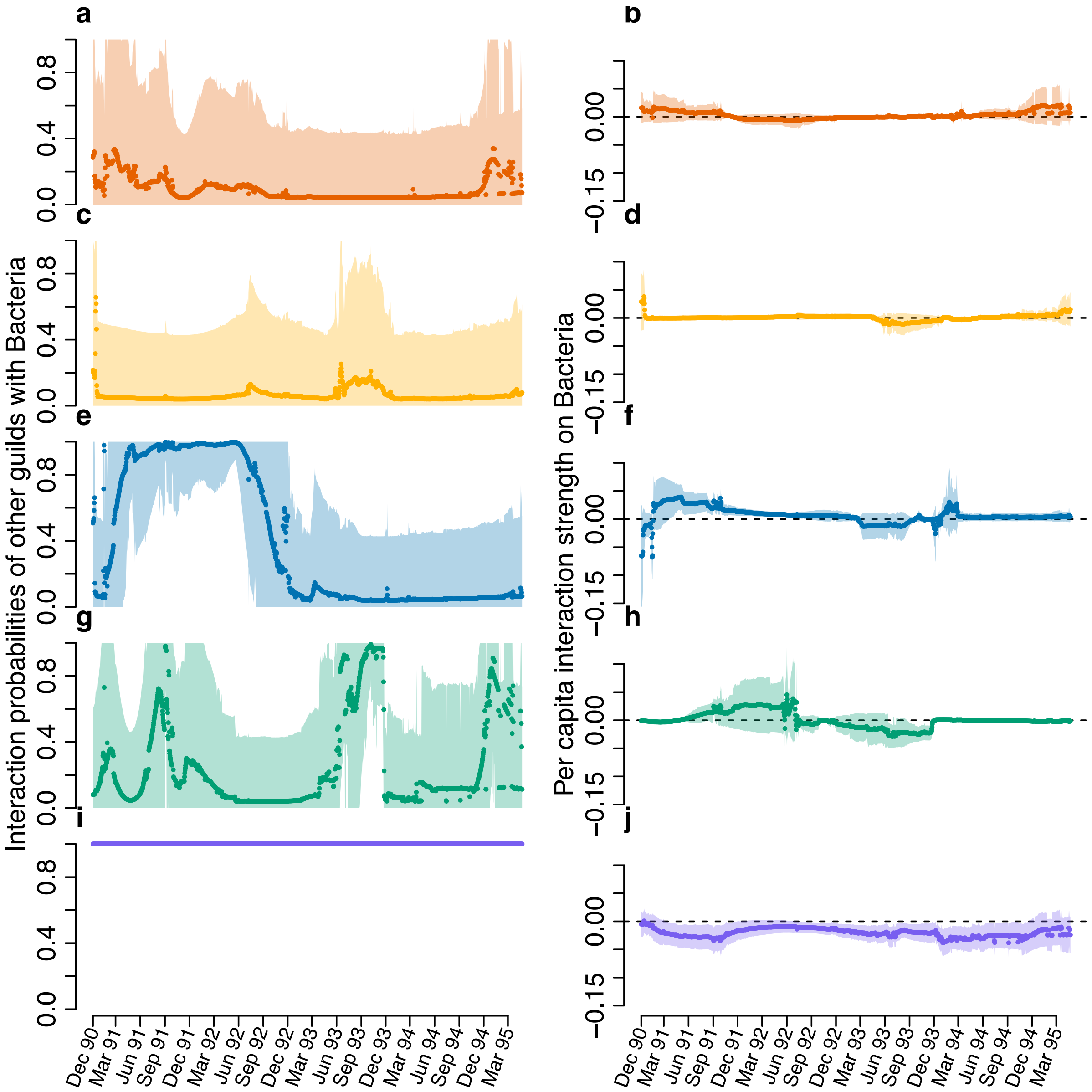

**Supplementary Fig S59. Inference of probabilities of interactions and interaction strengths of other groups on bacteria.**

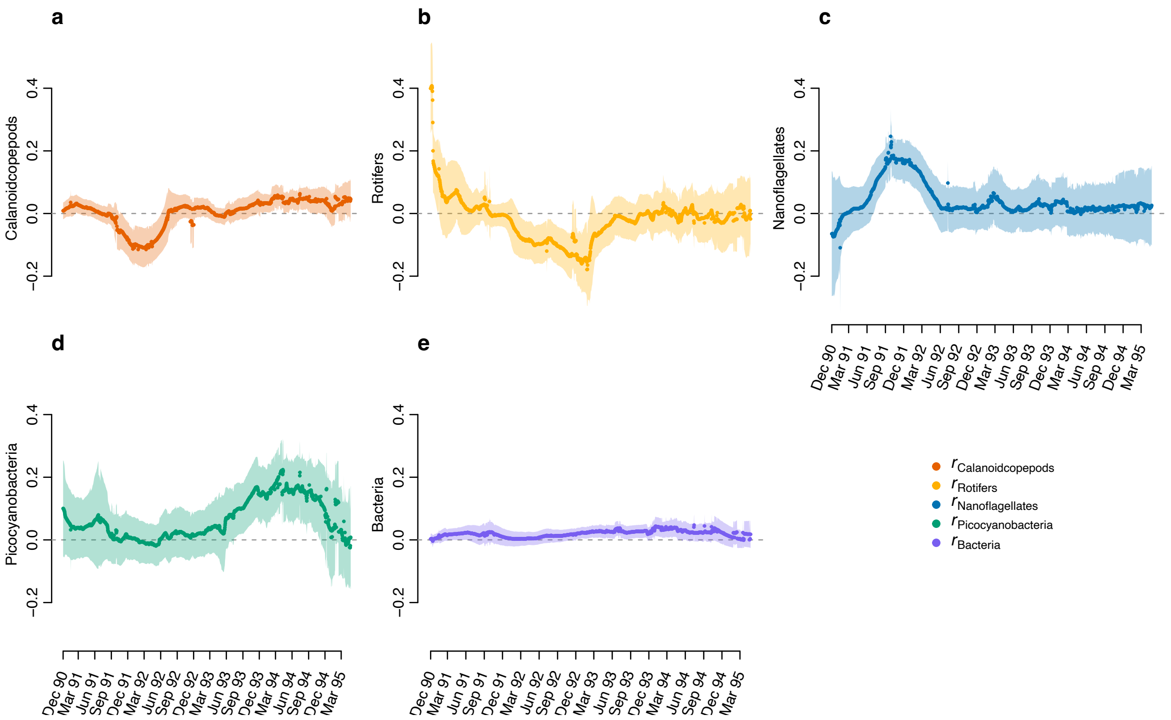

**Supplementary Fig S60. Inference of intrinsic growth rates for calanoid copepods, rotifers, nanoflagellates, picocyanobacteria and bacteria.**

##### S5.6.4 Inferred network using AIC

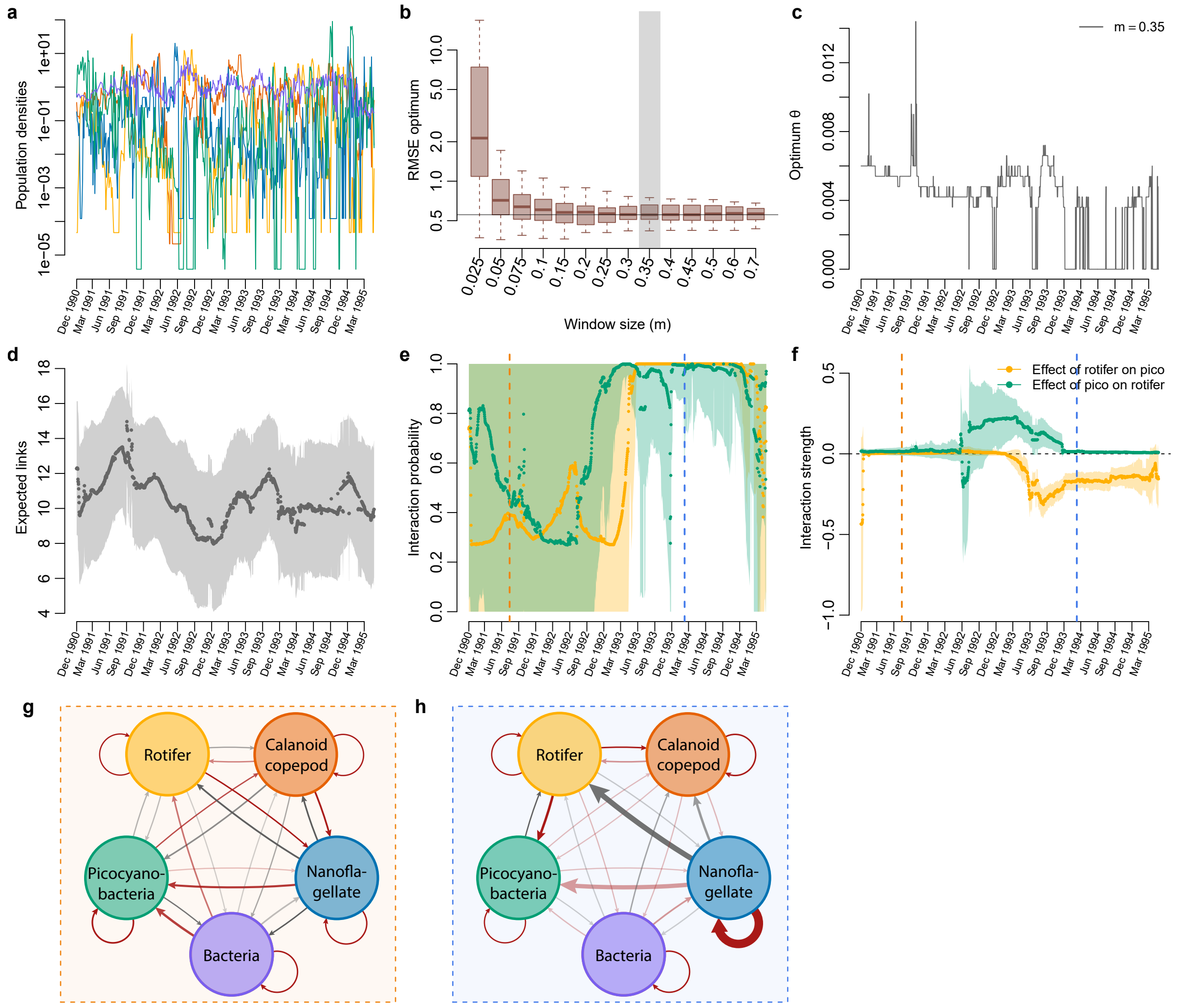

**Supplementary Fig S61. Inference of the mesocosm experiment using the AIC weight.** **(a)** Population densities. **(b)** Optimum RMSE with respect to different window sizes. **(c)** Time series of optimum $\theta$. **(d)** Inferred expected number of links across time. **(e)** Inferred interaction probabilities between rotifer and picocyanobacteria. **(f)** Inferred *per capita* interaction strengths between rotifer and picocyanobacteria. **(g)** The inferred community network on 14 July 1991 (dashed vertical orange line). **(h)** The inferred community network on 8 February 1994 (dashed vertical blue line). Link thickness indicates interaction strength. Link color indicates whether the interaction is positive (gray) or negative (red), and link transparency indicates interaction probability. This figure is the AIC-weight counterpart of figure 5 in the main text.

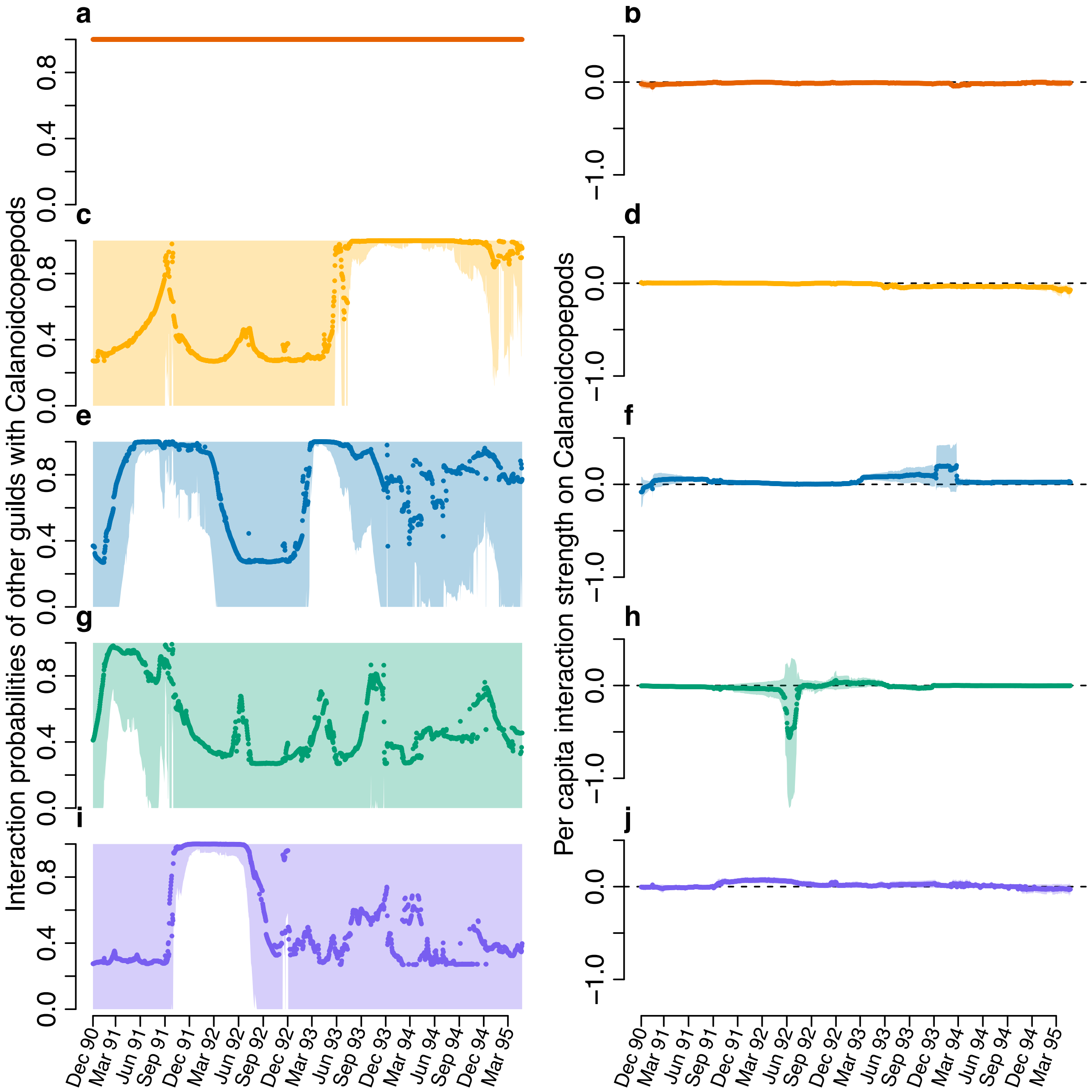

**Supplementary Fig S62. Inference of probabilities of interactions and interaction strengths of other groups on calanoid copepods.**

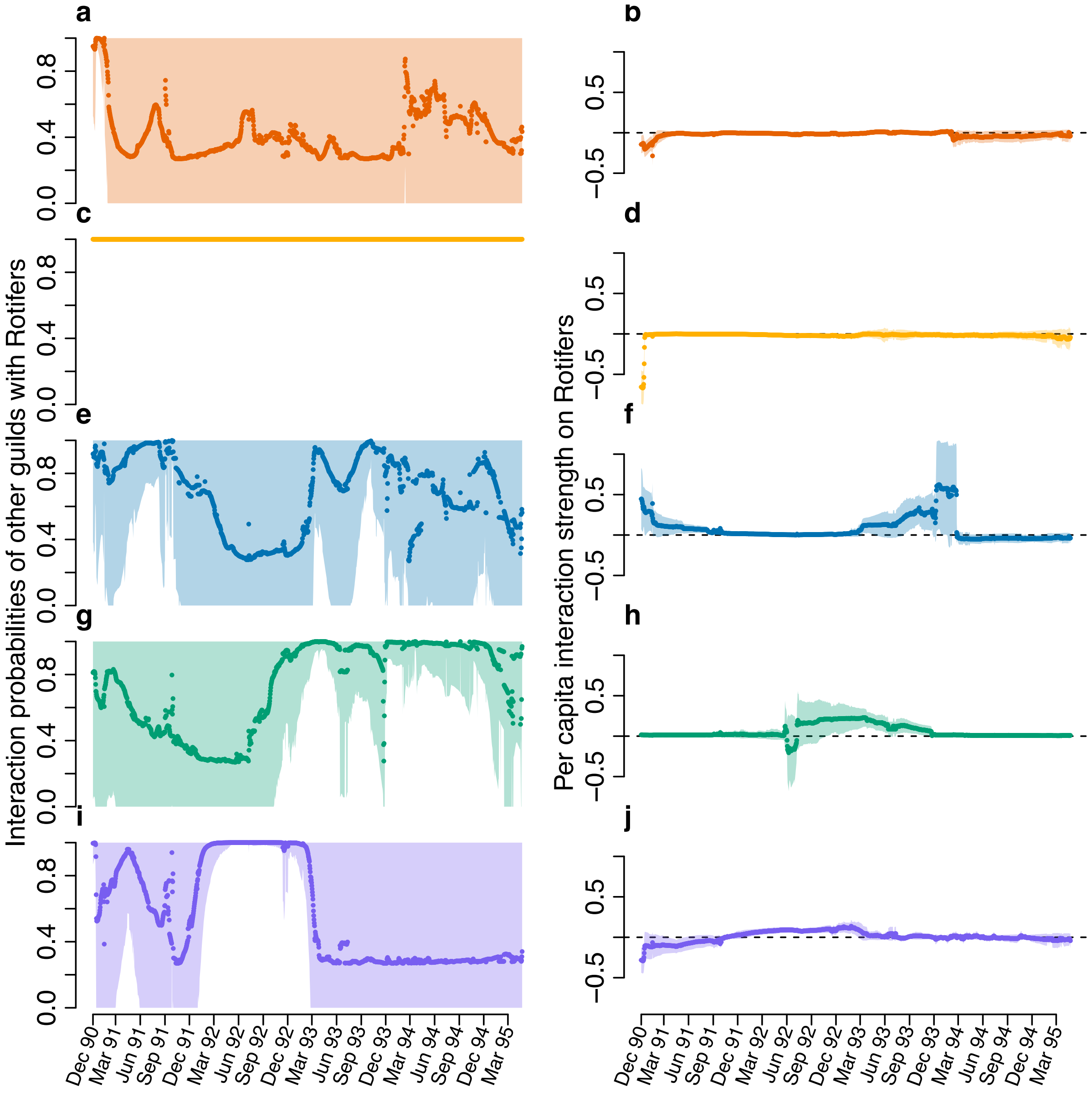

**Supplementary Fig S63. Inference of probabilities of interactions and interaction strengths of other groups on rotifers.**

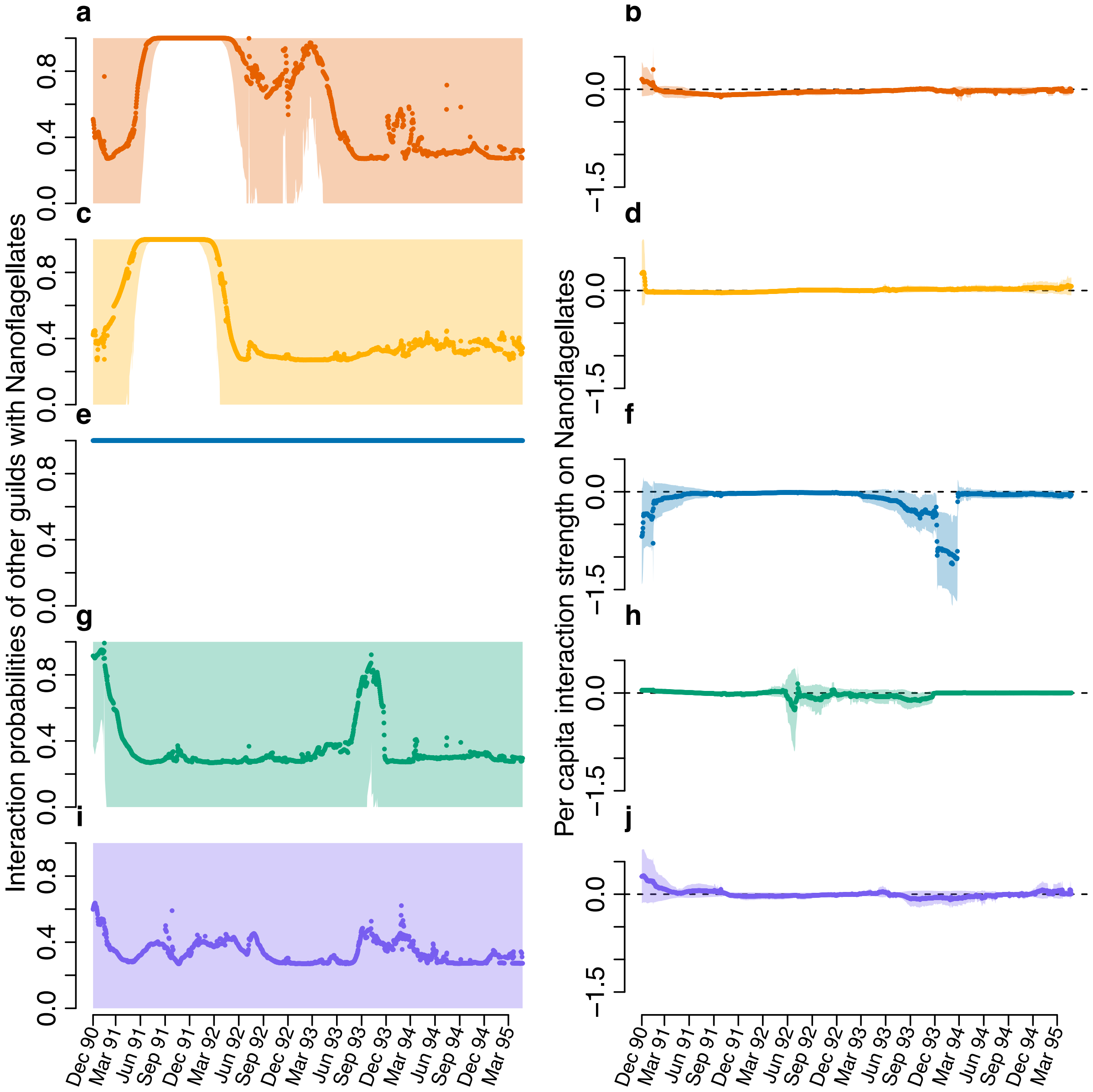

**Supplementary Fig S64. Inference of probabilities of interactions and interaction strengths of other groups on nanoflagellates.**

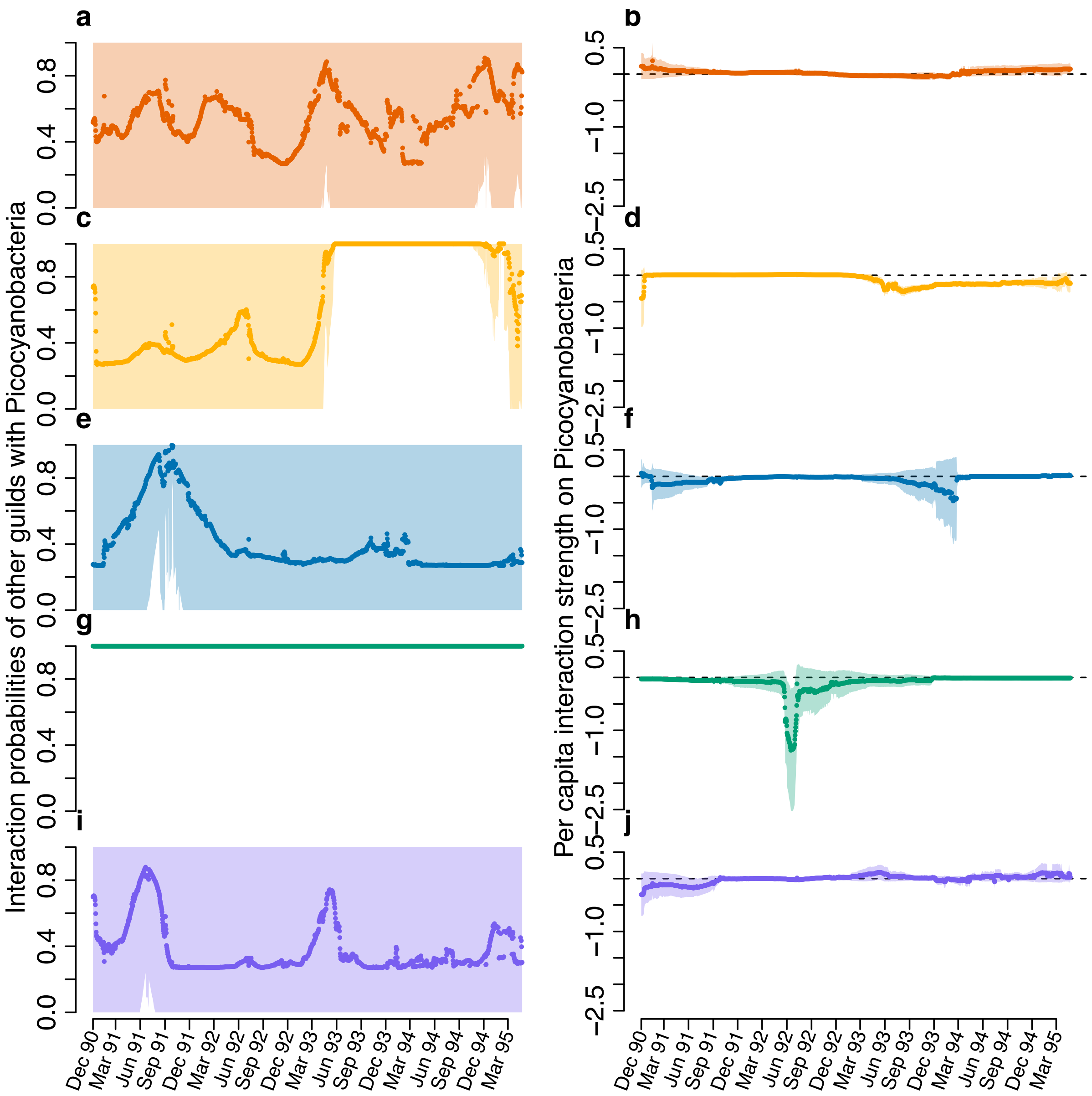

**Supplementary Fig S65. Inference of probabilities of interactions and interaction strengths of other groups on picocyanobacteria.**

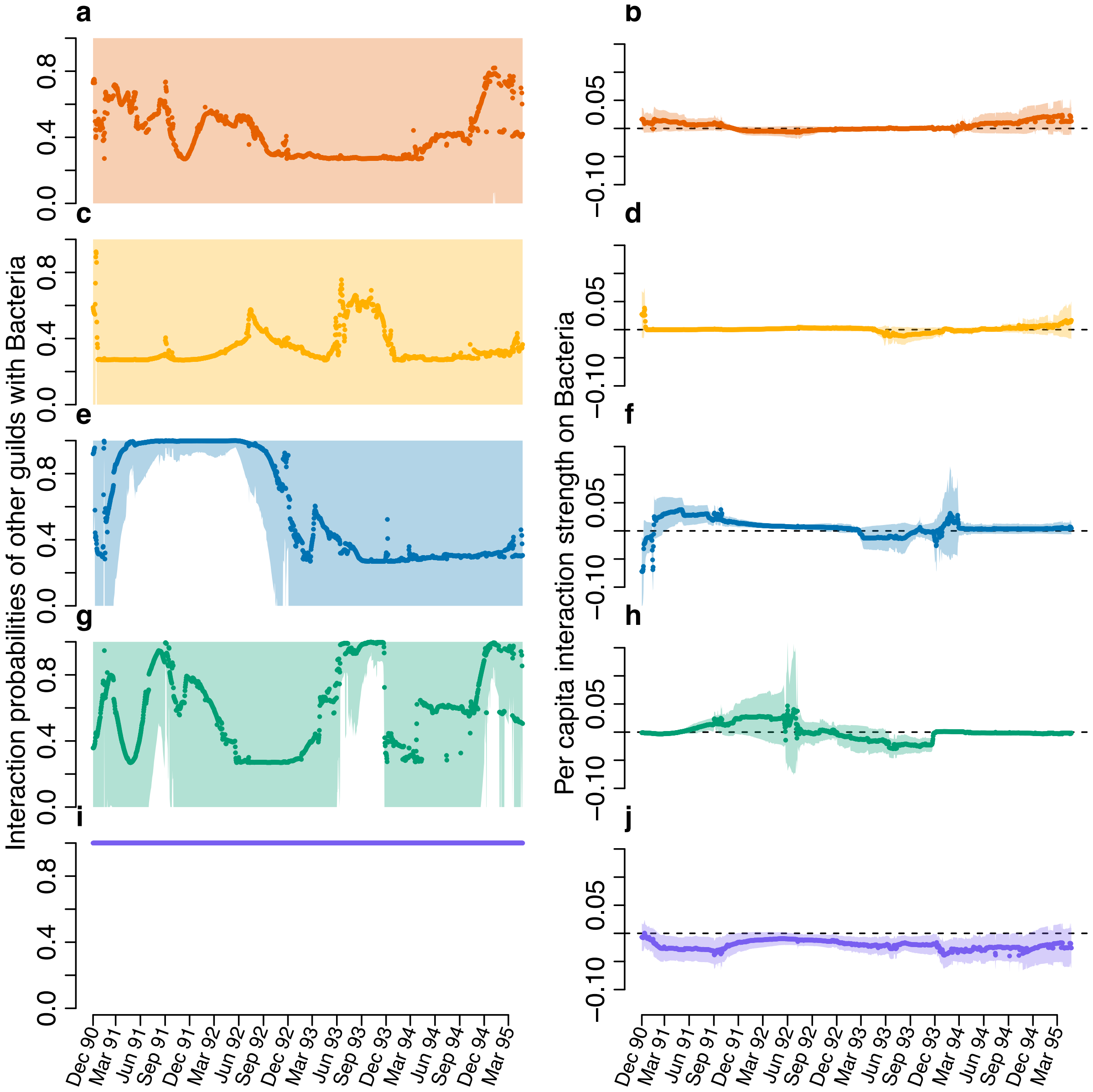

**Supplementary Fig S66. Inference of probabilities of interactions and interaction strengths of other groups on bacteria.**

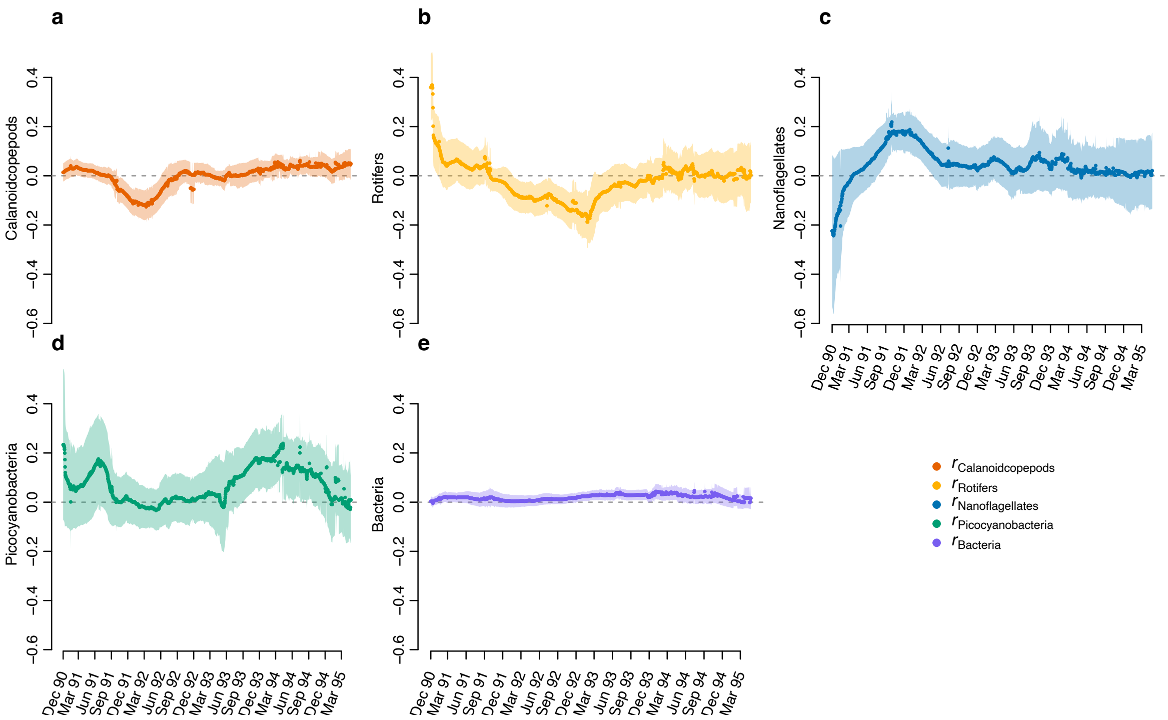

**Supplementary Fig S67. Inference of intrinsic growth rates of calanoid copepods, rotifers, nanoflagellates, picocyanobacteria and bacteria.**
